# Sustained non-photochemical quenching and regulation of PSII repair cycle during combined low temperature and high light stress in lettuce

**DOI:** 10.1101/2024.11.21.624684

**Authors:** Tapio Lempiäinen, Dorota Muth-Pawlak, Julia P. Vainonen, Eevi Rintamäki, Mikko Tikkanen, Eva-Mari Aro

**Author notes:** Current address: Organismal and Evolutionary Biology Research Programme, Faculty of Biological and Environmental Sciences, University of Helsinki, Helsinki, Finland (Julia P. Vainonen).

## Abstract

In nature, light and other environmental conditions are constantly changing, requiring plants to have several overlapping regulatory mechanisms to keep light reactions and metabolism in balance. Here, we show that high light (HL) induces a much stronger down-regulation of light reactions when lettuce plants are exposed to 1500 µmol photons m^−2^ s^−1^ for 4 h at 13°C (low temperature, LT) compared to 23°C (growth temperature, GT). GT/HL treatment induced non-photochemical quenching (NPQ), which relaxed during 1 h recovery in darkness. In contrast, LT/HL treatment induced an exceptionally high NPQ that only partially relaxed during 1 h in darkness at GT. Such a high sustained NPQ (sNPQ) cannot be explained by canonical NPQ mechanism(s). Instead, sNPQ was associated with partial disassembly of PSII-LHCII complexes and a transient increase in phosphorylation of the minor antenna proteins LHCB4.1/LHCB4.2. This coincided with increased expression of the light-harvesting-like proteins SEP2 and ELIP1.2, the PSII assembly proteins HCF173 and LPA3, and accumulation of the pre-D1 protein. These results lead us to propose that LHCB4.1/LHCB4.2 phosphorylation- dependent disassembly of PSII-LHCII supercomplexes allows SEP2 to bind to CP47 and hypothetically quenches the inner PSII core antenna, while free CP43 released during PSII repair is proposed to be protected by LPA3.

## 1. Introduction

Photosynthesis converts light energy into a chemical form that is used to assimilate carbon dioxide into carbohydrates. Two light-driven reactions, catalysed by photosystem II (PSII) and photosystem I (PSI), together with the cytochrome b6f complex, form the major thylakoid protein complexes involved in linear electron transfer (LET) and the coupled proton transport to the thylakoid lumen. LET ultimately reduces ferredoxin (Fd) at the acceptor side of PSI by electrons extracted from water by PSII. Reduced Fd is used by ferredoxin NADP^+^ reductase to form NADPH, while ATP synthase uses the proton gradient to produce ATP from ADP and Pi. NADPH and ATP generated by light reactions are used in stromal metabolism, where carbon assimilation in the Calvin-Benson-Bassham (CBB) cycle is the major sink.

Both photosystems have inner and external light-harvesting antenna that collect excitation energy and transfer it to the reaction centre (RC) chlorophylls (Chl) to ensure their function even in low light conditions. The PSII RC proteins D1 and D2 bind only small amounts of Chl, and two inner or core antenna proteins, CP43 and CP47, contain most of the Chl pigments bound in the PSII core complex (Su et al., 2017). In angiosperms, the PSII inner antenna is associated with an external antenna system composed of three minor LHCII antenna, LHCB4, LHCB5 and LHCB6, and several major LHCII antenna composed of the trimers of LHCB1, LHCB2 and LHCB3, forming LHCII-PSII supercomplexes (sc). The major LHCII antenna is referred to as strongly, moderately and loosely bound trimers (S-LHCII, M-LHCII and L-LHCII, respectively) according to their binding strength to the PSII core complex (Caffarri et al., 2009). In PSI, the inner antenna pigments are bound to the same proteins, PSAA and PSAB, that form the RC complex. The external LHCA antenna belt of PSI is composed of four monomeric LHCA proteins (Mazor et al., 2015). In addition, PSI can receive excitation energy from the LHCII antenna lake shared with PSII, either by forming specific phosphorylation-dependent LHCII-PSI sc or through the LHCA belt (Grieco et al., 2015; Schiphorst et al., 2021).

While the efficiency of light harvesting and the transfer of excitation energy to RCs are largely independent of temperature, even a modest decrease in temperature slows all enzymatic reactions, including the CBB cycle, and thus reduces the strength of electron sinks in the stroma (Hüner et al., 2022). Therefore, the exposure of plants to low temperatures, even under moderate light conditions, requires active mechanisms to dissipate excess energy. The combination of decreasing temperature and increasing light intensity is particularly challenging for the photosynthetic machinery and easily leads to the accumulation of electrons in the LET, increasing the likelihood of harmful side reactions and the generation of reactive oxygen species (ROS). In order to balance light reactions and metabolism to avoid potential ROS-induced hazards under over-excitation conditions, plants must increase the capacity of stromal metabolism by up-regulating the CBB cycle and alternative electron sinks, or dissipate the excess energy in the light-harvesting antenna system or via RC quenching (Schöner and Krause, 1990; Hüner et al., 2016; Herrmann et al., 2019; Bag et al., 2023; Grebe et al., 2024).

The excess excitation energy collected by the LHCII antenna is dissipated by several partially overlapping non-photochemical quenching (NPQ) mechanisms (Bassi and Dall’Osto, 2021; Ruban and Saccon, 2022). The fastest mechanisms in angiosperms involve the energy- and zeaxanthin-dependent components of NPQ (qE and qZ, respectively), which are activated by lumen acidification (Holzwarth et al., 2009; Nilkens et al., 2010; Niyogi and Truong, 2013). In qE, the low lumen pH rapidly induces the protonation of the PSBS protein, which then alters the function of the major LHCII antenna system, leading to the dissipation of excess energy as heat. Activation of the qZ begins shortly after induction of qE, and often both NPQ mechanisms (qE and qZ) continue in parallel under excess light conditions. The xanthophyll cycle catalyses the reversible interconversion of violaxanthin via antheraxanthin to zeaxanthin. Violaxanthin promotes light harvesting while its de-epoxidation to zeaxanthin promotes NPQ.

Apart from the canonical NPQ mechanisms discussed above, several types of sustained NPQ (sNPQ) mechanisms, characterised by a delayed relaxation of NPQ as compared to that of qE, have evolved in different species. The mechanisms of sNPQ are poorly understood, but, particularly in overwintering evergreens, they have been reported to be associated with zeaxanthin accumulation and the expression of the light-harvesting-like (LIL) proteins, but their specific function at the molecular level is poorly defined (Demmig-Adams and Adams, 2006; Zarter et al., 2006b; Zarter et al., 2006a; Levin and Schuster, 2023; Ye et al., 2024). More recently, the specific phosphorylation of LHCII proteins and energy spillover from PSII to PSI have been attributed to sNPQ mechanisms in conifers (Bag et al., 2020; Grebe et al., 2020), and at least one type of sNPQ is known to be activated by the lipocalin protein in Arabidopsis (*Arabidopsis thaliana*) LHCII (qH) (Malnoë et al., 2017; Bru et al., 2022).

Despite several NPQ and other protective mechanisms, PSII is known to undergo a continuous cycle of damage and repair under all light conditions (Tyystjärvi and Aro, 1996), and the inhibition of PSII photochemical activity can only be detected when the rate of damage exceeds the rate of repair (Aro et al., 1993). The photoinhibited PSII core complexes with damaged D1 protein (Savitch et al., 2002) are not photochemically active, but it has recently been reported that they are still able to quench the excitation energy non-photochemically (qIRC) (Nawrocki et al., 2021). However, it remains unclear how qIRC and the PSII repair cycle function as part of the network of NPQ mechanisms.

Here, we investigated how a 10°C decrease in temperature affects the induction and relaxation of NPQ in lettuce. A decrease in temperature was found to induce a strong sNPQ that cannot be explained by the previously described NPQ mechanisms. To unravel the processes behind such a strong sNPQ in lettuce and to investigate the underlying processes behind the slow relaxation of sNPQ in the dark at physiological temperature, we used of biochemical, biophysical, and proteomic approaches according to the experimental protocol in Figure 1.

**Figure 1.**
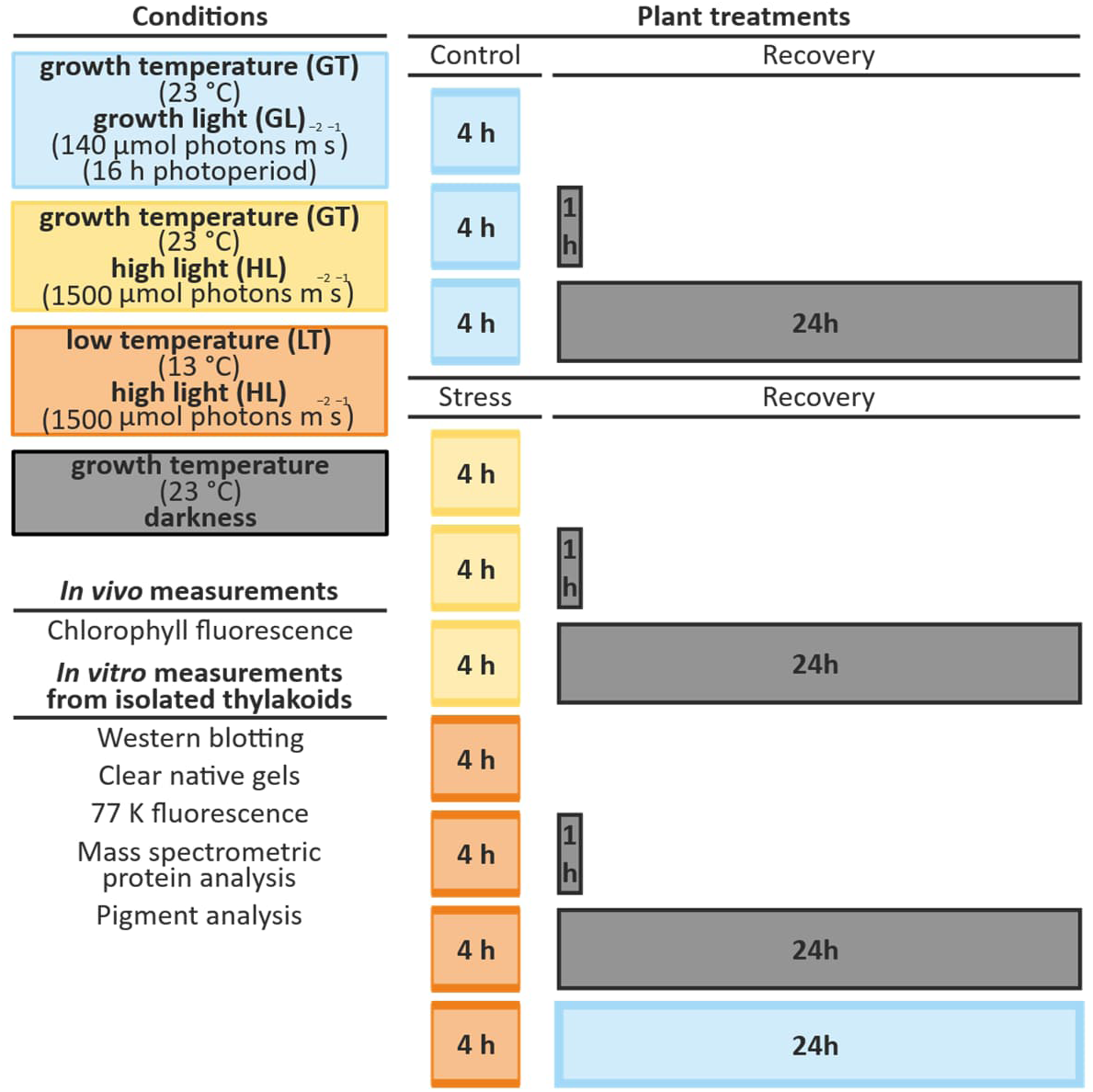
Experimental set-up to study the short-term acclimation mechanisms of the photosynthetic apparatus of lettuce to excessive excitation stress induced by simultaneous high light (HL) and low temperature (LT) stress. Long-day grown lettuce plants were treated at 13 °C under white light of 1500 µmol photons m^−2^ s^−1^ for 4 h (LT/HL) (orange boxes), while control plants were kept at growth temperature and light conditions (23 °C, 140 µmol photons m^−2^ s^−1^) (GT/GL) (light blue boxes), or treated at HL in growth temperature (GT/HL) (yellow boxes), after which all plants were transferred to recover for 1 h and 24 h in darkness (grey boxes) or for 24 h in normal long-day conditions at GT/GL. *In-vivo* Chl fluorescence measurements were performed from all samples. In addition, thylakoids were isolated from LT/HL-treated plants and from GT/GL-control plants directly after the treatment (blue and orange boxes) and subsequent recovery periods of 1 h and 24 h in darkness or growth conditions. Isolated thylakoids were used for biochemical analyses and 77K *in-vitro* Chl fluorescence measurements.

## 2. Materials and methods

### 2.1 Lettuce growth conditions, light treatments and recovery conditions

Hilde White Boston cultivar of lettuce (*Lactuca sativa*) was grown in a 16 h photoperiod in moderate white light (140 µmol photons m^−2^ s^−1^) with POWERSTAR HQI-T 400W/D metal halide lamps (OSRAM GmbH) as the light source, at growth temperature, 23 °C (GT). The plants used in the experiments were 4 weeks old.

Plants were treated (Figure 1) with high light (HL, 1500 µmol photons m^−2^ s^−1^) for 4 h with Heliospectra Dyna LED lamps at low temperature (LT, 13 °C) of groth (orange boxes). After the light treatment, one part of the plants was transferred to recover in darkness at GT for 1 h, and for 24 h (grey boxes), and the other part was transferred to recover in growth conditions (GT/GL) for 24 h (light blue boxes). Control plants, without the LT/HL-treatment, were transferred to similar recovery conditions. In addition, lettuce plants were also exposed to a sole HL treatment (yellow box) (GT/HL) and monitored only by Chl fluorescence after the HL treatment.

### 2.2 Chl fluorescence measurements

Chl fluorescence was measured with a PAR-FluorPen FP 110 using the saturating pulse method (Photon Systems Instruments, Drásov, Czech Republic) with default settings. The quantum yield of PSII photochemistry (Y(II)) was estimated as (FM’-F0’)/FM’ in light- and as (FM-F0)/FM in dark-acclimated plants. NPQ was calculated as (FM^ref^/FM’)-1 in light- and as (FM^ref^/FM)-1 in dark-acclimated plants using as the reference value (FM^ref^), the average of FM from control plants transferred to darkness for 24 h. qLT (Porcar-Castell, 2011) was calculated as ((1/ F0’)-(1/ FM’))/((1/ F0^ref^)-(1/ FM^ref^)) using as the reference values (FM^ref^ and F0^ref^) the averages of FM and F0 from control plants shifted to darkness for 24 h.

### 2.3 Thylakoid isolation

Thylakoids were isolated from GT/GL and LT/HL-treated and recovered plants similarly to as described in (Gunell et al., 2023) with minor modifications (see Supplemental file 1). Chl concentration was determined in buffered acetone according to (Porra et al., 1989).

### 2.4 Western blotting

Western blot analysis was performed as described in (Gunell et al., 2023) with minor modifications (see Supplemental file 1).

### 2.5 Clear native and 2D gel electrophoresis and phosphoprotein staining

Clear native gel electrophoresis (CN) and 2D gel electrophoresis were performed as described in (Järvi et al., 2011) with minor modifications (see Supplemental file 1). Phosphoprotein staining with ProQ was performed according to the manufacturer’s instructions (Invitrogen), and the gels were imaged on a Perkin Elmer Geliance 1000 with a Cy3 filter.

### 2.6 77 K Chl fluorescence

77 K Chl fluorescence measurements were performed with small thylakoid aliquots (50 µl) diluted to a Chl concentration of 10 µg/ml, to minimise self-absorption. Thylakoids were excited with 480 nm light in liquid nitrogen and fluorescence was detected using Ocean Optics S2000 spectrophotometer. Spectra were normalised to 685 nm peak and the ratio between the height of the 735 nm peak to 685 nm peak (F735/F685) was calculated to illustrate the distribution of excitation energy between PSII and PSI.

### 2.7 Proteomics analysis

Proteins from isolated thylakoids were solubilised, digested, and desalted according to a previously described protocol (Huokko et al., 2019), with minor modifications (see Supplemental file 1). The nLC-ESI-FAIMS-MS/MS analyses were performed on a nanoflow HPLC system (Easy-nLC 2000, Thermo Fisher Scientific) coupled to an Orbitrap Fusion Lumos mass spectrometer (Thermo Fisher Scientific) equipped with a high-field asymmetric waveform ion mobility spectrometry (FAIMS Pro^TM^) interface in data-independent acquisition (DIA) mode (see Supplemental file 1).

Data analysis was performed using Spectronaut (version 16.1) software (Bruderer et al., 2015) (Biognosys, Schlieren, Switzerland). DIA data were searched against a customised FASTA file containing 37 834 protein sequences (Supplemental file 5) using the Pulsar directDIA^TM^ algorithm. The customized FASTA file contained the manually curated lettuce proteomes from UniProt (retrieved 2023.02.15) and Swiss-Prot (retrieved 2023.03.28) (Supplemental file 2). UniProt homologs of Swiss-Prot sequences were manually removed from the combined database based on BLAST searches (Supplemental files 3 and 4). Peptide identification was performed with trypsin as an enzyme allowing a maximum of 2 missed cleavages, as well as carbamidomethylation set as static modification, while methionine oxidation, N-terminal acetylation, and lysine acetylation in addition to serine, threonine, and tyrosine phosphorylation set as dynamic modifications. The FDR identification threshold (q-value) for peptides and proteins was set at 0.01. Data for target proteins (identified based on homology to Arabidopsis sequences (Supplemental files 6 and 7)) were extracted and analysed in Excel files. Protein abundances were normalised within samples, to the average of the detected PSI, ATP synthase, and Cyt b6f subunits to avoid, as much as possible, the effect of contaminants from the isolations and differential attachment of stromal proteins to the thylakoids under changing experimental conditions (e.g. a switch to HL rapidly recruits ribosomes to the thylakoid membrane), similarly to previous studies on the plant thylakoid proteome (Flannery et al., 2021b; Flannery et al., 2021a). The original data and protein quantification project (Spectronaut file) are deposited in PRIDE Archive database (Vizcaíno et al., 2016) (PXD055190).

### 2.9 Pigment analysis

Pigments were extracted and separated according to (Gilmore and Yamamoto, 1991), with minor modifications (see Supplemental file 1). Separated pigments were identified by their absorbance spectra and relative retention times. Relative pigment abundances were estimated by the area of chromatographic peaks detected at 440 nm and normalised within the sample to the area of Chl a.

## 3. Results

### 3.1 Temperature reduction from 23°C to 13°C during HL treatment of lettuce induces strong sNPQ

Lettuce plants were exposed to high light at low temperature (LT/HL) and at growth temperature (GT/HL) for 4 h to investigate how photosynthetic acclimation differs when metabolism can operate at optimal temperature compared to limiting metabolism by lowering temperature by 10 °C. Chl fluorescence analysis revealed a drastic decrease in PSII quantum yield (Y(II)) from 0.78 in GT/GL controls (Figure 2A) to 0.08 during the 4 h LT/HL treatment (Figure 2A), but only a modest decrease to 0.45 during the 4 h GT/HL treatment (Figure 2E). After 1 h recovery in the dark at 23 °C, Y(II) in GT/HL-treated lettuce recovered to 0.65, but only to 0.10 in LT/HL-treated lettuce (Figures 2E and 2A). After 24 h recovery in darkness at 23 °C, Y(II) was fully recovered in GT/HL-treated lettuce, but only to 0.49 in the case of LT/HL-treated lettuce (Figures 2E and 2A), suggesting that part of the reduction in Y(II) was due to PSII inhibition, which did not recover in darkness. This interpretation was supported by the fact that Y(II) of LT/HL-treated lettuce recovered to 0.75 within 24 h under GT/GL (Figure 2A), where the PSII repair cycle is active.

**Figure 2.**
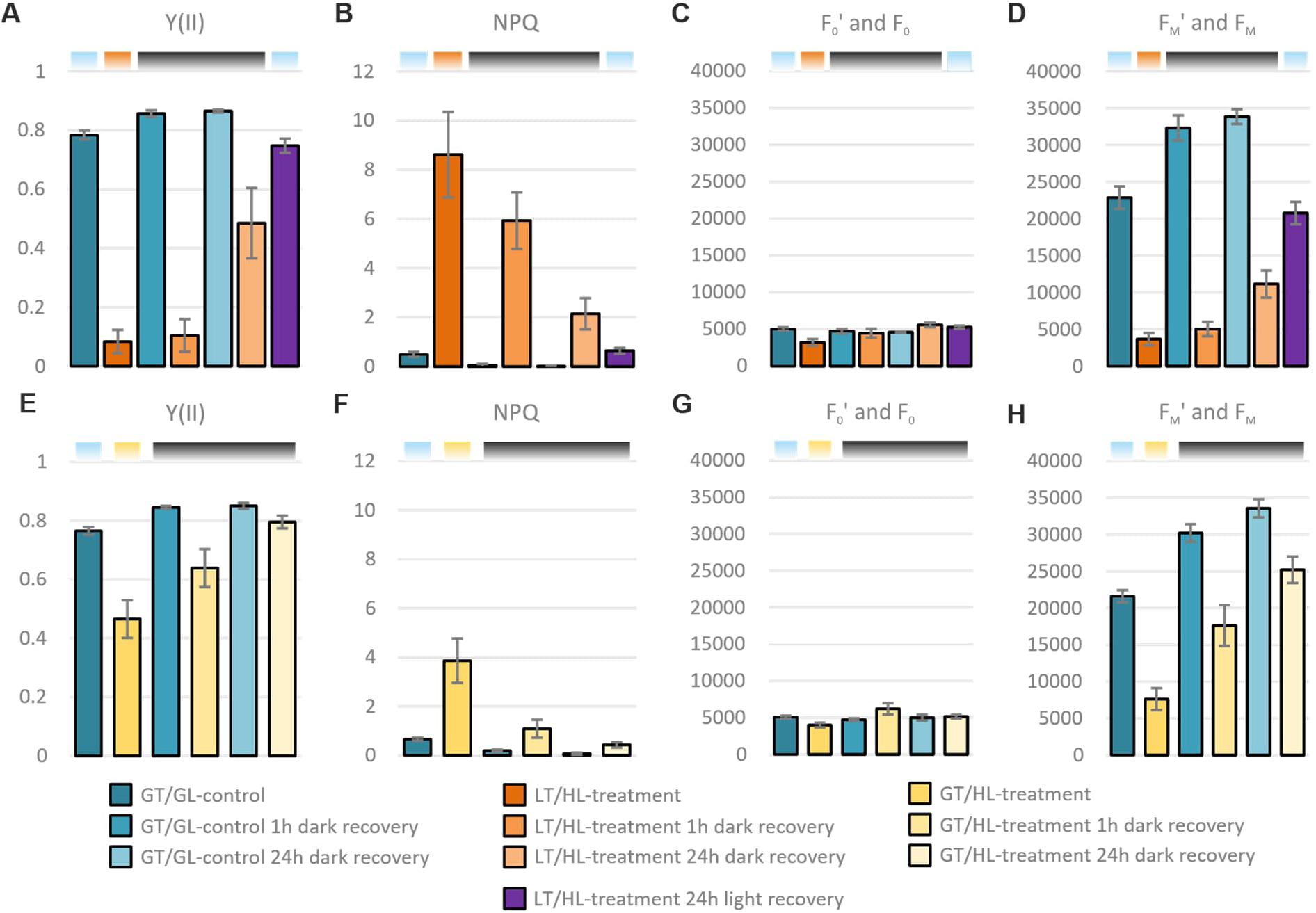
Comparison of low temperature and high light treatment (LT/HL) with high light treatment alone (GT/HL) on the formation of sustained excitation energy quenching in lettuce. **A)** and **E)** PSII quantum yield (Y(II)). **B)** and **F)** Non-photochemical quenching (NPQ). **C)** and **G)** Dark fluorescence yields (F0’ and F0, arbitrary units). **D)** and **H)** Maximum and maximal fluorescence yields (FM’ and FM, arbitrary units). Long day-grown lettuce plants were illuminated under 1500 µmol photons m^−2^ s^−1^ of white light at 13 °C for 4 h (LT/HL) or at 23 °C (GT/HL). Control plants were kept at growth conditions (23 °C, 140 µmol photons m^−2^ s^−1^) (GT/GL), after which all plants were transferred to recover at 23 °C for 1 h and 24 h in the dark or for 24 h in the long day growth conditions. F0’, F0, FM’, and FM were determined with Fluorpen directly after the treatment and after subsequent recovery periods of 1 h and 24 h. Y(II) was calculated as (FM’-F0’)/FM’ in light- and as (FM-F0)/FM in dark-acclimated plants. NPQ was calculated as (FM^ref^/FM’)-1 in light- and as (FM^ref^/FM)-1 in dark-acclimated plants using as the reference value (FM^ref^) the average of FM from control plants shifted to darkness for 24 h. Error bars show standard deviations among biological replicates (n = 4-16). Coloured bars above the graphs represent the light conditions in which the measurement was conducted: light blue for growth light, orange and yellow for high light, and black for darkness.

We then calculated the NPQ parameter in the control and treated lettuce (GT/HL and LT/HL, respectively) (Figures 2B and 2F), using the average FM of dark recovered controls (GT/GL 24 h dark recovery) as a reference value. NPQ was normal at 3.86 in GT/HL-treated lettuce and very high at 8.16 in LT/HL-treated lettuce immediately after the treatments (Figures 2F and 2B). In GT/HL-treated lettuce, the NPQ was mostly relaxed after 1 h and 24 h recovery in darkness (Figure 2F). In contrast, the high NPQ in LT/HL-treated lettuce relaxed slowly and was 5.93 after 1 h and 2.09 after 24 h recovery in darkness (Figure 2B). When raw fluorescence yields (F0’ and F0) were also considered, both GT/HL and LT/HL treated lettuce showed a decrease in F0’ proportional to NPQ (Figures 2C and 2G), in agreement with previous reports (Rees et al., 1992). After 1 h of dark recovery, such a clear difference between LT/HL treated and GT/GL control lettuce had disappeared, and after 24 h of dark recovery, the F0 value was higher in LT/HL treated than in the GT/GL control, indicating persistent PSII photoinhibition and complete relaxation of NPQ. Maximum and maximal fluorescence yields (FM’ and FM, respectively) (Figures 2D and 2H) showed similar dynamics to Y(II), suggesting that the changes in Y(II) are mainly due to changes in FM’ and FM with only a minor contribution from changes in F0’ and F0.

Since the comparison of fluorescence parameters between the LT/HL-treated (Figures 2A-D) and GT/HL-treated (Figures 2 E-H) lettuce revealed typical NPQ in GT/HL-treated lettuce but a high sNPQ in LT/HL-treated lettuce, we will from now on focus on elucidating the sNPQ mechanism(s) induced by the LT/HL treatment and compare the results with GT/GL controls.

### 3.2 LT/HL treatment and recovery alter thylakoid protein phosphorylation and composition of pigment- protein complexes

The phosphorylation of the LHCII proteins has previously been linked to sNPQ (Grebe et al., 2020), which prompted us to analyse the phosphorylation of thylakoid proteins with the p-Thr antibody from the GT/GL control, LT/HL treated and recovered lettuce plants (Figure 3A). The LT/HL treatment increased the phosphorylation level of the PSII core proteins CP43, D2 and D1, but completely abolished the phosphorylation of LHCB1 and LHCB2 (Figure 3A). LHCII phosphorylation returned during the 1 h dark recovery, whereas the prolonged 24 h dark recovery led to almost complete dephosphorylation of the D1, D2 and LHCII proteins in both GT/GL control and LT/HL treated lettuce. In turn, the recovery in growth conditions restored thylakoid protein phosphorylation to the level of GT/GL control plants. Strikingly, we also detected strong phosphorylation of the minor antenna protein LHCB4 in the LT/HL treated lettuce (Figure 3A).

**Figure 3.**
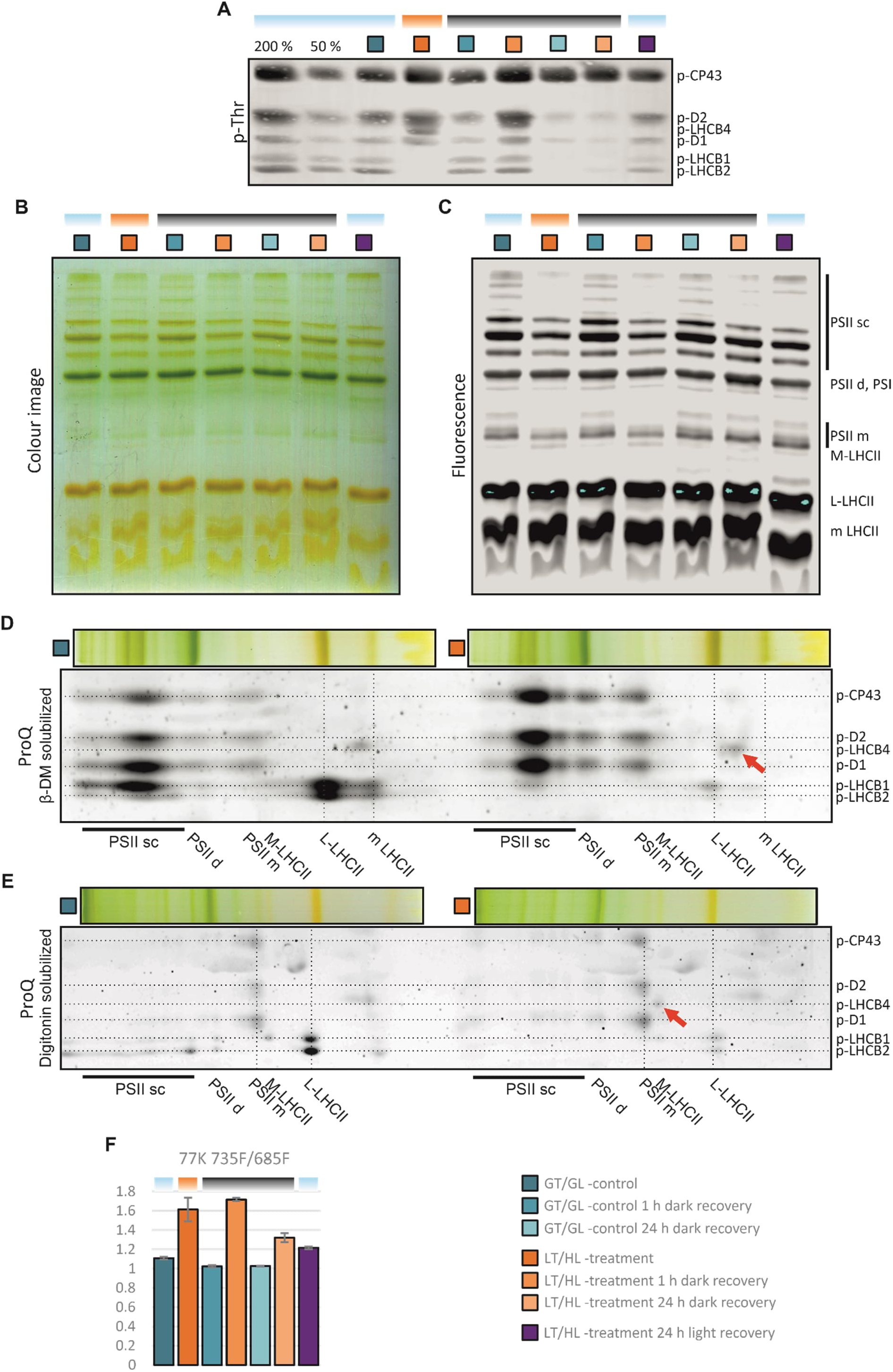
Effect of the low temperature and high light treatment, and the subsequent recovery, on phosphorylation-dependent regulation of light harvesting in lettuce. **A)** Immunodetection of protein phosphorylation. Thylakoid proteins were separated with SDS-PAGE, transferred to the PVDF membrane and immunodetected with p-Thr antibody and IR-dye labelled secondary antibody. **B)** Colour image and **C)** Fluorescence image of β-DM solubilized pigment-protein complexes separated with clear native gel electrophoresis. The fluorescence of separated pigment-protein complexes was visualised by exciting the complexes with a 685 nm laser with Odyssey CLx imager (light blue regions in the L-LHCII indicate oversaturation). **D and E)** Phosphoprotein staining (ProQ) of the 2D gels of thylakoid proteins solubilised with β-DM (D) and digitonin (E) before electrophoresis in the first dimension. LCHB4 is indicated with red arrows. **F)** Fluorescence ratio of PSI (735 nm) to PSII (685 nm) from the 77K Chl fluorescence emission spectra of isolated thylakoids (F735/F685). Error bars show standard deviations among technical replicates (*n* =3). Long day-grown lettuce plants were illuminated under 1500 µmol photons m^−2^ s^−1^ of white light at 13 °C for 4 h (LT/HL), while control plants were kept at growth conditions (23 °C and 140 µmol photons m^−2^ s^−1^ with 16 h photoperiod) (GT/GL), after which all plants were transferred to recover for 1 h and 24 h in darkness or for 24 h in long day growth conditions. Thylakoid membranes used in the analyses were isolated directly after the treatment and the recovery periods. Coloured bars above the graphs represent the temperature and light conditions from which the thylakoids used in the analysis were isolated: light blue for GT/GL, orange for LT/HL and black for GT/darkness. Abbreviations: PSII supercomplexes (PSII sc), PSII dimer (PSII d), Photosystem I (PSI), PSII monomer (PSII m), Moderately bound LHCII trimer (M-LHCII), Loosely bound LHCII trimer (L-LHCII) and monomeric LHCII (m LHC).

Next, we analysed the effects of the LT/HL treatment and recovery on the organisation of the pigment-binding thylakoid protein complexes, using a CN gel electrophoresis (Figure 3B). After electrophoresis, the fluorescence of the separated complexes was analysed to reveal the possible involvement of qH (Bru et al., 2022), as a component of the strong sNPQ, but no major changes in the fluorescence of L-LHCII trimers were detected (Figure 3C). On the other hand, the CN gels showed reduced amounts of PSII sc and increased amounts of M-LHCII in LT/HL-treated lettuce compared to the GT/GL control (Figure 3B). Detached M-LHCII was not fully reassociated with the PSII core during dark recovery, as the amount of PSII sc remained at a lower level than in GT/GL controls, which could be caused by degradation of damaged PSII cores or degradation of M-LHCII. These results were consistent with the fluorescence analysis of the gels (Figure 3C), which also showed a reduction in the fluorescence of monomeric PSII complexes in thylakoids isolated from LT/HL-treated lettuce, which was also visible after 1 h of dark recovery.

To confirm the identity of the LT/HL treatment induced phosphoprotein as p-LHCB4 (Figure 3A), we separated the thylakoid proteins by 2D gel electrophoresis, and visualised them with a phosphoprotein specific stain (Figure 3D). The position of the p-LHCB4 in the 2D gels was affected by the choice of detergent in the native gel electrophoresis used for separation in the first dimension. With β-dodecyl maltoside (β-DM) as a solubilising agent, the phosphoprotein was found to migrate between L-LHCII trimers and monomeric antenna proteins (Figure 3D, red arrow), whereas with digitonin it migrated in the M-LHCII complex (Figure 3E, red arrow). M-LHCII is composed of the LHCB1/LHCB3 trimer together with the LHCB4 and LHCB6 proteins, confirming the identification of the additional phosphoprotein as p-LHCB4, the only component of M-LHCII with the same mobility in CN. This suggested that the association of p-LHCB4 with M-LHCII is relatively weak, as it is detached from the complex with β-DM (Figure 3D, red arrow), which is a slightly stronger detergent than digitonin. Furthermore, p-LHCB4 was not detected in PSII sc (Figures 3D and 3E), further suggesting that LHCB4 phosphorylation detaches M-LHCII from the PSII core.

Further information on changes in the relative antenna sizes of PSI and PSII during LT/HL treatment and subsequent recovery was obtained from the 77K Chl fluorescence emission spectra recorded from isolated thylakoids. The LT/HL treatment increased the fluorescence emission ratio of PSI to PSII (F735/F685) from 1.11 in GT/GL control to 1.61 (Figure 3F), which is consistent with the detachment of M-LHCII from PSII sc, thereby reducing the relative excitation of PSII. Recovery in darkness for 1 h further increased the fluorescence ratio to 1.72 compared to 1.02 of the GT/GL control, suggesting that the increased phosphorylation of LHCB2, which is associated with the LHCII-PSI complex formation, has an additional effect on top of the detachment of M-LHCII from PSII sc. After 24 h recovery in the dark, the LT/HL treated samples still showed a higher F735/F685 ratio 1.31 compared to that of the GT/GL controls 1.03. This could be due to the still reduced amount of PSII sc in the 24 h recovered plants compared to the GT/GL-control (Figure 3B). The increased F735/F685 ratio after LT/HL treatment persisted not only after the 24 h dark recovery but also after recovery in light compared to the GT/GL control (Figure 3B).

### 3.3 Thylakoid protein level changes during LT/HL treatment and subsequent recovery compared to GT/GL controls

To gain a more comprehensive insight into the mechanisms involved in the quenching of excitation energy in LT/HL-treated lettuce and during the subsequent sNPQ relaxation, we took a targeted proteomic approach to gain a broader view on how the proteins of PSII core, light harvesting system, and repair cycle were affected by the experiment. To this end, proteins from isolated thylakoids were detergent-solubilised, digested with trypsin, and the resulting peptides were analysed by nLC-ESI-FAIMS-MS/MS in DIA mode that provides reliable protein quantification.

#### 3.3.1 LT/HL treatment leads to only minor depletion of the PSII core proteins

PSII photoinhibition damages D1 and D2 proteins and induces qIRC. Therefore, we analysed the levels of major PSII core proteins and minor antenna proteins (Figure 4). The decrease in the levels of the PSII RC proteins D1 and D2 was small, about 15-20%, and occurred during the LT/HL treatment and remained at the reduced level also during the subsequent recovery in darkness (Figures 4A, 4B). Notably, the levels of the inner antenna proteins CP47 and CP43 remained stable during the LT/HL treatment, but a small fraction of them was degraded during recovery in darkness and light (Figure 4C and 4D), significantly later than for the PSII RC proteins D1 and D2. The amounts of the minor light-harvesting antenna (LHCB4-6) were slightly variable (Figures 4E, 4F, 4G, 4H, 4I and 4J), but there were no trends like those observed for the PSII core proteins.

**Figure 4.**
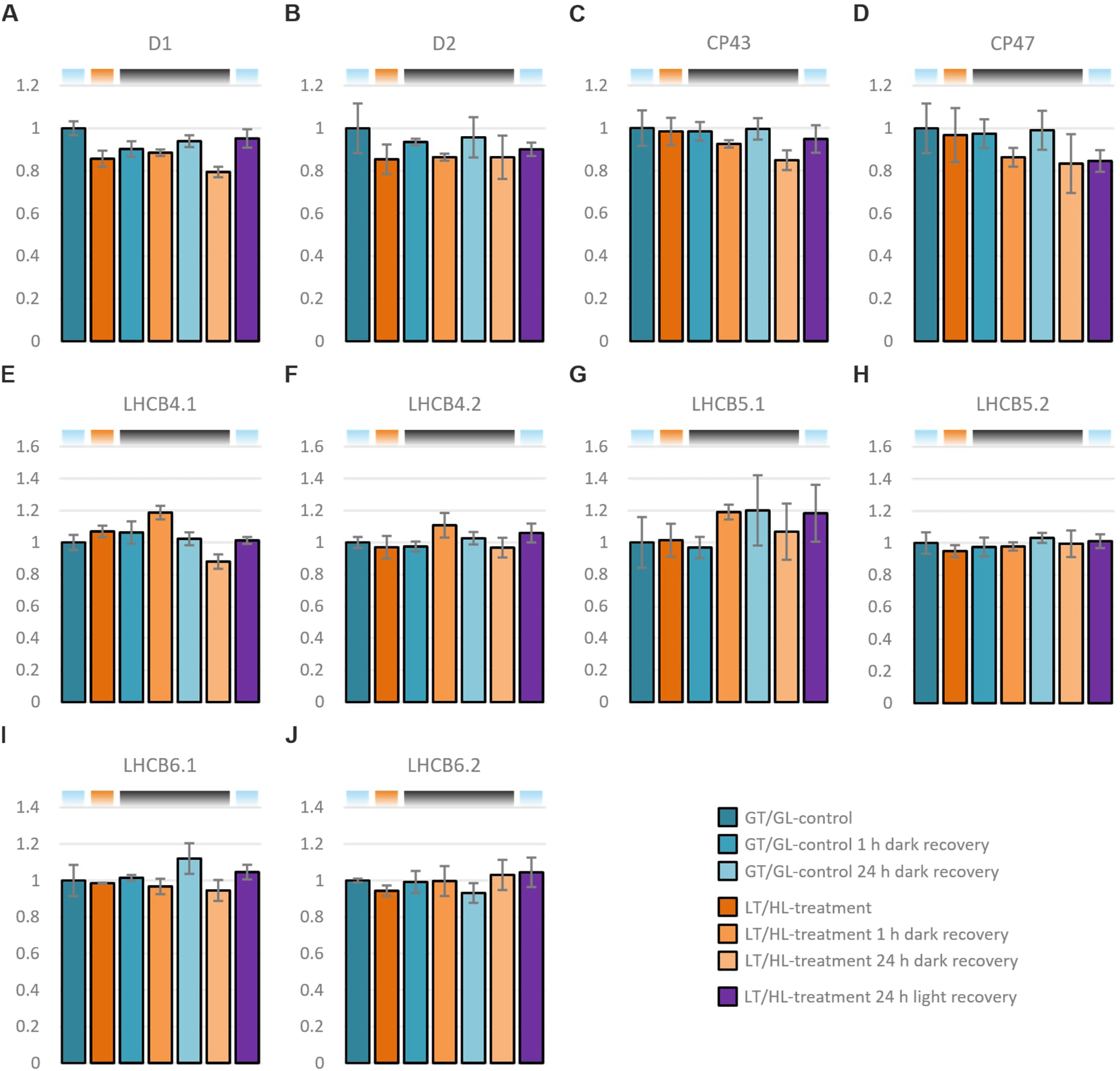
Effect of the low temperature and high light treatment, and the subsequent recovery, on major PSII core complex proteins and minor antenna proteins. **A)** D1 **B)** D2 **C)** CP43 **D)** CP47 **E)** LHCB4.1 **F)** LHCB4.2 **G)** LHCB5.1 **H)** LHCB5.2 **I)** LHCB6.1 **J)** LHCB6.2. Long day-grown lettuce plants were illuminated under 1500 µmol photons m^−2^ s^−1^ of white light at 13 °C for 4 h (LT/HL), while control plants were kept at growth conditions (23 °C and 140 µmol photons m^−2^ s^−1^ with 16 h photoperiod) (GT/GL), after which all plants were transferred to recover for 1 h and 24 h in darkness or for 24 h in long day growth conditions. Thylakoid membranes used in the analyses were isolated directly after the treatment and the recovery periods. Isolated thylakoids were solubilized with detergent and isolated proteins were digested with trypsin. Resulted peptide mixtures were analysed with nLC-ESI-FAIMS-MS/MS in DIA mode and protein abundances were determined with Spectronaut software. Protein abundances were normalised to the average of control plants in growth light. Error bars show standard deviations among technical replicates (n = 3). Coloured bars above the graphs represent the temperature and light conditions from which the thylakoids used in the analyses were isolated: light blue for GT/GL, orange for LT/HL and black for GT/darkness.

#### 3.3.2 Only a specific subset of PSII repair machinery is upregulated by LT/HL treatment

Since we detected changes in the fluorescence of PSII monomers (Figure 3C), associated with the PSII repair cycle, we focused on proteins known to be involved in PSII repair and biogenesis, such as high chlorophyll fluorescence (HCF) 173, HCF244, albino 3 (ALB3), and low PSII accumulation proteins (LPA1, LPA2, and LPA3) (Figure 5). HCF173 binds psbA transcripts and assists in translation initiation, and the interaction between HCF173 and HCF244 recruits D1-translating polysomes to the thylakoid membrane (Schult et al., 2007; Link et al., 2012; Chotewutmontri and Barkan, 2020; Wang and Grimm, 2021). ALB3 is required for the insertion of newly translated D1 into the CP43-less PSII (Schneider et al., 2014), whereas HCF136 (Ycf48 in cyanobacteria) has a functional role in preventing the premature formation of the oxygen evolving complex (Zhao et al. 2023). In addition, LPA1 has been implicated in the initial stages of D1 insertion and it acts in concert with HCF136, whereas LPA2 and LPA3 have been reported to act in later stages of PSII assembly and repair by binding free CP43 and supporting its rebinding to the RC47 PSII intermediate (Nickelsen & Rengstl 2013; Schneider et al. 2014).

**Figure 5.**
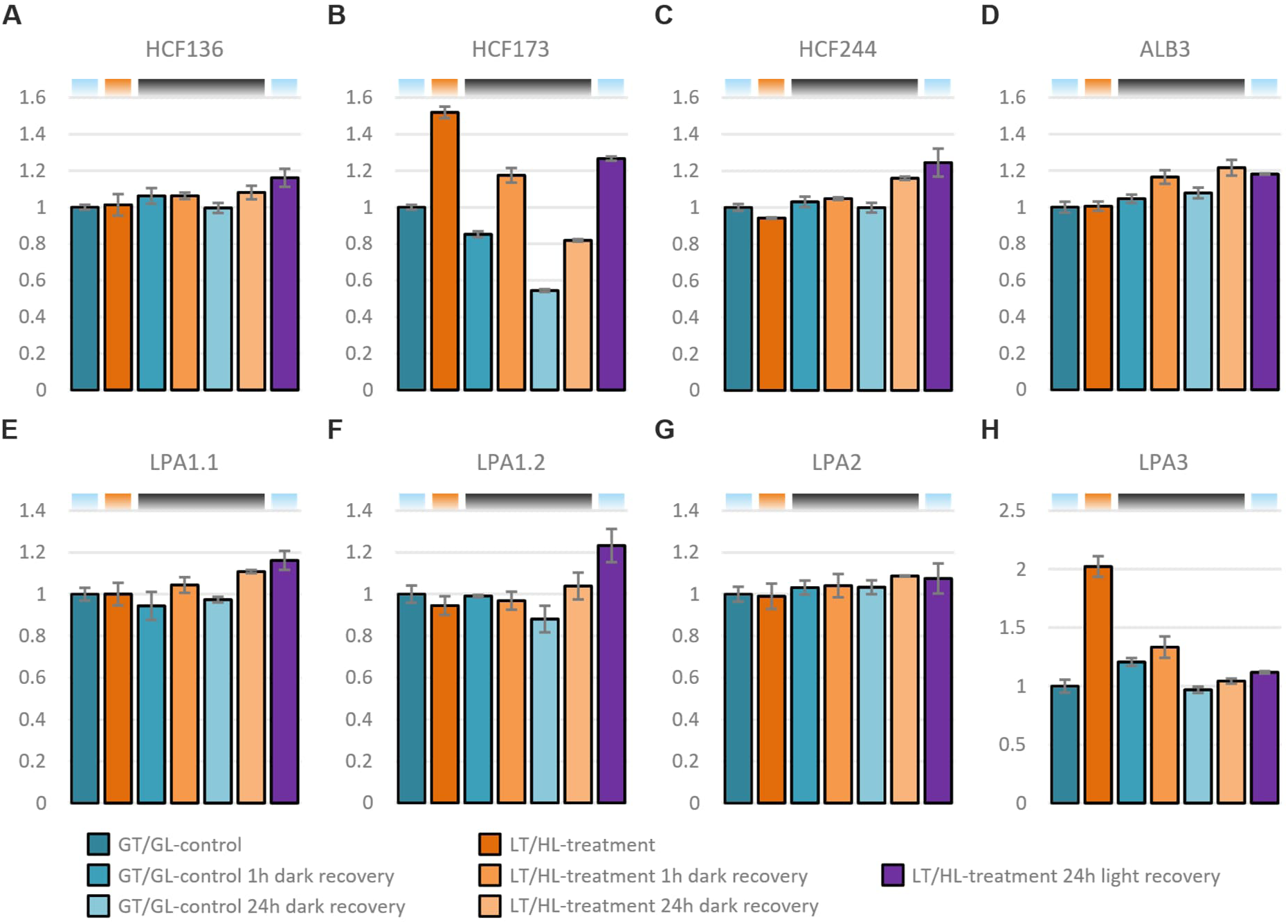
Effect of low temperature and high light treatment, and the subsequent recovery, on PSII repair and assembly factors. **A)** high chlorophyll fluorescence 136 (HCF136) **B)** high chlorophyll fluorescence 173 (HCF173) **C)** high chlorophyll fluorescence 244 (HCF244) **D)** albino 3 (ALB3) **E)** Low PSII accumulation 1.1 (LPA1.1) **F)** Low PSII accumulation 1.2 (LPA1.2) **G)** Low PSII accumulation 2 (LPA2) **H)** Low PSII accumulation 3. (LPA3). Long day-grown lettuce plants were illuminated under 1500 µmol photons m^−2^ s^−1^ of white light at 13 °C for 4 h (LT/HL), while control plants were kept at growth conditions (23 °C and 140 µmol photons m^−2^ s^−1^ with 16 h photoperiod) (GT/GL), after which all plants were transferred to recover for 1 h and 24 h in darkness or for 24 h in long day growth conditions. Thylakoid membranes used in the analyses were isolated directly after the treatment and the recovery periods. Isolated thylakoids were solubilized with detergent and isolated proteins were digested with trypsin. Resulted peptide mixtures were analysed with nLC-ESI-FAIMS-MS/MS in DIA mode and protein abundances were determined with Spectronaut software. Protein abundances were normalised to the average of control plants in growth light. Error bars show standard deviations among technical replicates (n = 3). Coloured bars above the graphs represent the temperature and light conditions from which the thylakoids used in the analyses were isolated: light blue for GT/GL, orange for LT/HL and black for GT/darkness.

The abundance of HCF136, HCF244, and ALB3 increased slightly during the dark recovery of LT/HL treated plants compared to the control (Figures 5A, 5C, and 5D), but in the case of HCF136 and HCF244, the most pronounced increase was observed in plants recovered in the light, where PSII repair is active (Figures 5A and 5C). HCF173 behaved differently and increased in abundance already during the LT/HL treatment (Figure 5B). During the dark recovery, the abundance of HCF173 decreased in both LT/HL treated and GT/GL control lettuce, but it remained at a higher level in the treated lettuce than in the control lettuce (Figure 5B). Among the LPA proteins, only the amount of LPA3 increased during the LT/HL treatment and then decreased to the control levels during the dark recovery (Figures 5E, 5F, 5G, and 5H). These results suggest that HCF173 and LPA3 have specific functions during the LT/HL treatment and subsequent recovery.

#### 3.3.3 Specific LIL proteins show differential accumulation during LT/HL treatment and subsequent recovery

LIL proteins have been associated with sNPQ in previous studies (Demmig-Adams et al. 2006; Zarter et al. 2006a b). Therefore, the accumulation of this protein family was checked after the LT/HL treatment and subsequent dark recovery (Figure 6). The lettuce genome contains several *LIL1* genes encoding the early light-induced proteins (ELIP) (Supplemental file 7). Of these seven isoforms, we detected only two, designated ELIP1.2 and ELIP1.6 (Figures 6A and 6B), in LT/HL treated and recovered samples. Accumulation of ELIP1.2 was induced by the LT/HL treatment and its abundance continued to increase together with ELIP1.6 during 1 h recovery in darkness. Recovery for 24 h in darkness reduced the levels of ELIPs, particularly ELIP1.2, similar to those after the LT/HL treatment, whereas recovery for 24 h in growth conditions returned the levels to control or even lower levels. ELIP1.2 was not detected in control samples, probably due to its low levels in unstressed thylakoids, and for this reason the amount of ELIP1.2 was normalised to that measured in plants that had recovered for 24 h under growth conditions.

**Figure 6.**
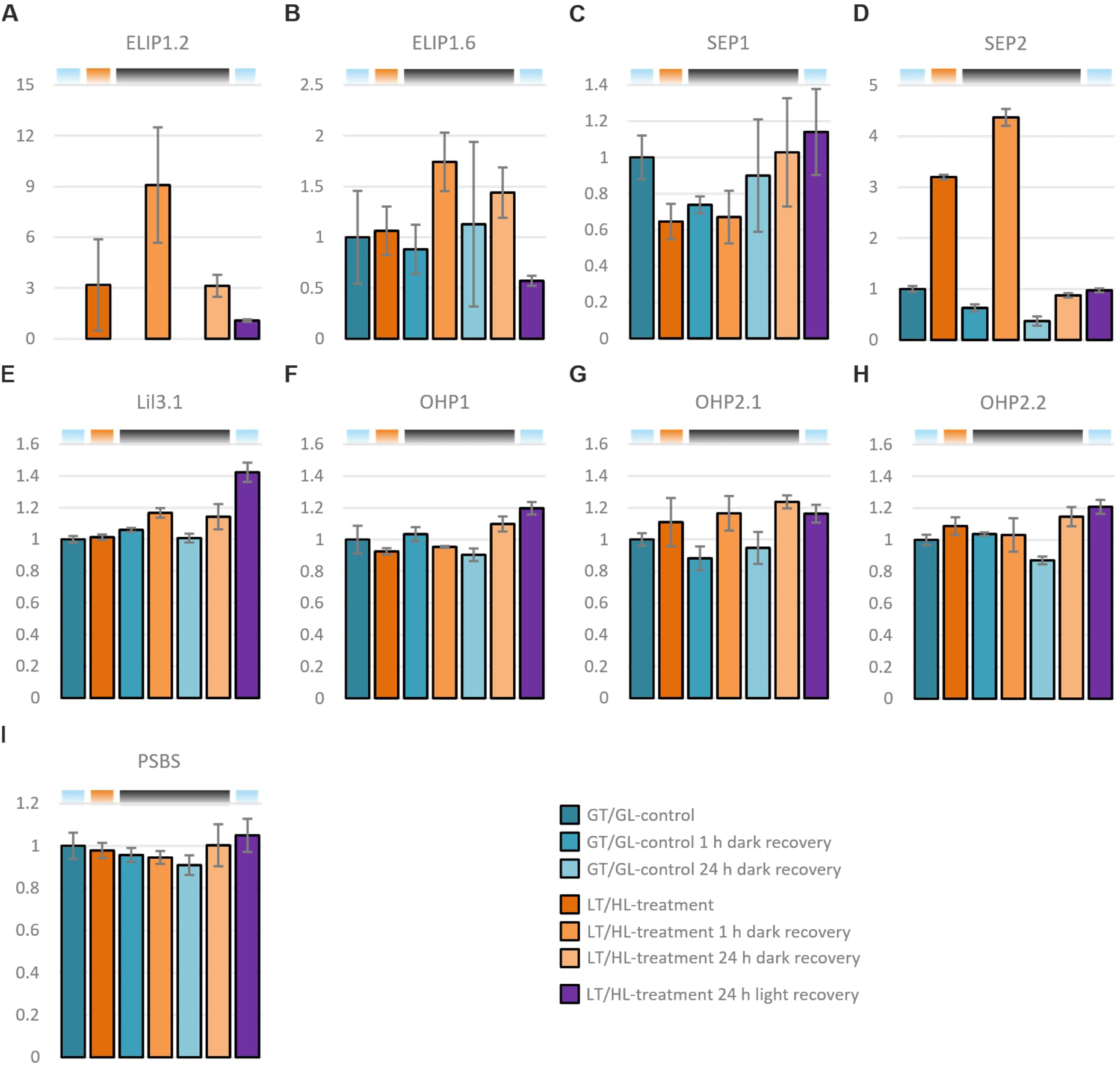
Effect of low temperature and high light treatment, and subsequent recovery, on light-harvesting-like proteins. **A)** early light-induced protein 1.2 (ELIP1.2, normalised to plants recovered for 24 h in growth conditions) **B)** early light-induced protein 1.6 (ELIP1.6) **C)** stress-enhanced protein 1 (SEP1) **D)** stress-enhanced protein 2 (SEP2) **E)** light-harvesting-like 3:1 (LIL3.1 **F)** one helix protein 1 (OHP1) **G)** one helix protein 2.1 (OHP2.1**) H)** one helix protein 2.2 (OHP2.2) **I)** photosystem II subunit S (PSBS). Long day- grown lettuce plants were illuminated under 1500 µmol photons m^−2^ s^−1^ of white light at 13 °C for 4 h (LT/HL), while control plants were kept at growth conditions (23 °C and 140 µmol photons m^−2^ s^−1^ with 16 h photoperiod) (GT/GL), after which all plants were transferred to recover for 1 h and 24 h in darkness or for 24 h in long day growth conditions. Thylakoid membranes used in the analyses were isolated directly after the treatment and the recovery periods. Isolated thylakoids were solubilized with detergent and isolated proteins were digested with trypsin. Resulted peptide mixtures were analysed with nLC-ESI-FAIMS-MS/MS in DIA mode and protein abundances were determined with Spectronaut software. Protein abundances were normalised to the average of control plants in growth light. Error bars show standard deviations among technical replicates (n = 3). Coloured bars above the graphs represent the temperature and light conditions from which the thylakoids used in the analyses were isolated: light blue for GT/GL, orange for LT/HL and black for GT/darkness.

Surprisingly, the two stress-enhanced proteins SEP1 and SEP2 (Heddad & Adamska 2000), encoded by the *LIL4* and *LIL5* genes, behaved differently from each other (Figures 6C and 6D). The amount of SEP1 (quantified on a single peptide basis) decreased during the LT/HL treatment, whereas the amount of SEP2 increased during the treatment and especially during the 1 h recovery in darkness. In contrast to ELIPs, SEP2 levels returned to control levels during the 24 h recovery in darkness, suggesting that these stress-related proteins may have different functions. The levels of other LIL proteins, light-harvesting-like 3:1 (LIL3.1), one helix protein 1 (OHP1), one helix protein 2.1 (OHP2.1) and one helix protein 2.2 (OHP2.2), encoded by the *LIL3.1*, *LIL2*, *LIL6.1* and *LIL6.2* genes, accumulated mainly in plants that had recovered from LT/HL treatment in the light (Figures 6E, 6F, 6G and 6H). OHPs, which are involved in D1 translation in a complex with HCF244 (Hey and Grimm, 2018), showed similar trends of accumulation with HCF244 during the LT/HL treatment and subsequent recovery (Figures 5C, 6F, 6G and 6H). Lettuce has two isoforms for LIL3 (Supplementary file 6), but we could only detect LIL3.1. Finally, the abundance of another LIL protein, PSBS, which is required for fast qE induction, did not change throughout the experiment (Figure 6I). Of all the LIL proteins detected, only the abundance of SEP2, together with ELIP1.2, correlated with sNPQ formation and relaxation.

#### 3.3.4 LT/HL treatment and subsequent recovery alter the amounts of proteins involved in the metabolism of photosynthetic pigments

Since zeaxanthin accumulation is an important part of qZ, we focused on enzymes involved in pigment biosynthesis and the xanthophyll cycle (Figure 7). The LT/HL treatment resulted in a large increase in the amount of β-carotene 3-hydroxylase (BCH2) (Figure 7D), which synthesises zeaxanthin from β-carotene.

**Figure 7.**
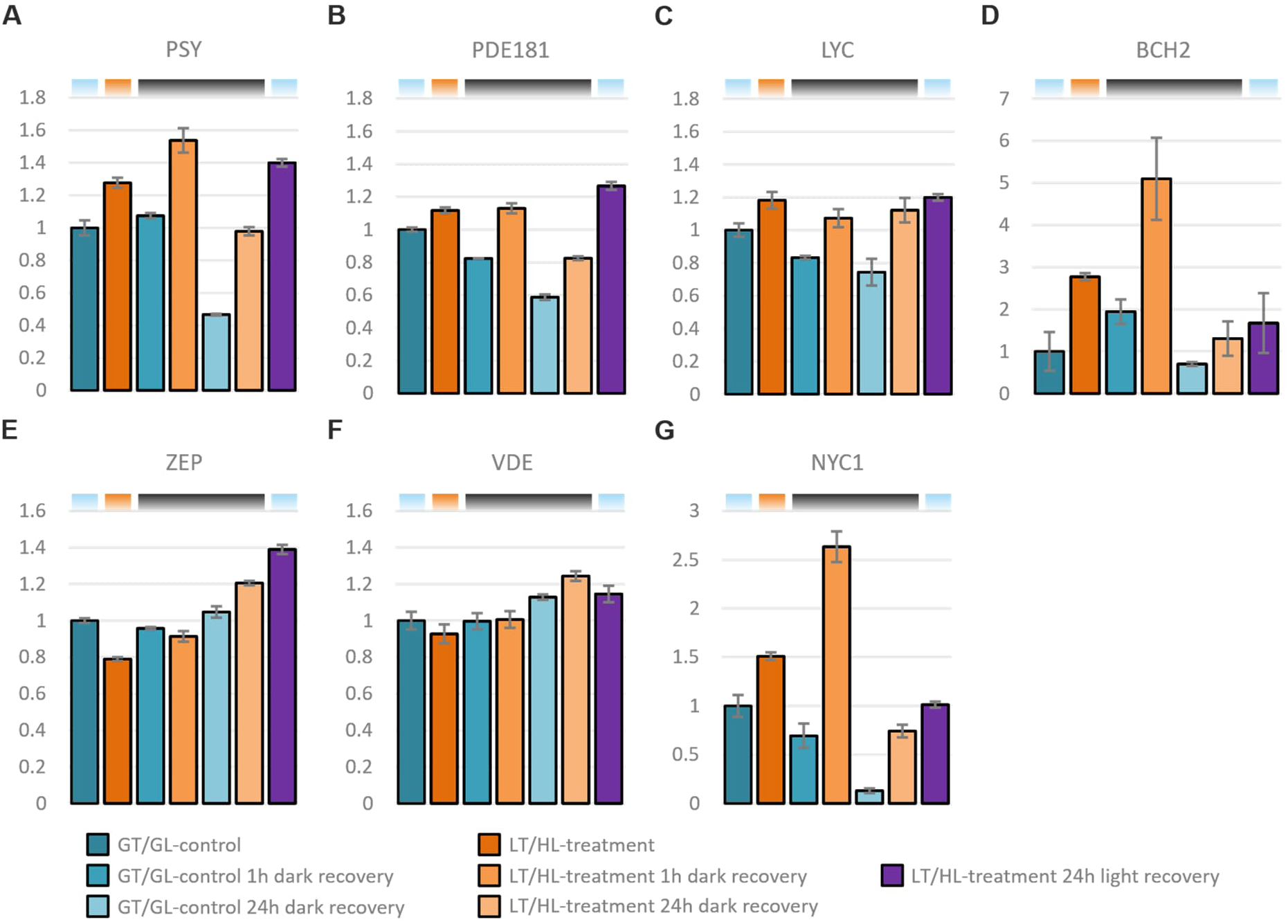
Effect of low temperature and high light treatment, and the subsequent recovery, on enzymes in the metabolism of photosynthetic pigments. **A)** Phytoene synthase (PSY) **B)** ζ-carotene desaturase (PDE181) **C)** Lycopene β-cyclase (LYC) **D)** β-carotene hydroxylase (BCH2) **E)** Zeaxanthin epoxidase (ZEP) **F)** Violaxanthin de-epoxidase (VDE) **G)** Chlorophyll b reductase (NYC1). Long day-grown lettuce plants were illuminated under 1500 µmol photons m^−2^ s^−1^ of white light at 13 °C for 4 h (LT/HL), while control plants were kept at growth conditions (23 °C and 140 µmol photons m^−2^ s^−1^ with 16 h photoperiod) (GT/GL), after which all plants were transferred to recover for 1 h and 24 h in darkness or for 24 h in long day growth conditions. Thylakoid membranes used in the analyses were isolated directly after the treatment and the recovery periods. Isolated thylakoids were solubilized with detergent and isolated proteins were digested with trypsin. Resulted peptide mixtures were analysed with nLC-ESI-FAIMS-MS/MS in DIA mode and protein abundances were determined with Spectronaut software. Protein abundances were normalised to the average of control plants in growth light. Error bars show standard deviations among technical replicates (n = 3). Coloured bars above the graphs represent the temperature and light conditions from which the thylakoids used in the analyses were isolated: light blue for GT/GL, orange for LT/HL and black for GT/darkness.

BCH2 further increased during the 1 h recovery in darkness before returning to control levels after the 24 h recovery in darkness or growth conditions. BCH2 is upregulated by β-ionone, an oxidation product of β-carotene (Felemban et al., 2023), suggesting that β-carotene oxidation induces zeaxanthin synthesis. In addition, three enzymes leading to β-carotene formation, the 15-cis phytoene synthase (PSY), zeta-carotene desaturase (PDE181), and lycopene β-cyclase (LYC), had slightly higher abundances after LT/HL treatment when compared to GT/GL control and the amounts did not decrease as much during the dark recovery as in the controls (Figures 7A, 7B and 7C). The levels of the xanthophyll cycle enzymes, zeaxanthin epoxidase (ZEP) and violaxanthin de-epoxidase (VDE), were not induced, but in the case of ZEP actually decreased by LT/HL treatment, while an increase in both was observed after 24 h recovery period (Figures 7E and 7F). ZEP levels have been shown to decrease during HL treatment (Bethmann et al., 2019), but in addition, we detected an increasing trend in ZEP and VDE levels during dark and light recovery after the LT/HL treatment. In addition, the LT/HL treatment also increased the amount of Chl b reductase (NYC1), which reached the highest level during 1 h dark recovery compared to GT/GL controls (Figure 7G). NYC1 catalyses the first step of Chl b degradation, which is associated with the degradation of LHCII under HL illumination (Sato et al., 2015).

#### 3.3.5 LT/HL treatment and subsequent recovery have little effect on the abundance of enzymes involved in thylakoid protein phosphorylation or proteins required for qH induction and relaxation

To gain more information about the regulation of photosynthesis in our experimental conditions, we analysed the proteins involved in thylakoid protein phosphorylation and qH (Supplemental figure 1). State transition 7 (STN7) phosphorylates LHCII, which is required for formation of the LHCII-PSI complex, whereas State transition 8 (STN8) phosphorylates the PSII core proteins CP43, D1, D2, PSBH and the minor antenna protein LHCB4. LT/HL treatment had little effect on the abundance of the two kinases, but a moderate increase occurred during the 24 h recovery period for both STN8 and STN7 (Supplemental figure 1A and 1B). The levels of thylakoid-associated phosphatase 38 (TAP38) and photosystem II core phosphatase (PBCP) decreased during the LT/HL treatment, and both recovered to GT/GL control levels during dark recovery (Supplemental figure 1C and 1D). Levels of STN8, TAP38 and PBCP were highest in plants that had recovered for 24 h under growth conditions.

The amount of chloroplastic lipocalin (CHL), which is required for qH formation in LHCII trimers, was not altered by the LT/HL treatment or the subsequent recovery (Supplemental figure 1G). The amount of suppressor of quenching 1 (SOQ1), which inhibits CHL, decreased slightly during the LT/HL treatment but was restored during recovery, where it even increased to a higher level than in control (Supplemental figure 1E). The amount of relaxation of qH (ROQH), which discharges the quenching at qH active LHCII, showed more changes in abundance. The abundance decreased slightly during the LT/HL treatment, and even more during dark recovery but, in this case, the decrease occurred both in GT/GL control and LT/HL treated plants, suggesting that in lettuce the putative qH relaxation is not related to the LT/HL treatment and PSII recovery from photoinhibition (Supplemental figure 1F).

#### 3.3.6 LT/HL treatment and subsequent recovery induce phosphorylation dynamics of the PSII core proteins and the minor LHCII antenna protein LHCB4.1

Following the detection of LHCB4 phosphorylation by p-Thr antibody and phosphoprotein staining of 2D gels (Figures 3A and 3E), the LHCB4 phosphorylation was further investigated by quantifying p-LHCB4.1 and identifying the phosphosite from our MS data. In parallel to LHCB4.1, we also examined the phosphorylation dynamics of the PSII core proteins D1, D2 and PSBH (Figure 8A, 8B and 8C), which are known targets of the STN8 kinase. Since the phosphorylation and dephosphorylation of the D1 and D2 proteins are important parts of the PSII repair cycle, it was of interest to analyse the phosphorylation dynamics of these proteins. LT/HL treatment induced N-terminal D1 and D2 phosphorylation, which remained high during 1 h dark recovery but decreased within 24 h dark recovery (Figures 8A and 8B). PSBH, a small PSII subunit with a transmembrane helix and a long stromal extension located at the interface of CP47 and LHCB4 (Su et al., 2017), was also phosphorylated during the LT/HL treatment, but curiously, the phosphorylation of T3 and T5 increased even further during 1 h dark recovery until it returned closer to the control levels after 24 h dark recovery (Figure 8C). The effect of LT/HL treatment on LHCB4.1 T109 phosphorylation was even more dramatic, being 20-fold higher in LT/HL-treated plants compared to control plants, but rapidly returning to control levels during 1 h dark recovery (Figure 8D). LHCB4.1 T109 phosphorylation site is located in the stromal loop of LHCB4.1, which in the PSII-LHCII complex is bound to CP47 on the stromal side of the complex (Supplemental figure 2). Therefore, the phosphorylation of T109 could affect the interaction between LHCB4.1 and CP47, possibly explaining the detachment of M-LHCII from the PSII core (Figure 3B and 3C).

**Figure 8.**
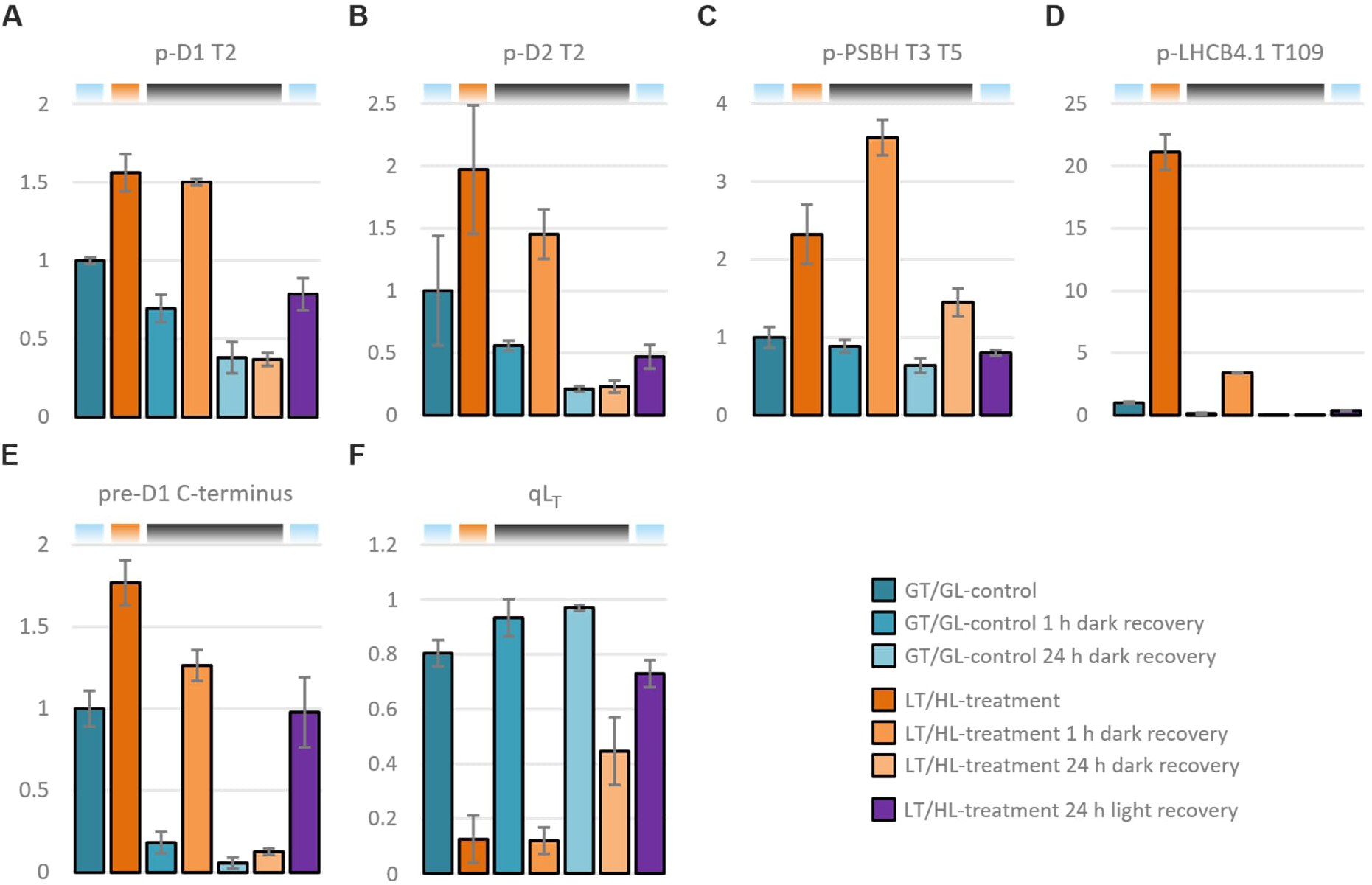
Effect of low temperature and high light treatment, and the subsequent recovery, on phosphorylation of the PSII core proteins and the minor antenna LHCB4, D1 processing and recovery of PSII function. **A)** p-D1 T2 **B)** p-D2 T2 **C)** p-LHCB4.1 T109 **D)** 2p**-** PSBH T3 T5 **E)** pre-D1 C-terminus **F)** fraction of open and functional PSII reaction center. Long day-grown lettuce plants were illuminated under 1500 µmol photons m^−2^ s^−1^ of white light at 13 °C for 4 h (LT/HL), while control plants were kept at growth conditions (23 °C and 140 µmol photons m^−2^ s^−1^ with 16 h photoperiod) (GT/GL), after which all plants were transferred to recover for 1 h and 24 h in darkness or for 24 h in long day growth conditions. Thylakoid membranes used in the analyses were isolated directly after the treatment and the recovery periods. Isolated thylakoids were solubilized with detergent and isolated proteins were digested with trypsin. Resulted peptide mixtures were analysed with nLC-ESI-FAIMS-MS/MS in DIA mode and peptide abundances were determined with Spectronaut software. Peptide abundances were normalised to the average of control plants in growth light. Specific peptides for p-D1 T2, p-D2 T2, 2p-PSBH T3 T5, p-LHCB4.1 T109 and pre D1 are [N-acetyl]T[p]AILER, [N-acetyl]T[p]IALGKVTK, AT[p]QT[p]VENGAR, NLAGDVIGT[p]RFEDADVK and NAHNFPLDLAAIEAPSTNG respectively. F0’, F0, FM’, and FM were determined with Fluorpen directly after the treatment and after subsequent recovery periods of 1 h and 24 h. qLT was calculated as ((1/ F0’)-(1/ FM’))/((1/ F0^ref^)-(1/ FM^ref^)) using as the reference values (FM^ref^ and F0^ref^) the averages of FM and F0 from control plants shifted to darkness for 24 h. Error bars show standard deviations among technical replicates (A-F) (n = 3) and biological replicates (G) (n = 4-16). Coloured bars above the graphs represent the temperature and light conditions from which the thylakoids used in the analyses were isolated: light blue for GT/GL, orange for LT/HL and black for GT/darkness.

#### 3.3.7. LT/HL treatment leads to the accumulation of pre-D1 protein, which allows partial restoration of PSII function during dark recovery

More detailed analysis of C-terminal pre-D1 peptide revealed changes during the LT/HL treatment in processing of the C-terminal extension of D1 protein by CTPA (Figures 8E), which is required for the assembly of functional PSII complexes (Che et al., 2013), indicating slow D1 turnover and PSII repair during the LT/HL treatment. Conversely, the pre-D1 levels decreased during subsequent dark recovery both in control and LT/HL treated plants, implying that D1 protein C-terminal processing, and also some later steps of PSII repair, can proceed in the dark (Pavlovič et al., 2016). The cleavage of D1 C-terminal extension by CTPA allows the assembly of photochemically functional PSII (Che et al., 2013).

Since the PSII quantum yield, Y(II), also increased during dark recovery of LT/HL treated plants (Figure 2A), we next calculated the fraction of open and functional PSII centres (qLT) (Porcar-Castell, 2011) in control and LT/HL-treated plants (figure 8F). qLT can be used to assess the amount of functional PSII centres, since the PQ pool is mostly oxidized in darkness. qLT was low, 0.13, directly after the LT/HL-treatment but increased to 0.44 during 24 h in dark recovery conditions. These results strongly suggest that directly after the LT/HL-treatment, a great portion of the mature D1 was damaged in photoinhibited PSII RCs and the recovery in qLT resulted from resumed processing of the D1 C-terminal extension by CTPA.

### 3.4 LT/HL treatment and subsequent recovery alter the pigment composition of lettuce thylakoids

Since our proteomic analysis suggested an increase in Chl b catabolism and zeaxanthin synthesis after the LT/HL treatment and especially after 1 h dark recovery (Figure 7), this prompted us to analyse the pigment composition of isolated thylakoids. However, the decrease in the relative amount of Chl b was small and only visible after 24 h of dark recovery (Figure 9A). In contrast, the decrease in β-carotene, mainly bound to the PSI and PSII cores, occurred already during the LT/HL treatment (Figure 9B), which could be related to minor degradation of PSII core proteins and hydroxylation of released β-carotene to zeaxanthin (Figures 4A-D) (Beisel et al., 2010). LT/HL treatment and subsequent recovery also increased the levels of all xanthophylls, with lactucaxanthin and neoxanthin being less affected (Figures 9C-9H), consistent with previous studies (Esteban et al., 2015) and our proteomic analysis (Figure 7).

**Figure 9.**
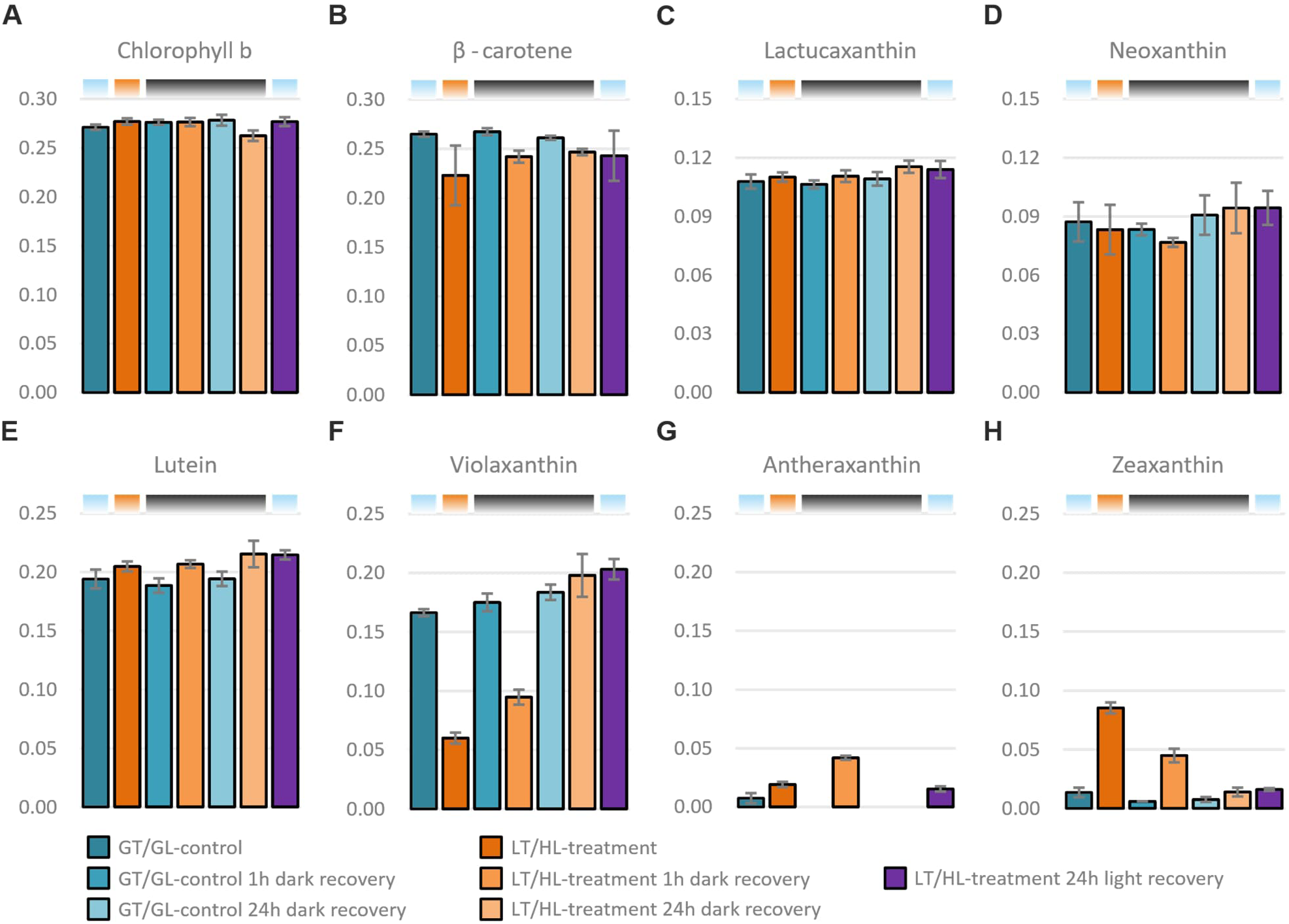
Effect of low temperature and high light treatment, and the subsequent recovery, on photosynthetic pigments. **A)** Chlorophyll b **B)** β-carotene **C)** Lactucaxanthin **D)** Neoxanthin **E)** Lutein **F)** Violaxanthin **G)** Antheraxanthin **H)** Zeaxanthin. Long day-grown lettuce plants were illuminated under 1500 µmol photons m^−2^ s^−1^ of white light at 13 °C for 4 h (LT/HL), while control plants were kept at growth conditions (23 °C and 140 µmol photons m^−2^ s^−1^ with 16 h photoperiod) (GT/GL), after which all plants were transferred to recover for 1 h and 24 h in darkness or for 24 h in long day growth conditions. Thylakoid membranes used in the analyses were isolated directly after the treatment and the recovery periods. Pigments were extracted from isolated thylakoids with acetone and the extracts were analysed with HPLC-DAD. Pigments were identified by absorbance spectra and relative retention times. Relative pigment abundances were estimated by the area of chromatographic peaks detected at 440 nm and normalised within the sample to the area of Chl a. Error bars show standard deviations among four biological replicates (n = 4). Coloured bars above the graphs represent the temperature and light conditions from which the thylakoids used in the analyses were isolated: light blue for GT/GL, orange for LT/HL and black for GT/darkness.

## 4. Discussion

### 4.1 Canonical NPQ mechanisms do not play a major role in LT/HL treatment-induced sNPQ in lettuce

qE is the fastest NPQ component induced in the light and depends on lumen acidification followed by protonation of the PSBS protein, with the strength of qE proportional to the expression level of the PSBS protein (Li et al., 2002). Because qE is dependent on lumen acidification, it relaxes within seconds in the dark (Niyogi and Truong, 2013). In our lettuce experiments, we used 1 h as the shortest dark recovery time, which allowed about 70% relaxation of the NPQ induced by GT/HL treatment, but only about 30% relaxation of the NPQ induced by LT/HL treatment (Figures 2B and 2F). The amount of PSBS remained stable throughout the LT/HL treatment and subsequent recovery periods (Figure 6I), suggesting that the magnitude of qE during LT/HL treatment is similar to that during the GT/HL treatment, and according to NPQ measurements, qE appears to be fully relaxed within 1 h dark recovery (Figures 2B and 2F).

Slightly slower relaxing NPQ mechanisms in plants compared to qE have been attributed to qZ, which is based on the retention of zeaxanthin in the external LHCII antenna (Nilkens et al., 2010). Zeaxanthin is formed in the light-induced xanthophyll cycle under abiotic stress conditions (Demmig-Adams and Adams, 2006) and promotes the quenching of excitation energy in the external LHCII antenna systems (Bassi and Dall’Osto, 2021; Ruban and Saccon, 2022). By the end of the 1 h dark recovery, about half of the zeaxanthin had been converted to antheraxanthin (Figures 9G and 9H), which is unlikely to contribute to qZ because zeaxanthin de-epoxidation releases the formed antheraxanthin from LHCII into the lipid bilayer (Küster et al., 2023). Therefore, the remaining amount of zeaxanthin can only account for part of the high sNPQ remaining after 1 h dark recovery of LT/HL treated lettuce (Figures 2B and 9H). Furthermore, zeaxanthin accumulation does not correlate with sNPQ in all plant species (Míguez et al., 2015; Míguez et al., 2017). Taken together, these results suggest that the LT/HL treatment-induced sNPQ, which is preserved after 1 h of dark recovery, does not represent the PSBS or zeaxanthin-dependent NPQ, and therefore other mechanisms must exist for sNPQ in lettuce.

We also considered a different form of sNPQ, a lipocalin-dependent qH described in Arabidopsis (Malnoë, 2018). However, its involvement in the sNPQ generated in lettuce during the LT/HL treatment and its subsequent relaxation during dark recovery is difficult to assess, due to the lack of clear molecular signatures to track qH. The known qH-related proteins, CHL, SOQ1, and ROQH1, showed no differential expression in LT/HL treated lettuce compared to GT/GL control (Supplemental figures 1E-G). Slightly reduced fluorescence of L-LHCII trimers in CN gels has been attributed to qH in Arabidopsis (Bru et al., 2022). In lettuce, however, no apparent changes in the fluorescence of L-LHCII trimers were observed during LT/HL treatment or subsequent recovery phases (Figure 3C), suggesting that qH is not particularly active under the stress conditions applied here to lettuce.

### 4.2. Role of LHCB4 phosphorylation in the formation of sNPQ in LT/HL treated lettuce?

Canonical LHCII phosphorylation-dependent quenching (qT) is not expected to occur under HL in plants, because the STN7 kinase, which phosphorylates LHCB1 and LHCB2, is inhibited under such conditions, preventing the formation of the LHCII-PSI sc (Rintamäki et al., 2000; Bassi and Dall’Osto, 2021; Cutolo et al., 2023). As expected, after the LT/HL-treatment of lettuce, we detected almost complete dephosphorylation of LHCB1 and LHCB2. However, a major phosphorylation of LHCB4 appeared during the LT/HL treatment (Figure 3A), which in angiosperms has previously been reported only in grasses (Chen et al., 2013; Betterle et al., 2017). Grasses, lettuce and most angiosperms have two isoforms of LHCB4 (LHCB4.1 and LHCB4.2), whereas Arabidopsis, pea (*Pisum sativum*), and spinach (*Spinacia oleracea*), which are commonly used in photosynthesis research have three isoforms (LHCB4.1, LHCB4.2, and LHCB4.3), of which the LHCB4.3 is expressed only in HL, leading to dissociation of the M-LHCII trimer and LHCB6 from the PSII core (Albanese et al., 2019; Grebe et al., 2019).

Phosphorylation of lettuce LHCB4 was found to occur on the stromal loop, at the site that serves as a binding site for CP47 in unphosphorylated LHCB4 (Supplemental figure 2). It is therefore conceivable that the STN8-dependent phosphorylation of LHCB4 (Betterle et al., 2017) leads to the dissociation of p-LHCB4, together with the attached M-LHCII trimer and LHCB6 (i.e. the entire M-LHCII complex), from the PSII core (Figures 3B and 12), performing the same function as LHCB4.3 in other species. This interpretation is consistent with the absence of p-LHCB4 in PSII sc (Figures 3D and 3E). The dissociation of the M-LHCII complex from the PSII core during the LT/HL-treatment (Figure 3B) would decrease the relative antenna size of PSII, which is consistent with the increase in the F735/F685 ratio (Figure 3F). However, the increased F735/F685 ratio, partially disassembled PSII complexes, and high levels of free M-LHCII were maintained after 1 h recovery in darkness, even though the LHCB4. proteins were already dephosphorylated (Figures 3A, 3B, and 3F). This probably means that the reassociation of M-LHCII with the PSII core is prevented by some other mechanism(s), as will be discussed in the following chapters. The reduction in PSII antenna size can explain part of the sNPQ, but based on the F735/F685 ratio there must be also other mechanism to explain the high sNPQ.

### 4.3. Regulation of PSII repair leads to pausing of D1 C-terminal processing and inhibition of D1 degradation during LT/HL treatment and subsequent dark recovery leading to accumulation of different PSII populations

Our analysis of PSII repair-associated proteins revealed that LT/HL-treatment induced a clear accumulation of HCF173 and LPA3 proteins (Figures 5B and 5H). HCF173 binds *psbA* transcripts, recruits D1-translating polysomes to the thylakoid membrane and facilitates the insertion of nascent D1-chains into partially assembled PSII subcomplexes (Schult et al., 2007; Link et al., 2012; Chotewutmontri and Barkan, 2020; Wang and Grimm, 2021), and LPA3 facilitates the repair of damaged PSII (Cai et al., 2010). In contrast, the majority of the proteins involved in the later steps of the PSII repair cycle were not upregulated (Figures 5A, 5C, 5D-G and 6F-H).

A clear pausing of PSII repair during the LT/HL treatment occurred at the level of pre-D1 processing (Figure 8E), which proceeded more slowly than the HCF173 facilitated translation. Accumulation of the pre-D1 protein likely results from inhibition of pre-D1 C-terminal processing protease (CTPA) by the thioredoxin system (Hall et al., 2010; Järvi et al., 2015), linking the regulation of PSII repair to stromal redox state. Subsequent transfer of lettuce to recovery conditions in the dark, where no new initiations of D1 translation take place (Chotewutmontri and Barkan, 2018), revealed the conversion of accumulated pre-D1 to mature D1 by activation of CTPA.

Not only the processing of pre-D1, but also the degradation of damaged D1 proteins, seems to be stalled during the LT/HL treatment and subsequent dark recovery and a great portion of the PSII centers with mature D1 protein were non-functional as deduced from the low qLT (Figures 8F and 8G). Those PSIIs most likely represent PSII complexes that need a replacement of damaged D1 protein with a newly synthesised one. The accumulation of damaged D1 relates to the elevated N-terminal phosphorylation of D1 protein (Figure 8A), which is known to prevent D1 degradation (Rintamäki et al., 1996; Kato and Sakamoto, 2014; Puthiyaveetil et al., 2014). Moreover, a slow dephosphorylation of D1 and D2 during the LT/HL treatment, in comparison to control plants (Figure 8A), implies that PBCP, the PSII core protein phosphatase, is inhibited by the LT/HL treatment (Liu et al., 2019). Thus, the degradation of damaged D1 is strictly regulated not only by kinases but also but also by phosphatases.

Based on the comparisons of changes in D1 protein levels (Figure 4A) and in the amount of pre-D1 peptide (Figure 8E), with qLT values (Figure 8F), we estimate that during LT/HL treatment up to 20% of PSII RCs are degraded, approximately 40% of the PSII RCs have a damaged D1 protein and 30% of the PSII RCs have a pre-D1, while only 10% of the PSII RCs are functional (Supplemental figure 3). In comparison, the control plants without the LT/HL treatment, comprised 80% active PSIIs with mature D1 protein, 15% with pre-D1 and 5% with damaged D1 protein. Recovery in darkness for 24 h after the LT/HL treatment allowed the processing of the C-terminal extension of the p-D1 protein by CTPA (Figure 8E), but the full recovery of PSII eventually requires also light (Figure 8F).

The accumulation of pre-D1 during the LT/HL treatment coincides with the accumulation of the LPA3 protein (Figures 8E and 5H), which has been shown to interact with CP43 during the PSII repair cycle in Arabidopsis (Cai et al., 2010). D1 C-terminal extension prevents the correct binding of CP43 to D1 during the repair cycle, leading to accumulation of free CP43 (Che et al., 2013; Shi et al., 2021). The recently reported spectroscopic studies show that free CP43 isolated from Synechocystis (*Synechocystis* sp. PCC 6803) (Biswas et al., 2023) is poorly protected under excess illumination, leading the authors to speculate that some weakly interacting protein, lost during isolation, is likely to quench the free CP43 under more natural conditions. Considering these two reports, we propose that LPA3 binds to CP43 released from the PSII core upon initiation of the PSII repair cycle and protects the free CP43 from light-induced damage and proteolytic degradation (Figures 4C, 5H, and 10A).

### 4.4 SEP2-dependent quenching of the PSII inner antenna CP47 as a putative mechanism for sNPQ in lettuce

Among the LIL proteins in lettuce, the accumulation dynamics of SEP2 protein coincided with the formation and relaxation of sNPQ (Figures 2B and 6D). Plant LIL proteins have evolved from cyanobacterial high light-induced proteins (Hlip), which are replaced with the OHP and SEP proteins in the green lineage (Engelken et al., 2010). Synechocystis has four Hlips that are involved in PSII assembly and repair (Komenda and Sobotka, 2016). The HliCD heterodimer delivers pigments to the newly translated D1 protein and protects the complex from light-induced damage (Knoppová et al., 2014; Staleva et al., 2015). In plants, the OHP proteins are functional equivalents of Synechocystis HliC and HliD, but in lettuce OHPs showed only minor changes in abundance during the LT/HL treatment or subsequent recovery (Figure 6F-H). On the other hand, Synechocystis HliA and HliB form heterodimers with HliC and the dimers bind to free CP47, and to the PSII assembly intermediate RC47 complex (Konert et al., 2022). The HliAC heterodimer appears to be involved in PSII repair, whereas HliBC heterodimer functions in the assembly of CP47 (Rahimzadeh-Karvansara et al., 2022). Related to different functions of HliA and HliB in Synechocystis during HL stress (Konert et al., 2022; Rahimzadeh-Karvansara et al., 2022), we found similarities in the behaviour of the SEP1 and SEP2 proteins in lettuce under our experimental conditions (Figures 6C and 6D). While the amount of SEP1 decreased during the LT/HL treatment (Figure 6C), the SEP2 levels increased substantially and coincided with apparent stalling of the PSII repair cycle. These evolutionary and functional considerations led us to hypothesise that lettuce SEP2 binds to PSII repair intermediates and quenches them (Figures 3C and 10A).

**Figure 10.**
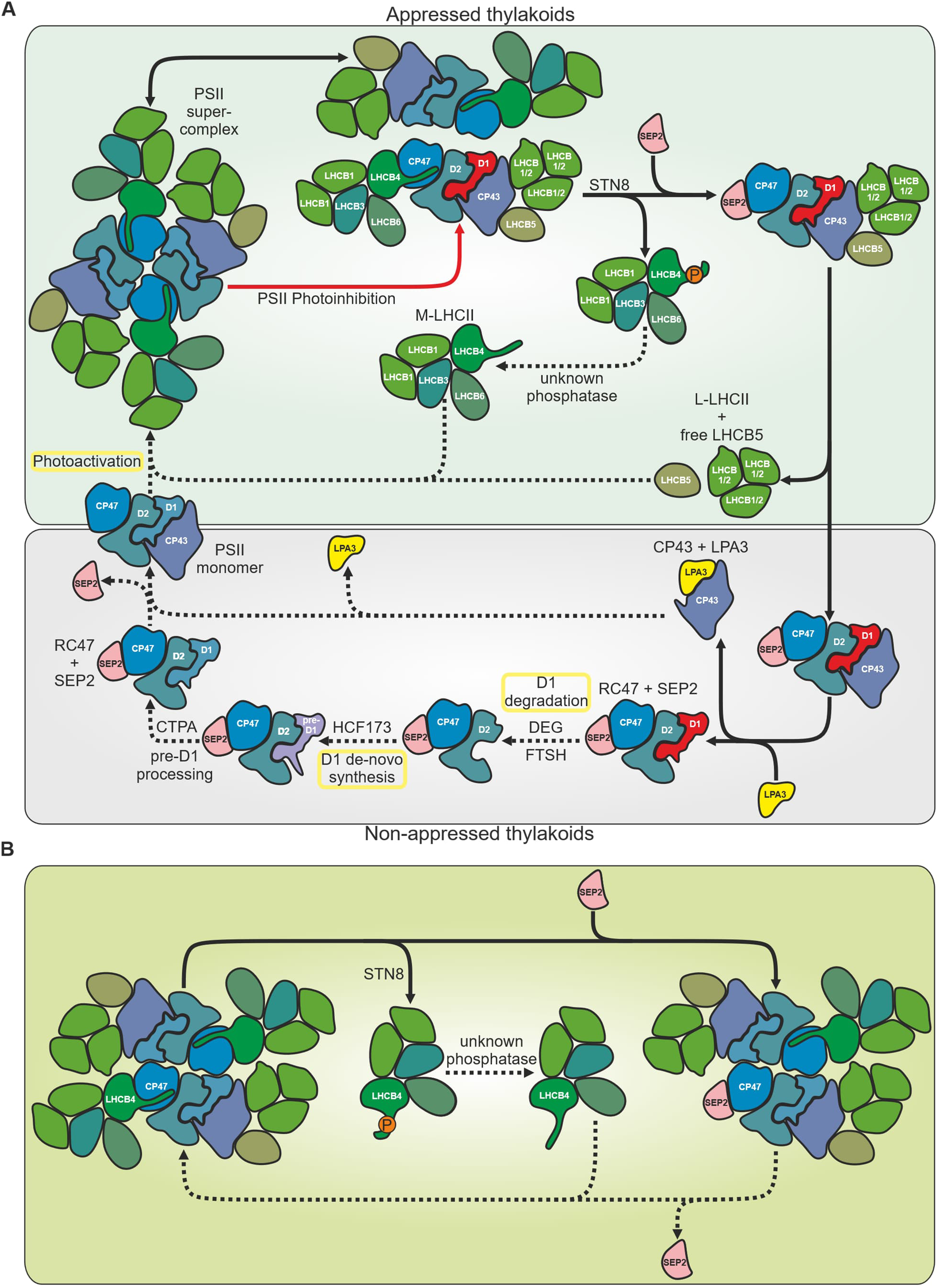
Proposed roles of STN8-dependent LHCB4 phosphorylation, SEP2, and LPA3 proteins in photoprotection of stalled PSII repair cycle (A) and formation of sNPQ in functional PSII complexes (B) in lettuce during the LT/HL treatment and subsequent recovery. Solid arrows represent processes occurring during the LT/HL treatment: STN8-dependent phosphorylation of LHCB4 leads to the dissociation of M-LHCII from the PSII sc. The release of M-LHCII allows SEP2 to bind to CP47 and quench the excitation energy at the inner antenna of PSII, either in damaged (A) or functional (B) PSII cores. Release of CP43 from damaged PSII complexes allows the degradation of the non-phosphorylated damaged D1 protein. LPA3 binds to free CP43 protecting it from photodamage and degradation (A). Dashed arrows represent processes that are slowed down during the LT/HL treatment and resumed during the subsequent recovery: Dephosphorylation of LHCB4 and degradation of SEP2 allow reassociation of M-LHCII with PSII cores (A and B). The paused PSII repair cycle is completed by degradation of dephosphorylated damaged D1 and co-translational insertion of a new D1 copy into the RC47 complex. CTPA catalysed pre-D1 processing allows the re-binding of CP43 and photoactivation of PSII (A). Processes marked with a yellow rectangle are light dependent and therefore do not occur during dark recovery. PSII photoinhibition and damaged D1 proteins are shown in red and phosphorylation of the PSII core proteins is omitted for clarity.

Further analysis of evolutionary aspects revealed that Synechocystis HliA and HliB interact also with the Psb35 protein, which is thought to protect CP47, HliA, and HliB from FtsH-mediated degradation by covering the N-terminal parts of these proteins (Pascual-Aznar et al., 2021). Lettuce, and plants in general, do not have Psb35 but the second helix of SEP1 shares some homology with Psb35 (Pascual-Aznar et al., 2021). Moreover, Psb35 and the HliA and HliB proteins are fused in some Synechococcus strains (Kilian et al., 2008) suggesting that such an early evolutionary event could be linked to the evolution of SEPs and OHPs in the green lineage. Intriguingly, the HliA and HliB proteins have been proposed to bind to CP47 in Synechocystis at the same site where LHCB4 in plants binds to CP47 (Pascual-Aznar et al., 2021). It is therefore conceivable that SEP2 binds to the same site in both damaged and functional PSII complexes. In the latter case, the binding of SEP2 to CP47 would explain why LHCB4 in lettuce, after dephosphorylation, does not rebind to the PSII core during 1 h recovery in darkness (Figures 3A, 3C, and 8D). The binding of SEP2 to CP47 could promote energy dissipation as heat rather than directing light to the PSII RC. Such a competition between the binding of SEP2 and LHCB4 to the PSII core would provide an additional regulatory loop between light harvesting and photoprotection in lettuce (Figure 10B).

### 4.5 Upregulation of ELIP1.2 may provide safe storage for pigments released from degraded PSII cores and LHCII antenna proteins

ELIPs, one group of the LIL proteins, have been proposed to function as pigment carriers (Adamska, 1997; Tzvetkova-Chevolleau et al., 2007). In lettuce, ELIP1.2 accumulation occurred during the LT/HL treatment but even more prominently during the subsequent 1 h recovery in the dark (Figure 6A), coinciding with the degradation of the damaged PSII cores (Figures 4A-D), leading us to suggest that Chls released from damaged PSII cores are bound to ELIP1.2 under conditions where the PSII repair cycle is not fully active (Supplemental figure 4) (Beisel et al., 2010). A similar coincidence between the accumulation of damaged PSII cores and the accumulation of ELIPs has been reported for overwintering evergreens (Zarter et al., 2006a; Zarter et al., 2006b). The LT/HL treatment of lettuce also induced the synthesis of zeaxanthin, most likely by BCH2 catalysed hydrolysis of the β-carotene released from the degradation of damaged PSII RCs (Figures-3A-D, 7D, 9B, and 9H), as the abundance of BCH2 increased already during the LT/HL-treatment but especially during the 1 h recovery in the dark (Figure 7D). ELIPs are known to also bind zeaxanthin (Rossini et al., 2006; Skotnicová et al., 2021), which would allow quenching of the bound Chls. In summary, ELIP1.2 is proposed to act as a safe storage site for Chl a until it is reused for PSII repair or ultimately degraded.

In addition to binding Chl a from damaged PSII complexes, lettuce ELIP1.2 may have also another role in rescuing both Chl a and Chl b from degraded LHCII (Supplemental figure 4). This interpretation is based on the fact that the abundance of the NYC1, which catalyses the first step of converting Chl b to Chl a, was upregulated in our experiment, especially after 1 h of dark recovery (Figure 7G). On the other hand, the accumulation of ELIP2 in Arabidopsis under constitutive expression has been shown to down- regulate Chl biosynthesis (Tzvetkova-Chevolleau et al., 2007). This suggests that the formation of ELIP1.2- dependent Chl stores may also regulate Chl metabolism by inhibiting Chl biosynthesis, when there is a high level of stored Chl. This hypothesis of ELIPs as safe pigment stores, would also imply that ELIPs do not play a role in the formation of sNPQ, which is consistent with unchanged Y(II) in the ELIP-overexpressing Arabidopsis (Tzvetkova-Chevolleau et al., 2007).

## 5. Conclusions

Lettuce plants form strong NPQ during the LT/HL treatment due to excessive excitation stress. Based on biochemical, biophysical, and quantitative targeted proteome analyses, we propose that the STN8 kinase-dependent phosphorylation of LHCB4, detected in grasses and lettuce among the land plants studied, initially dissociates the M-LHCII complex from PSII sc (Figure 10). This reduces the excitation energy transfer to the PSII cores and simultaneously increases the excitation lifetime in the shared LHCII pool, which increases the probability of excitation energy quenching by the qE-active LHCII trimers, the qZ-active minor LHCII antennas (LHCB4, LHCB5, and LHCB6) as well as via the PSI complexes. At the same time, however, the LT/HL treatment induces an additional strong sNPQ in lettuce, which is largely retained after 1 h recovery in darkness, when canonical NPQ mechanisms are largely relaxed. This novel sNPQ is related to accumulation of SEP2, which is proposed to bind and quench the CP47 inner antenna of PSII core (Figure 10) while free CP43, accumulating due to the stalling of pre-D1 maturation, is protected by LPA3. At the same time, STN8 kinase dependent phosphorylation hampers the degradation of damaged D1 leading to accumulation of non-functional PSII centres. On the other hand, a strong accumulation of ELIP1.2 as an early event during the LT/HL treatment and subsequent 1 h recovery in the dark seems to be important to bind and store the Chls released from damaged and degraded pigment-protein complexes (Supplemental figure 4), thereby preventing the generation of singlet oxygen by free Chls.

It is important to emphasise that a physiologically relevant understanding of the regulation of the photosynthetic apparatus requires disentangling the roles of multiple and often condition-specific mechanisms acting simultaneously in a highly controlled manner. Nevertheless, the redox regulation is a unifying feature of a number of such mechanisms. It is therefore conceivable that excessive excitation stress triggers in the photosynthetic apparatus a wave of redox activation or inactivation of different enzymes and signalling cascades, leading to photoprotection of the photosynthetic apparatus and acclimation to the new conditions over a longer time scale. The hypotheses presented here would be a starting point for further in-depth studies on the regulation of photoprotective networks in plant chloroplasts and far beyond.

## Acknowledgements

The work was supported by the Jane and Aatos Erkko Foundation and the University of Turku Graduate School. Steffen Grebe, Minna Konert, Marjaana Rantala and Andrea Trotta are thanked for helpful discussions, and Mika Keränen and Virpi Paakkarinen for their expert technical advice.

## Data sharing and data availability

The proteomics raw-data supporting the findings of this work are available in the PRIDE Archive database (PXD055190). More detailed information is available on reasonable request from the corresponding authors.

## Supporting information

Supplemental files contain detailed methods and databases used to analyse the MS results.

