## Supplementary material for "Sustained non-photochemical quenching and regulation of PSII repair cycle during combined low temperature and high light stress in lettuce": Supplemental file 1_Detailed methods, supplemental figures and link to other files.docx

### Sustained non-photochemical quenching and photoprotection of PSII repair cycle during combined high light and low temperature stress in lettuce

Tapio Lempiäinen, Dorota Muth-Pawlak, Julia P. Vainonen, Eevi Rintamäki, Mikko Tikkanen, Eva-Mari Aro

#### Detailed methods

##### Thylakoid isolation

Leaves from GT/GL and LT/HL treated and recovered plants were ground in ice-cold grinding buffer (50 mM Hepes-NaOH pH 7.5, 330 mM sorbitol, 5 mM MgCl_2_, 0.05 % (w/v) BSA and 10 mM NaF) and filtered through Miracloth (Millipore). Chloroplasts were collected by centrifugation at 1350×g for 5 min at 4 °C and osmotically ruptured in ice-cold shock buffer (50 mM Hepes-NaOH pH 7.5, 5 mM sorbitol, 10 mM MgCl_2_ and 10 mM NaF). The released thylakoid membranes were collected by centrifugation at 1350×g for 5 min at 4 °C, washed twice with storage buffer (50 mM Hepes-NaOH pH 7.5, 100 mM sorbitol, 10 mM MgCl_2_ and 10 mM NaF), and finally resuspended in storage buffer.

##### Western blotting

Thylakoid proteins were solubilised in denaturation buffer (138 mM Tris-HCl pH 6.8, 6 M urea, 22.2 % (v/v) glycerol, 4.3 % (w/v) SDS) with 10 % (v/v) β-mercaptoethanol, and separated by sodium dodecyl sulphate polyacrylamide gel electrophoresis (SDS-PAGE) gel containing 12 % acrylamide and 6 M urea. The separated proteins were electroblotted onto a PVDF membrane (Millipore) using a transfer buffer (20% methanol, 40 mM glycine, 50 mM Tris, 5 mM SDS). The membranes were blocked with 5 % non-fat milk (Bio-Rad) in TBS (20 mM Tris pH 7.5, 150 mM NaCl), and the blocked membranes were incubated overnight with anti-P-Thr primary antibody (New England Biolabs) diluted in 1 % non-fat milk in TTBS (TBS with 0.05% Tween 20). The membranes were then washed with TTBS, and incubated with IR dye-labelled secondary antibody (IRDye® 800CW goat anti-rabbit IgG secondary antibody, Li-Cor) diluted in 1 % non-fat milk in TTBS for 1 h. The secondary antibody was detected using an Odyssey CLx imager (Li‑Cor).

##### Clear Native and 2D gel electrophoresis

Isolated thylakoids were suspended in ice-cold 25BTH20G buffer (25 mM BisTris-HCl pH 7.0, 20 % (v/v) glycerol and 0.25 mg/ml Pefabloc). Resuspended thylakoids were solubilised with an equal volume of 2 % β‑*D*-dodecyl maltoside (β‑DM) or 3 % digitonin in 25BTH20G buffer. β-DM samples were solubilised for 5 min on ice and digitonin samples were solubilised for 8 min at room temperature with gentle mixing. Insoluble material was removed by centrifugation at 16 000 *g* for 20 min at 4°C and deoxycholate was added to the supernatant to a final concentration of 10 %. Solubilised thylakoid protein complexes were separated on 5 % ‑ 12 % or 3 % - 12 % acrylamide gradient gels. Separated pigment-protein complexes were imaged using an office scanner and an Odyssey CLx imager (Li-Cor) with 685 nm excitation and 700 nm fluorescence cut-off. For separation of proteins in the second dimension, a gel slice cut from the native gel was incubated in the denaturation buffer with 5 % (v/v) β-mercaptoethanol for 30 min at RT and then loaded onto the top SDS-PAGE gel containing 12 % acrylamide and 6 M urea to separate individual proteins in each of the protein complexes.

##### Proteomics analysis

Equal amounts of isolated thylakoids, based on Chl content, were solubilised by mixing the thylakoids with a buffer (0,1M Tris-NaOH pH 8) containing 6 M urea and 0.1 % RapiGest (Waters). The concentrations of solubilised proteins were determined by the Bradford assay, and aliquots of solubilised proteins corresponding to 100 µg of protein were then subjected to reduction with DTT, and alkylation with IAA, followed by precipitation of the proteins in a cold acetone/ethanol mixture at -20°C overnight. The solubilised proteins were then digested with trypsin (1 µl 1 µg/µl, 1:100) in a buffer (5 % ACN, 0.05 M Tris-NaOH pH 8) for 4 h, after which a second batch of trypsin (1µ 1 µg/µl, 1:100) was added and the digestion was continued overnight. The peptide mixtures were desalted in parallel with Wet Sep-Pak 100 mg C18 96 (Waters) according to the manufacturer's protocol.

The peptide concentration of the desalted peptides was determined using a DeNovix DS-11+ spectrophotometer and equal amounts of peptides were injected for nLC-ESI-FAIMS-MS/MS analyses. Prior to injection, the samples were spiked with iRT synthetic peptides (Biognosys). The injected samples were first trapped on a pre-column and then separated on an analytical C18 column (75 μm x 15 cm, ReproSil-Pur 3 μm 120 Å C18-AQ, Dr. Maisch HPLC GmbH, Ammerbuch-Entringen, Germany) by a two-step, 110 min gradient from 5 to 26% solvent B over 70 min, followed by 26 to 49% B increase over 30 min. The mobile phase consisted of water with 0.1% formic acid (solvent A) or acetonitrile/water (80:20 (v/v)) with 0.1% formic acid (solvent B). The MS data were acquired automatically using Thermo Xcalibur 4.6 software (Thermo Fisher Scientific). FAIMS was operated at two compensation voltages (CV) values: -50 and -70 for both MS and MS/MS modes. The data-independent acquisition (DIA) method consisted of a survey scan of the mass range 395-1005 m/z with a maximum injection time of 50 ms and an automatic gain control (AGC) target set to 7e5 ions, followed by a series of tMS2 scans triggered by a set of variable isolation window for molecular ions. The isolation width of windows ranged from 15–110 m/z covering mass range of 395-1005 m/z for molecular ions (Supplemental table 1 contains the full list of isolation windows) with a maximum injection time of 52 ms and an AGC target of 1e6. All ions within the isolation windows were subjected to HCD fragmentation and the fragments were registered within 180-2000 m/z range (Supplemental table 1 contains the full list of isolation windows). All precursors within the isolation windows were fragmented with a normalised collision energy of 28 %. The spectra were registered at a resolution of 120 000 and 30 000 (at m/z 200) for full scan and fragment spectra, respectively.

##### Pigment analysis with HPLC-DAD

Pigments were extracted from isolated thylakoids at room temperature under dim light. Thylakoid samples equivalent to 20 µg Chl were extracted with 100 µl of acetone by continuous vortex mixing for 5 min. The samples were then centrifuged and the supernatants were collected. The pellet was re-extracted with 100 µl of acetone and the supernatants were pooled and filtered through 0.2 µm PTFE filters. The extracted pigments were separated on a reverse phase C18 column (LiChroCART 125-4, Hewlett Packard) attached to a Series 1100 HPLC instrument with a diode array and fluorescence detectors (Agilent Technologies). Buffer A was acetonitrile-methanol-Tris-HCl buffer 0.1 M pH 8.0 (72:8:3) and buffer B was methanol-hexane (4:1). A constant flow rate of 0.5 ml min^-1^ was used. Pigment separation was started with a 4 min isocratic run with 95% buffer A and 5% buffer B, followed by a 15 min linear gradient from 95% buffer A and 5% buffer B to 0% buffer A and 100% buffer B. The gradient was followed by a 26 min isocratic run with buffer 100% buffer, which was followed by a 15 min re-equilibration with 95% buffer A and 5% buffer B.

#### Supplemental figures

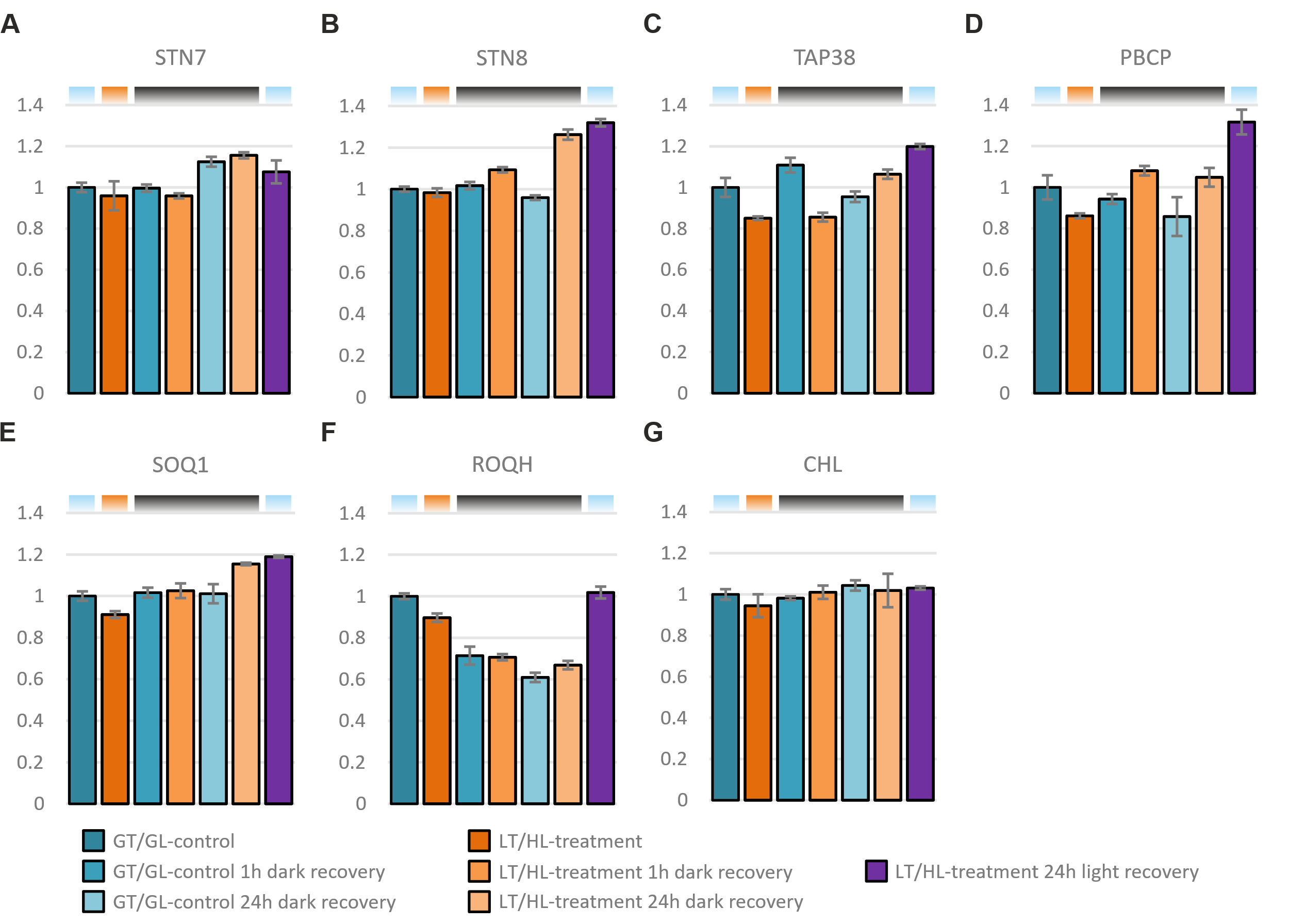
**Supplemental figure 1.** Effect of low temperature and high light treatment, and the subsequent recovery, on regulatory proteins of photosynthesis. **A)** State transition 7 (STN7) **B)** State transition 8 (STN8) **C)** Thylakoid-associated phosphatase 38 (TAP38) **D)** Photosystem II core phosphatase (PBCP) **E)** Suppressor of quenching 1 (SOQ1) **F)** Relaxation of qH (ROQH1) **G)** Chloroplastic lipocalin (CHL). Long day-grown lettuce plants were illuminated under 1500 µmol photons m−2 s−1 of white light at 13 °C for 4 h (LT/HL), while control plants were kept at growth conditions (23 °C and 140 µmol photons m−2 s−1 with 16 h photoperiod) (GT/GL), after which all plants were transferred to recover for 1 h and 24 h in darkness or for 24 h in long day growth conditions. Thylakoid membranes used in the analyses were isolated directly after the treatment and the recovery periods. Isolated thylakoids were solubilized with detergent and isolated proteins were digested with trypsin. Resulted peptide mixtures were analysed with nLC-ESI-FAIMS-MS/MS in DIA mode and protein abundances were determined with Spectronaut software. Protein abundances were normalised to the average of control plants in growth light. Error bars show standard deviations among technical replicates (n = 3). Coloured bars above the graphs represent the temperature and light conditions from which the thylakoids used in the analyses were isolated: light blue for GT/GL, orange for LT/HL and black for GT/darkness.

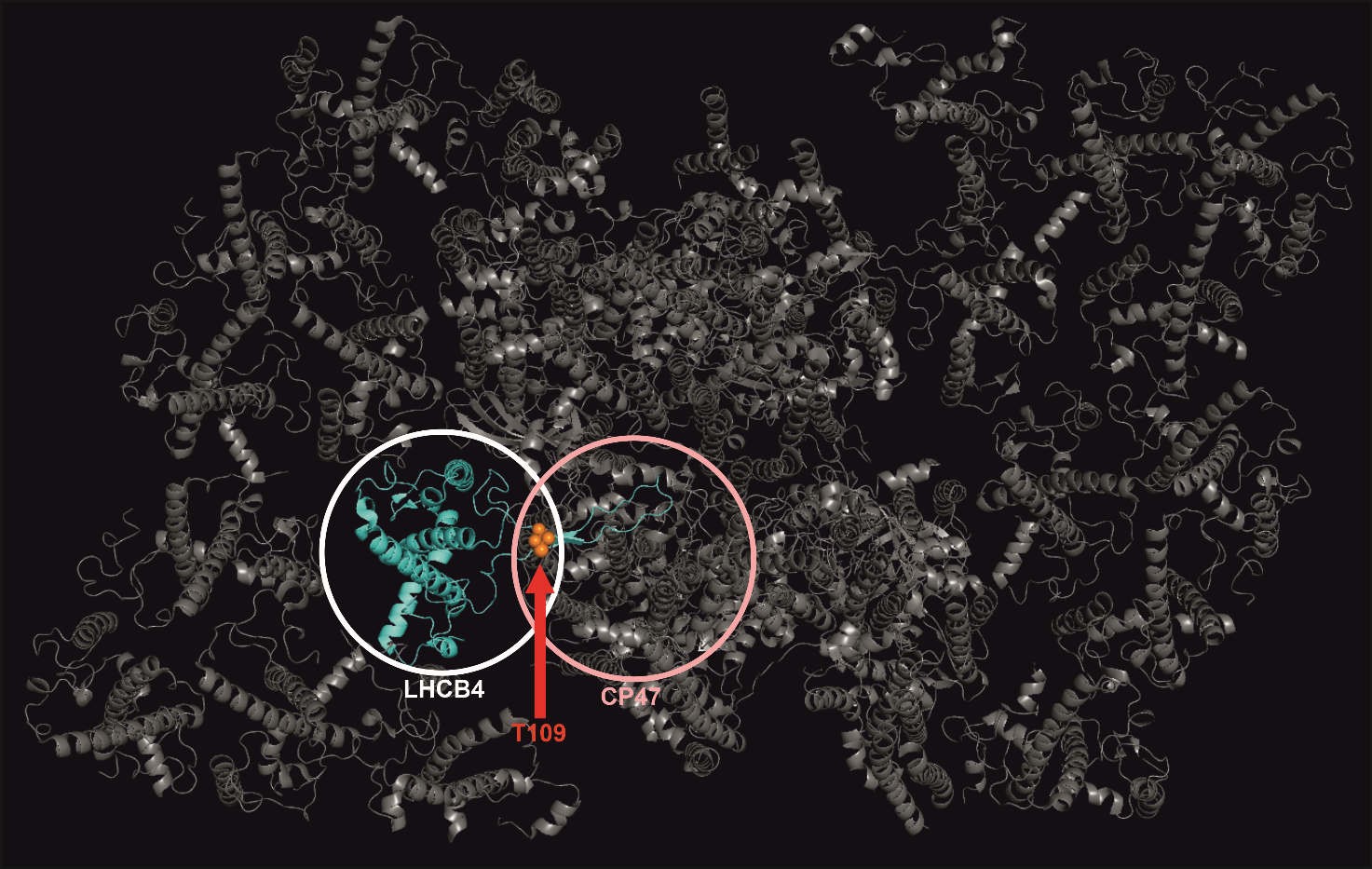

###### **Suppelemental figure 2.** LHCB4 and CP47 interaction in the PSII sc and the localisation of T109, which is highly phosphorylated in lettuce treater for 4 h at low temperature and high light treated (1500 µmol photons m^−2^ s^−1^ of white light at 13 °C). The figure was generated using PyMol software and it is based on the structure of Pea (*Pisum sativum*) PSII sc (Su et al. 2017).

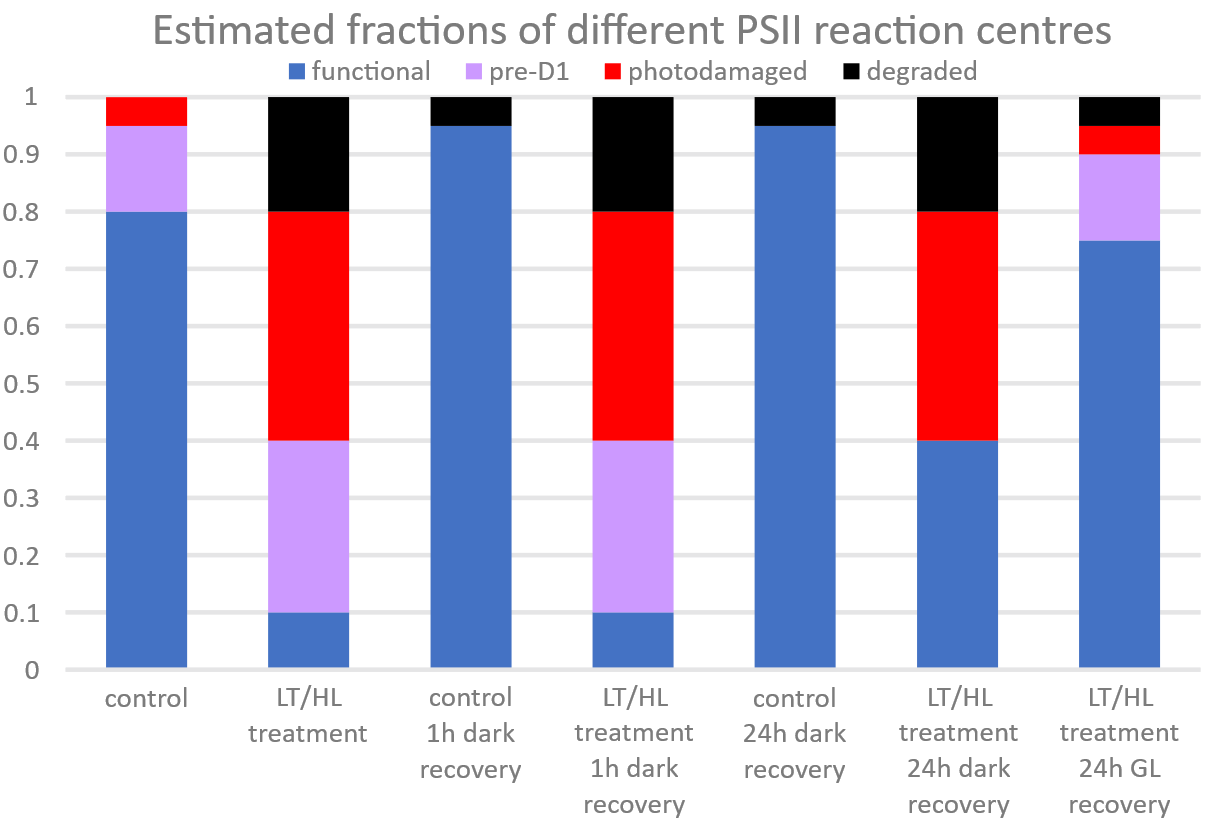
Supplemental figure 3. Estimation of different fractions of PSII during LT/HL-treatment and subsequent recovery.

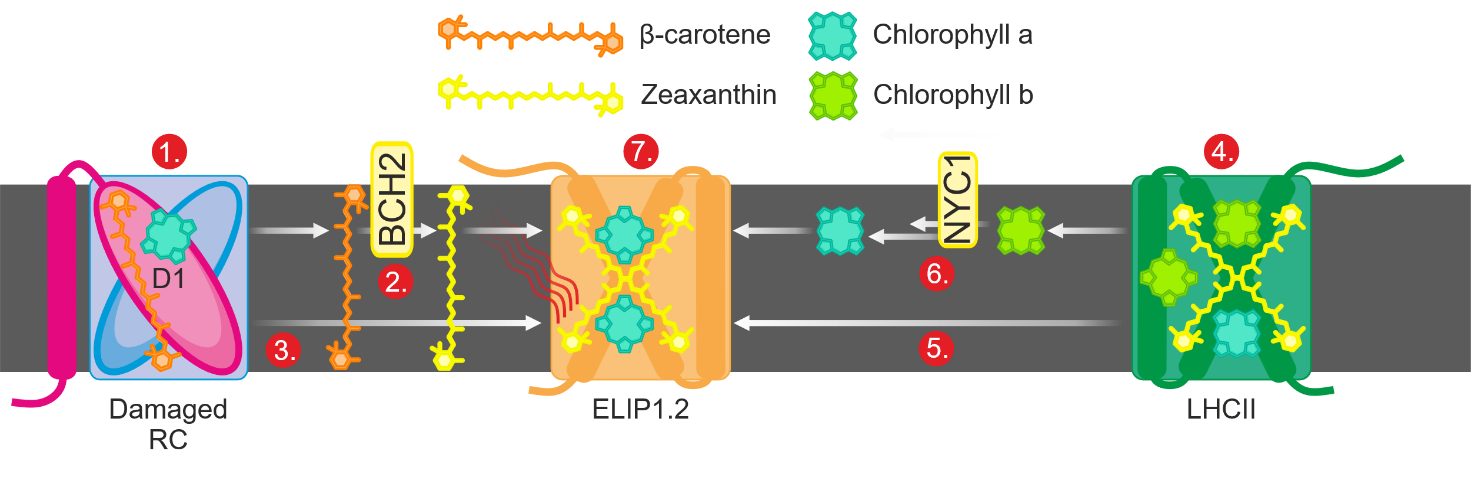

###### **Supplemental figure 4.** Proposed role for ELIP1.2 protein in storing the Chl from damaged and degraded PSII and LHCII in thylakoids of the LT/HL‑treated lettuce: **1.** Damaged PSII cores are degraded by FTSH protease and pigments are released from the complex. **2.** The released β‑carotene is hydrolysed by BCH2 (β-carotene hydroxylase) to zeaxanthin, which binds to the *de­‑novo* translated ELIP1.2. **3.** Chls released from degraded PSII cores are scavenged by ELIP1.2 when the Chl salvage pathway is not active enough to immediately incorporate the released Chl a into the PSII repair cycle. **4.** Degradation of LHCII releases bound Chl a and Chl b. **5.** Chl a released from LHCII is transferred to ELIP1.2. **6.** Reduction of Chl b to Chl a is initiated by NYC1 (chlorophyll b reductase) and the formed Chl a is bound to ELIP1.2. **7.** ELIP1.2 quenches the bound Chl a with zeaxanthin, making ELIP1.2 a safe pigment store until the bound Chl a can be reused or eventually degraded.
