## Supplementary material for "Sustained non-photochemical quenching and regulation of PSII repair cycle during combined low temperature and high light stress in lettuce": Supplemental file 2_Lettuce SwissProt sequences.docx

>sp|A0A2J6L8Y7|IF4E1_LACSA Eukaryotic translation initiation factor 4E-1 OS=Lactuca sativa OX=4236 GN=eIF4E PE=1 SV=1

MVEEIMKSEEQKLIDVNKHRGVRSDGEEDEQLEEGEIVGGDADTLSSSSSSRPGTAIAQH

PLEHSWTFWFDTPSAKSKQVAWGSSMRPIYTFSSVEEFWSLYNNIHRPSKLAQGADFYCF

KNKIEPKWEDPVCANGGKWTMTFTKAKSDTCWLYTLLAMIGEQFDHGDDICGAVVNVRAR

QEKIALWTKNAANESAQLSIGKQWKEFIDYNDTIGFIFHEDAKTLDRSAKNKYTV

>sp|D5J9U8|GAO_LACSA Germacrene A hydroxylase OS=Lactuca sativa OX=4236 GN=GAO1 PE=1 SV=1

MELSITTSIALATIVFFLYKLATRPKSTKKQLPEASRLPIIGHMHHLIGTMPHRGVMDLA

RKHGSLMHLQLGEVSTIVVSSPKWAKEILTTYDITFANRPETLTGEIIAYHNTDIVLAPY

GEYWRQLRKLCTLELLSVKKVKSFQSIREEECWNLVKEVKESGSGKPINLSESIFTMIAT

ILSRAAFGKGIKDQREFTEIVKEILRQTGGFDVADIFPSKKFLHHLSGKRARLTSIHKKL

DNLINNIVAEHHVSTSSKANETLLDVLLRLKDSAEFPLTADNVKAIILDMFGAGTDTSSA

TVEWAISELIRCPRAMEKVQAELRQALNGKEKIQEEDIQDLAYLNLVIRETLRLHPPLPL

VMPRECREPVNLAGYEIANKTKLIVNVFAINRDPEYWKDAEAFIPERFENNPNNIMGADY

EYLPFGAGRRMCPGAALGLANVQLPLANILYHFNWKLPNGASHDQLDMTESFGATVQRKT

ELLLVPSF

>sp|F8S1I0|C7BL2_LACSA Costunolide synthase OS=Lactuca sativa OX=4236 GN=CYP71BL2 PE=1 SV=1

MEPLTIVSLAVASFLLFAFWALSPKTSKNLPPGPPKLPIIGNIHQLKSPTPHRVLRNLAK

KYGPIMHLQLGQVSTVVVSTPRLAREIMKTNDISFADRPTTTTSQIFFYKAQDIGWAPYG

EYWRQMKKICTLELLSAKKVRSFSSIREEELRRISKVLESKAGTPVNFTEMTVEMVNNVI

CKATLGDSCKDQATLIEVLYDVLKTLSAFNLASYYPGLQFLNVILGKKAKWLKMQKQLDD

ILEDVLKEHRSKGRNKSDQEDLVDVLLRVKDTGGLDFTVTDEHVKAVVLDMLTAGTDTSS

ATLEWAMTELMRNPHMMKRAQEEVRSVVKGDTITETDLQSLHYLKLIVKETLRLHAPTPL

LVPRECRQACNVDGYDIPAKTKILVNAWACGTDPDSWKDAESFIPERFENCPINYMGADF

EFIPFGAGRRICPGLTFGLSMVEYPLANFLYHFDWKLPNGLKPHELDITEITGISTSLKH

QLKIVPILKS

>sp|P48706|RBL_LACSA Ribulose bisphosphate carboxylase large chain OS=Lactuca sativa OX=4236 GN=rbcL PE=3 SV=2

MSPQTETKASVGFKAGVKDYKLTYYTPEYETKDTDILAAFRVTPQPGVPPEEAGAAVAAE

SSTGTWTTVWTDGLTSLDRYKGRCYGIEPVPGEENQYIAYVAYPLDLFEEGSVTNMFTSI

VGNVFGFKALRALRLEDLRIPTAYVKTFQGPPHGIQVERDKLNKYGRPLLGCTIKPKLGL

SAKNYGRAVYECLRGGLDFTKDDENVNSQPFMRWRDRFLFCAEAIFKSQAETGEIKGHYL

NATAGTCEEMMKRAIFARELGVPIVMHDYLTGGFTANTSLAHYCRDNGLLLHIHRAMHAV

IDRQKNHGIHFRVLAKALRMSGGDHIHSGTVVGKLEGEREITLGFVDLLRDDFIEKDRSR

GIYFTQDWVSLPGVLPVASGGIHVWHMPALTEIFGDDSVLQFGGGTLGHPWGNAPGAVAN

RVALEACVQARNEGRDLATEGNEIIREATKWSPELAAACEVWKEIKFEFQAMDTLDQ

>sp|P69557|PSBA_LACSA Photosystem II protein D1 OS=Lactuca sativa OX=4236 GN=psbA PE=3 SV=2

MTAILERRESESLWGRFCNWITSTENRLYIGWFGVLMIPTLLTATSVFIIAFIAAPPVDI

DGIREPVSGSLLYGNNIISGAIIPTSAAIGLHFYPIWEAASVDEWLYNGGPYELIVLHFL

LGVACYMGREWELSFRLGMRPWIAVAYSAPVAAATAVFLIYPIGQGSFSDGMPLGISGTF

NFMIVFQAEHNILMHPFHMLGVAGVFGGSLFSAMHGSLVTSSLIRETTENESANEGYRFG

QEEETYNIVAAHGYFGRLIFQYASFNNSRSLHFFLAAWPVVGIWFTALGISTMAFNLNGF

NFNQSVVDSQGRVINTWADIINRANLGMEVMHERNAHNFPLDLAAIEAPSTNG

>sp|Q40251|VDE_LACSA Violaxanthin de-epoxidase, chloroplastic OS=Lactuca sativa OX=4236 GN=VDE1 PE=1 SV=1

MALSLHTVFLCKEEALNLYARSPCNERFHRSGQPPTNIIMMKIRSNNGYFNSFRLFTSYK

TSSFSDSSHCKDKSQICSIDTSFEEIQRFDLKRGMTLILEKQWRQFIQLAIVLVCTFVIV

PRVDAVDALKTCACLLKECRIELAKCIANPSCAANVACLQTCNNRPDETECQIKCGDLFE

NSVVDQFNECAVSRKKCVPRKSDVGEFPVPDRNAVVQNFNMKDFSGKWYITSGLNPTFDA

FDCQLHEFHMENDKLVGNLTWRIKTLDGGFFTRSAVQTFVQDPDLPGALYNHDNEFLHYQ

DDWYILSSQIENKPDDYIFVYYRGRNDAWDGYGGSVIYTRSPTLPESIIPNLQKAAKSVG

RDFNNFITTDNSCGPEPPLVERLEKTAEEGEKLLIKEAVEIEEEVEKEVEKVRDTEMTLF

QRLLEGFKELQQDEENFVRELSKEEKEILNELQMEATEVEKLFGRALPIRKLR

>sp|P0CC80|NU2C1_LACSA NAD(P)H-quinone oxidoreductase subunit 2 A, chloroplastic OS=Lactuca sativa OX=4236 GN=ndhB1 PE=3 SV=1

MIWHVQNENFILDSTRIFMKAFHLLLFDGSLIFPECILIFGLILLLMIDSTSDQKDIPWL

YFISSTSLVMSITSLLFRWREEPMISFSGNFQTNNFNEIFQFLILLCSTLCIPLSVEYIE

CTEMAITEFLLFVLTATIGGMFLCGANDLITIFVAPECFSLCSYLLSGYTKKDVRSNEAT

MKYLLMGGASSSILVHGFSWLYGSSGGEIELQEIVNGLINTQMYNSPGISIALIFITVGI

GFKLSPAPSHQWTPDVYEGSPTPVVAFLSVTSKVAASASATRIFDIPFYFSSNEWHLLLE

ILAILSMILGNLIAITQTSMKRMLAYSSIGQIGYVIIGIIVGDSNDGYASMITYMLFYIS

MNLGTFACIVLFGLRTGTENIRDYAGLYTKDPFLALSLALCLLSLGGLPPLAGFFGKLYL

FWCGWQAGLYFLVLIGLLTSVVSIYYYLKIIKLLMTGRNQEITPHVRNYRRSPLRSNNSI

ELSMIVCVIASTIPGISMNPIIAIAQDTLF

>sp|P0CC81|NU2C2_LACSA NAD(P)H-quinone oxidoreductase subunit 2 B, chloroplastic OS=Lactuca sativa OX=4236 GN=ndhB2 PE=3 SV=1

MIWHVQNENFILDSTRIFMKAFHLLLFDGSLIFPECILIFGLILLLMIDSTSDQKDIPWL

YFISSTSLVMSITSLLFRWREEPMISFSGNFQTNNFNEIFQFLILLCSTLCIPLSVEYIE

CTEMAITEFLLFVLTATIGGMFLCGANDLITIFVAPECFSLCSYLLSGYTKKDVRSNEAT

MKYLLMGGASSSILVHGFSWLYGSSGGEIELQEIVNGLINTQMYNSPGISIALIFITVGI

GFKLSPAPSHQWTPDVYEGSPTPVVAFLSVTSKVAASASATRIFDIPFYFSSNEWHLLLE

ILAILSMILGNLIAITQTSMKRMLAYSSIGQIGYVIIGIIVGDSNDGYASMITYMLFYIS

MNLGTFACIVLFGLRTGTENIRDYAGLYTKDPFLALSLALCLLSLGGLPPLAGFFGKLYL

FWCGWQAGLYFLVLIGLLTSVVSIYYYLKIIKLLMTGRNQEITPHVRNYRRSPLRSNNSI

ELSMIVCVIASTIPGISMNPIIAIAQDTLF

>sp|P23712|GLNA_LACSA Glutamine synthetase OS=Lactuca sativa OX=4236 PE=2 SV=2

MALLSDLVNLDLSSITDKIIAEYIWIGGSGMDLRSKARTLSGPVSDPSELPKWNYDGSST

GQAPGEDSEVIIYPQAIFKDPFRRGNHILVMCDAYTPAGEPIPTNKRAAAAKIFSNPEVE

KEVTWYGIEQEYTLLQKDTNWPLGWPLGGFPGPQGPYYCGIGADKAFGRDIVDAHYKACL

YAGVNISGINGEVMPGQWEFQVGPSVGIAAADQIWVARYILERITEIYGVVVSFDPKPIP

GDWNGAGAHTNYSTKTMREEGGYEVIKKAIEKLGLRHKEHIAAYGEGNERRLTGRHETAD

INTFLWGVANRGASIRVGRDTEKEGKGYFEDRRPASNMDPYVVTSMIAETTILWDNKS

>sp|P48493|TPIS_LACSA Triosephosphate isomerase, cytosolic (Fragment) OS=Lactuca sativa OX=4236 PE=2 SV=1

IQVAAQNCWVKKGGAFTGEVSAEMLANLGVPWVILGHSERRALLNETNEFVGDKVAYALS

QGLKVIACVGETLEQREAGTTMEVVAAQTKAIADKISSWDNVVLAYEPVWAIGTGKVASP

AQAQEVHAGLRKWFCDNVSAEVSASTRIIYGGSVSGSNCKELGGQTDVDGFLVGGASLKP

EFIDIIKAAEVKKSA

>sp|Q1KXG6|NU4C_LACSA NAD(P)H-quinone oxidoreductase chain 4, chloroplastic OS=Lactuca sativa OX=4236 GN=ndhD PE=3 SV=1

MNKFPWLTIIVVLPIFAGSFILFLPHKGNRVIRWYTICICMLELLLTTYAFCYHFQLDDP

LIQLVEDYKWINFFDFRWKLGIDGLSIGPVLLTGFITTLATLAAWPVTRDSRLFHFLMLA

MYSGQIGSFSSRDLLLFFIMWELELIPVYLLLSMWGGKKRLYSATKFILYTAGGSIFLLM

GVLGVGLSGSNEPTLNFETSVNQSYPVALEIIFYIGFFIAFAVKLPILPLHTWLPDTHGE

AHYSTCMLLAGILLKMGAYGLIRINMELLPHAHSIFSPWLMIVGTIQIIYAASTSPGQRN

LKKRIAYSSVSHMGFILIGIASITDTGLNGAILQIISHGFIGAALFFLAGTSYDRIRLVY

LDQMGGVAIPMPKIFTMFSSFSMASLALPGMSGFVAEVIVFLGIITSQKYLLMPKIAITF

VMAIGMILTPIYSLSMLRQMFYGYKLFNTPNAYVFDSGPRELFVSISIFLPVIGIGMYPD

FVLSLSVDKVEGILSNYFYR

>sp|Q32539|NU5C_LACSA NAD(P)H-quinone oxidoreductase subunit 5, chloroplastic OS=Lactuca sativa OX=4236 GN=ndhF PE=3 SV=2

MEQTYQYAWIIPFLPLPVPMLIGLGLLLFPTATKSLRRMWSFQSVLLLSIVMIFSMNLSI

QQINSSYVYQYVWSWIINNDFSLEFGYLIDPLTSIMLILITTVGIMVLIYSDNYMSHDHG

YLRFFAYMSFFSTSMLGLVTSSNLIQIYIFWELVGICSYLLIGFWFTRPVAAKACQKAFV

TNRVGDFGLLLGILGFYWITGSFEFRDLFQIFNNLISNNEVNFLFVTLCAILLFAGAIAK

SAQFPLHVWLPDAMEGPTPISALIHAATMVAAGIFLVARLMPLFIVIPHIMNFISLIGII

TVFLGATLALAQKDIKRGLAYSTMSQLGYMMLALGMGSYRSALFHLITHAYSKALLFLGS

GSVIHSMETLVGYCPKKSQNMVLMGGLTKHLPITKNSFLLGTLSLCGIPPLACFWSKDEI

LNDSWLYSPIFGIIAWSTAGLTAFYMCRIYLLTFEGHLNVHFQNYSGKRNTPFYSISLWG

KEGSKISNKNFSLVTLLKMKKNPRPSFFSNNKVYKIDENVRNMIQPFLSIPHFGNTKTYS

YPYESDNTMLFPILILILFTLFVGFLGIPFNQDVDILSKWLNPSINLLHQNSNNSIDWYE

FSKDAFFSVSIASFGIFIAFFLYKPVYSSFQNLEFLNTFVKMGPNRIFYDKIKNSIYDWS

YNRGYIDAFYGRFLTAGIRKLANFAHFFDRRIIDAIPNGVGLMSFFGAEVIKSVGGGRIS

SYLFFYFSYVAIFLLIYYFFNV

>sp|Q332S1|NDHH_LACSA NAD(P)H-quinone oxidoreductase subunit H, chloroplastic OS=Lactuca sativa OX=4236 GN=ndhH PE=3 SV=1

MTGPATRKDLMIVNMGPHHPSMHGVLRLIVTLDGEDVIDCEPILGYLHRGMEKIAENRTI

IQYLPYVTRWDYLATMFTEAITVNAPEQLGNIQVPKRASYIRVIMLELSRIASHLLWLGP

FMADIGAQTPFFYIFRERELIYDLFEAATGMRMMHNFFRIGGIAADLPHGWIDKCLDFCD

YFLTGIAEYQKLITRNPIFLERVEGVGIIGGEEAINWGLSGPMLRASGIQWDLRKVDHYE

CYDEFDWEVQWQNEGDSLARYLVRISEMTESIKIIQQALEGIPGGPYENLEIRRFDRVKD

TVWNEFDYRFISKKPSPTFELSKQELYARVEAPKGELGIFLIGDKGVFPWRYKIRPPGFI

NLQILPQLVKRMKLADIMTILGSIDIIMGEVDR

>sp|Q332S2|NU1C_LACSA NAD(P)H-quinone oxidoreductase subunit 1, chloroplastic OS=Lactuca sativa OX=4236 GN=ndhA PE=3 SV=1

MIIDTTEVQAINSFSILESLKEVYGIIWMLIPIFTLVLGITIGVLVIVWLEREISAGIQQ

RIGPEYAGPLGILQALADGTKLLFKENLLPSRGDTRLFSIGPSIAVISILLSYLVIPFSY

HLVLADLSIGVFLWIAISSIAPVGLLMSGYGSNNKYSFLGGLRAAAQSISYEIPLTLCVL

SISLLSNSSSTVDIVEAQSKYGFWGWNLWRQPIGFLVFLISSLAECERLPFDLPEAEEEL

VAGYQTEYSGIKFGLFYVASYLNLLVSSLFVTVLYLGGWNLSIPYIFVPEVFEITKRGRV

FGTIIGIFITLAKTYLFLFIPIATRWTLPRLRMDQLLNLGWKFLLPISLGNLLLTTCSQL

ISL

>sp|Q332S3|NDHI_LACSA NAD(P)H-quinone oxidoreductase subunit I, chloroplastic OS=Lactuca sativa OX=4236 GN=ndhI PE=3 SV=1

MFPMVTEFMNYGQQTVRAARYIGQGFMITLSHANRLPVTIQYPYEKLITSERFRGRIHFE

FDKCIACEVCVRVCPIDLPVVDWKLETDIRKKRLLNYSIDFGICIFCGNCVEYCPTNCLS

MTEEYELSTYDRHELNYNQIALGRLPMSVIDDYTIRTIFNLPEIKT

>sp|Q332S4|NU6C_LACSA NAD(P)H-quinone oxidoreductase subunit 6, chloroplastic OS=Lactuca sativa OX=4236 GN=ndhG PE=3 SV=1

MDLPGLIHDFLLVFLGLGLILGGLGVVLLPNPIYSAFSLGLVFVCISLFYILSNSHFVAA

AQLLIYVGAINVLIIFAVMFINGSEYSKDFHLWTVGDGITSVVCTSLFVSLITTIPDTSW

YGIIWTTKANQIIEQDLISNSQQIGIHLSTDFFLPFEFISIILLVALIGAIAVARQ

>sp|Q332S5|NU4LC_LACSA NAD(P)H-quinone oxidoreductase subunit 4L, chloroplastic OS=Lactuca sativa OX=4236 GN=ndhE PE=3 SV=1

MMLEHVLVLSAYLFSVGLYGLITSRNMVRALMCLELILNAVNLNFVTFSDFFDSRQLKGA

IFSIFVIAIAAAEAAIGLAIVSSIYRNRKSTRINQSNLLNK

>sp|Q332S6|PSAC_LACSA Photosystem I iron-sulfur center OS=Lactuca sativa OX=4236 GN=psaC PE=3 SV=1

MSHSVKIYDTCIGCTQCVRACPTDVLEMIPWDGCKAKQIASAPRTEDCVGCKRCESACPT

DFLSVRVYLWHETTRSMGIAY

>sp|Q332U5|RPOA_LACSA DNA-directed RNA polymerase subunit alpha OS=Lactuca sativa OX=4236 GN=rpoA PE=3 SV=1

MVREKITVSTRTLQWKCVESAADSKRLLYGRFILSPLMKGQADTIGIAMRRALLGEIEGT

CITRAKSEKISHEYSTIMGIQESVHEILMNLKEIVLRSNLYGTCEASICVRGPGYVTAQD

IILPPYVEILDNTQHIASLTEPIELVIGLQIEKNRGYLIKAPNTFQDGSYPIDPVFMPVR

NANHSIHSYENGNKEILFLEIWTNGSLTPKEALYEASRNLIDLLIPFLHTKEENLNLEGN

QHMVPLPPFTFYDKLAKLTKNKKKMALKSIFIDQSELPPRIYNCLKRSNIYTLLDLLNNS

QEDLMKMEHFRIEDVKQILGILEKNFVIDLPKNKF

>sp|Q332U6|PETD_LACSA Cytochrome b6-f complex subunit 4 OS=Lactuca sativa OX=4236 GN=petD PE=3 SV=2

MGVTKKPDLNDPVLRAKLAKGMGHNYYGEPAWPNDLLYIFPVVILGTIACNVGLAVLEPS

MIGEPADPFATPLEILPEWYFFPVFQILRTVPNKLLGVLLMVSVPAGLLTVPFLENVNKF

QNPFRRPVATTVFLIGTAVALWLGIGATLPIDKSLTLGLF

>sp|Q332U8|PSBH_LACSA Photosystem II reaction center protein H OS=Lactuca sativa OX=4236 GN=psbH PE=3 SV=1

MATQTVENGARSGPRRTTVGDLLKPLNSEYGKVAPGWGTTPLMGVAMALFAVFLSIILEI

YNSSVLLDGISMN

>sp|Q332V2|CLPP_LACSA ATP-dependent Clp protease proteolytic subunit OS=Lactuca sativa OX=4236 GN=clpP PE=3 SV=1

MPIGVPKVPFRSPGEEDASWVDIYNRLYRERLLFLGQEVDSEISNQLIGLMIYLSIEDDT

KDLYLFINSPGGWVIPGVALYDTMQFVQPDVHTICMGSAASMGSFILVGGEITKRLAFPH

ARVMIHQPAGSFSEVATGEFILEVGELLKLRETLTRVYVQRTGKPLWVVSEDMERDVFMS

ATEAQAYGIVDLVAVE

>sp|Q332W1|PSBE_LACSA Cytochrome b559 subunit alpha OS=Lactuca sativa OX=4236 GN=psbE PE=3 SV=1

MSGSTGERSFADIITSIRYWVIHSITIPSLFIAGWLFVSTGLAYDVFGSPRPNEYFTENR

QGIPLITGRFDSLEQLDEFSRSF

>sp|Q332W2|PSBF_LACSA Cytochrome b559 subunit beta OS=Lactuca sativa OX=4236 GN=psbF PE=3 SV=1

MTIDRTYPIFTVRWLTVHGLAVPTVSFLGSISAMQFIQR

>sp|Q332W9|ACCD_LACSA Acetyl-coenzyme A carboxylase carboxyl transferase subunit beta, chloroplastic OS=Lactuca sativa OX=4236 GN=accD PE=3 SV=2

MKRWWFNSMLFKKEFEHRCRLSKSMGSLGPIENASESKDPNRNDTDKNIQGWGGHDNYSN

VDLFFGVKDIRNFFSDDTFLVKDSNGDSYSIYFDIENHIFEIANDHPFCSELESSFYRNS

SDLNNGSKSKNPDHDRYMDDTQYTWNNHINSCIDSYLQYQICIDNYIVSGNDNSSNNNSS

NENSSNENSSNENSSNENSSNDYISSSISSQSENSSQNEDITTSDQTIPESSTHMGVTQQ

YRHLWVQCENCYGLNYKKFFKSKMHLCEQCGYHLKMSSSDRIELLIDPGTWEPMDEDMVS

LDPIEFHSEEEPYKNRIDSYQRNTGLTEAVQTGRGQLNGITVAIGVMDFQFMGGSMGSVV

GEKITRLIEYATKEFLPLIIVCASGGARMQEGSVSLMQMAKISSALYDYQSNKKLFYVPI

LTSPTTGGVTASFGMLGDIIIAEPNAYIAFAGKRVIEQTLNKTVPEGSQAAEYLFQKGLF

DLIVPRNPLKSVLSELFQLHTFFPLNQN

>sp|Q332X1|ATPB_LACSA ATP synthase subunit beta, chloroplastic OS=Lactuca sativa OX=4236 GN=atpB PE=3 SV=1

MRMNPTTSGSGVTTLDKKTLGRIAQIIGPVLDVAFPPGKMPNIYNALVVKGRDTAGQPIN

VTCEVQQLLGNNRVRAVAMSATDGLTRGMDVIDTGAPLSVPVGGATLGRIFNVLGEPVDN

LGPVDTSTTFPIHRSAPAFIQLDTKLSIFETGIKVVDLLAPYRRGGKIGLFGGAGVGKTV

LIMELINNIAKAHGGVSVFGGVGERTREGNDLYMEMKESGVINEKNIPESKVALVYGQMN

EPPGARMRVGLTALTMAEYFRDVNEQDVLLFIDNIFRFVQAGSEVSALLGRMPSAVGYQP

TLSTEMGSLQERITSTKEGSITSIQAVYVPADDLTDPAPATTFAHLDATTVLSRGLAAKG

IYPAVDPLDSTSTMLQPRIVGEEHYDTAQEVKQTLQRYKELQDIIAILGLDELSEEDRLT

VARARKIERFLSQPFFVAEVFTGSPGKYVGLAETIRGFQLILSGELDGLPEQAFYLVGNI

DEATAKAMNLEMESNLKK

>sp|Q332X3|NU3C_LACSA NAD(P)H-quinone oxidoreductase subunit 3, chloroplastic OS=Lactuca sativa OX=4236 GN=ndhC PE=3 SV=1

MFLLYEYDIFWAFLIISSLIPILVFFISGFLAPISKGPEKLSSYESGIEPIGDAWLQFRI

RYYMFALVFVVFDVETVFLYPWAMSFDVLGVSVFVEALIFVLILIVGLVYAWRKGALEWS

>sp|Q332X4|NDHK_LACSA NAD(P)H-quinone oxidoreductase subunit K, chloroplastic OS=Lactuca sativa OX=4236 GN=ndhK PE=3 SV=1

MNSIEFPLFDRTTQNSVISTTLNDLSNWSRLSSLWPLLYGTSCCFIEFASLIGSRFDFDR

YGLVPRSSPRQADLILTAGTVTMKMAPSLVRLYEQMPEPKYVIAMGACTITGGMFSTDSY

STVRGVDKLIPVDVYLPGCPPKPEAIIDAITKLRKKISREIYPDRTMSQRENRCFTTNHK

FQVGHSIHTGNYDQGFLYQPTSTSEIPPETFFKYKSSVSSPELVN

>sp|Q332X5|NDHJ_LACSA NAD(P)H-quinone oxidoreductase subunit J, chloroplastic OS=Lactuca sativa OX=4236 GN=ndhJ PE=3 SV=1

MQGHLSAWLVKHGLIHRSLGFDYQGIETLQIKPGDWHSIAVILYVYGYNYLRSQCAYDVA

PGGLLASVYHLTRIEYGADQPEEVCIKVFAPRRDPRIPSVFWVWKSVDFQERESYDMLGI

SYDNHPRLKRILMPESWIGWPLRKDYIAPNFYEIQDAH

>sp|Q332X6|RR4_LACSA 30S ribosomal protein S4, chloroplastic OS=Lactuca sativa OX=4236 GN=rps4 PE=3 SV=1

MSRYRGPRFKKIRRLGALPGLTNKRPRAGSDLRNQSRSGKKSQYRIRLEEKQKLRFHYGL

TERQLLKYVRIAGKAKGSTGQVLLQLLEMRLDNILFRLGMAPTIPGARQLVNHRHILVNG

RIVDIPSYRCKPRDTIAARDEQKSKVLIQNSLDSSPHEELPNHLTLQPFQYKGLVNQIID

SKWVGLKINELLVVEYYSRQT

>sp|Q332X8|PSAA_LACSA Photosystem I P700 chlorophyll a apoprotein A1 OS=Lactuca sativa OX=4236 GN=psaA PE=3 SV=1

MIIRSPEPEVKILVDRDHIKTSFEEWARPGHFSRTIAKGPETTTWIWNLHADAHDFDSHT

SDLEEISRKVFSAHFGQLSIIFLWLSGMYFHGARFSNYEAWLSDPTHIRPSAQVVWPIVG

QEILNGDVGGGFRGIQITSGFFQIWRASGITSELQLYCTAIGGLVFAALMLFAGWFHYHK

AAPKLAWFQDVESMLNHHLAGLLGLGSLSWAGHQVHVSLPINQFLNAGVDPKEIPLPHEF

ILNRDLLAQLYPSFAEGATPFFTLNWSKYADFLTFRGGLDPVTGGLWLTDTAHHHLAIAI

LFLIAGHMYRTNWGIGHGLKDILEAHKGPFTGQGHKGLYEILTTSWHAQLSLNLAMLGSL

TIVVAHHMYAMPPYPYLATDYGTQLSLFTHHMWIGGFLIVGAAAHAAIFMVRDYDPTTRY

NDLLDRVLRHRDAIISHLNWACIFLGFHSFGLYIHNDTMSALGRPQDMFSDTAIQLQPVF

AQWIQNTHALAPGATAPGATASTSLTWGGGDLVAVGGKVALLPIPLGTADFLVHHIHAFT

IHVTVLILLKGVLFARSSRLIPDKANLGFRFPCDGPGRGGTCQVSAWDHVFLGLFWMYNS

ISVVIFHFSWKMQSDVWGSISDQGVVTHITGGNFAQSSITINGWLRDFLWAQASQVIQSY

GSSLSAYGLFFLGAHFVWAFSLMFLFSGRGYWQELIESIVWAHNKLKVAPATQPRALSIV

QGRAVGVTHYLLGGIATTWAFFLARIIAVG

>sp|Q332X9|PSAB_LACSA Photosystem I P700 chlorophyll a apoprotein A2 OS=Lactuca sativa OX=4236 GN=psaB PE=3 SV=1

MALRFPRFSQGLAQDPTTRRIWFGIATAHDFESHDDITEERLYQNIFASHFGQLAIIFLW

TSGNLFHVAWQGNFESWVQDPLHVRPIAHAIWDPHFGQPAVEAFTRGGALGPVNIAYSGV

YQWWYTIGLRTNEDLYTGALFLLLISAISLIAGWLHLQPKWKPSVSWFKNAESRLNHHLS

GLFGVSSLAWTGHLVHVAIPASRGEYVRWNNFLDVLPHPQGLGPLFTGQWNLYAQNPDSG

SHLFGTSQGAGTAILTLLGGFHPQTQSLWLTDMAHHHLAIAFLFLIAGHMYRTNFGIGHS

MKDLLDAHIPPGGRLGRGHKGLYDTINNSLHFQLGLALASLGVITSLVAQHMYSLPAYAF

IAQDFTTQAALYTHHQYIAGFIMTGAFAHGAIFFIRDYNPEQNEDNVLARMLEHKEAIIS

HLSWASLFLGFHTLGLYVHNDVMLAFGTPEKQILIEPIFAQWIQSAHGKTSYGFDILLSS

TNGPAFNAGRSIWLPGWLNAINENSNSLFLTIGPGDFLVHHAIALGLHTTTLILVKGALD

ARGSKLMPDKKDFGYSFPCDGPGRGGTCDISAWDAFYLAVFWMLNTIGWVTFYWHWKHIT

LWQGNVSQFNESSTYLMGWLRDYLWLNSSQLINGYNPFGMNSLSVWAWMFLFGHLVWATG

FMFLISWRGYWQELIETLAWAHERTPLANLIRWRDKPVALSIVQARLVGLAHFSVGYIFT

YAAFLIASTSGKFG

>sp|Q332Y2|PSBC_LACSA Photosystem II CP43 reaction center protein OS=Lactuca sativa OX=4236 GN=psbC PE=3 SV=1

MKTLYSLRRFYPVETLFNGTLALAGRDQETTGFAWWAGNARLINLSGKLLGAHVAHAGLI

VFWAGAMNLFEVAHFVPEKPMYEQGLILLPHLATLGWGVGPGGEVIDTFPYFVSGVLHLI

SSAVLGFGGIYHALLGPETLEESFPFFGYVWKDRNKMTTILGIHLILLGIGAFLLVFKAL

YFGGVYDTWAPGGGDVRKITNLTLSPSIIFGYLLKSPFGGEGWIVSVDDLEDIIGGHVWL

GSICILGGIWHILTKPFAWARRALVWSGEAYLSYSLAAISVFGFIACCFVWFNNTAYPSE

FYGPTGPEASQAQAFTFLVRDQRLGANVGSAQGPTGLGKYLMRSPTGEVIFGGETMRFWD

LRAPWLEPLRGPNGLDLSRLKKDIQPWQERRSAEYMTHAPLGSLNSVGGVATEINAVNYV

SPRSWLATSHFVLGFFFFVGHLWHAGRARAAAAGFEKGIDRDFEPVLSMTPLN

>sp|Q332Y4|ATPA_LACSA ATP synthase subunit alpha, chloroplastic OS=Lactuca sativa OX=4236 GN=atpA PE=3 SV=1

MVTIQADEISNIIRERIEQYNREVKIVNTGTVLQVGDGIARIHGLDEVMAGELVEFEEGT

IGIALNLESTNVGVVLMGDGLLIQEGSSVKATGRIAQIPVSEAYLGRVINALAKPIDGRG

EISSSEYRLIESPAPGIISRRSVYEPLQTGLIAIDSMIPIGRGQRELIIGDRQTGKTAVA

TDTILNQQGKNVICVYVAIGQKASSVAQVVTNFQERGAMEYTIVVAETADSPATLQYLAP

YTGAALAEYFMYRERHTSIIYDDPSKQAQAYRQMSLLLRRPPGREAYPGDVFYLHSRLLE

RAAKLSSLLGEGSMTALPIVETQSGDVSAYIPTNVISITDGQIFLSADLFNAGIRPAINV

GISVSRVGSAAQIKAMKQVAGKLKLELAQFAELEAFAQFASDLDKATQNQLARGQRLREL

LKQSQSAPLGVEEQVLTIYTGTNGYLDSLEIGQVRKFLVELRTYLKTNKPQFQEIISSTK

TFTEEAEAILKEAIKEQRERFILQEQAA

>sp|Q56P05|PSBD_LACSA Photosystem II D2 protein OS=Lactuca sativa OX=4236 GN=psbD PE=3 SV=1

MTIALGKVTKDENDLFDIMDDWLRRDRFVFVGWSGLLLFPCAYFAVGGWFTGTTFVTSWY

THGLASSYLEGCNFLTAAVSTPANSLAHSLLLLWGPEAQGDFTRWCQLGGLWTFVALHGA

FGLIGFMLRQFELARSVQLRPYNAIAFSGPIAVFVSVFLIYPLGQSGWFFAPSFGVAAIF

RFILFFQGFHNWTLNPFHMMGVAGVLGAALLCAIHGATVENTLFEDGDGANTFRAFNPTQ

AEETYSMVTANRFWSQIFGVAFSNKRWLHFFMLFVPVTGLWMSALGVVGLALNLRAYDFV

SQEIRAAEDPEFETFYTKNILLNEGIRAWMAAQDQPHENLIFPEEVLPRGNAL

>sp|Q56P08|ATPH_LACSA ATP synthase subunit c, chloroplastic OS=Lactuca sativa OX=4236 GN=atpH PE=3 SV=1

MNPLISAASVIAAGLAVGLASIGPGVGQGTAAGQAVEGIARQPEAEGKIRGTLLLSLAFM

EALTIYGLVVALALLFANPFV

>sp|Q56P11|RPOC2_LACSA DNA-directed RNA polymerase subunit beta'' OS=Lactuca sativa OX=4236 GN=rpoC2 PE=3 SV=2

MEVLMAERPTQVFHNKVIDGTAMKRLISRFIDHYGIGYTSHILDQVKTLGFRQATAASIS

LGIDDLLTIPSKRWLVQDAEQQSFILEKHHHYGNVHAVEKLRQSIEIWYATSEYLRQEMN

PNFRMTDPFNPVHIMSFSGARGNASQVHQLVGMRGLMSDPQGQMIDLPIQSNLREGLSLT

EYIISCYGARKGVVDTAIRTSDAGYLTRRLVEVVQHIVVRRTDCGTVRGISVSPRNGMMT

DRIFIQTLIGRVLADDIYIGSRCIATRNQDIGVGLVSRFITFRAQPISIRTPFTCRSTSW

ICQLCYGRSPAHDDLVELGEAVGIIAGQSIGEPGTQLTLRTFHTGGVFTGGTAEHVRAPS

NGKIKFNEDLVHPTRTRHGHPAFLCSRDLYVTIESEDIIHNVCIPPKSFLLVQNDQYVES

EQVIAEIRARTSTLNLKEKVRKHIYSDSEGEMHWNTDVYHAPEFTYGNIHLLPKTSHLWI

LLGEPWRYSLGPCSIHKDQDQMNAYSLSVKPRYIANPSVTNNQVRHKFFSSYFSGKNQKG

DRIPDCSELNRMTCTDHSNLRYPAILDGNSDLLAKRRRNRFIIPLESIQEGENQLIPSSG

ISMEIPRNGILRRNSILAYFDDPRYIRKSSGLTKYETRELNSIVNEENLIEYRGVKVFWP

KYQKEVNPFFFIPVEVHILSESSSIMVRHNSIIGADTQITFNRRSRVGGLVRVKKKAEKM

KLIIFSGDIHFPGKTNKAFRLIPPGGGKPNSKEYKKLKNWLYIQRMKLSRYEKKYFVLVQ

PVVPYKKTDGINLGRLFPPDLLQESDNLQLRVVNYILYYDPILEIWDTSIQLVRTSLVLN

WDQDKKIEKACASFVEIRTNGLLRYFLRIDLAKSPISYTGKRNDLSGSGLISENGSDRAN

VNPFSSIYSYSKSRIKESLNPNQGTIHTLLNRNKESQSLIILSSSNCFRIGPFNDVKSPN

VIKESIKKNPLIPIRNSLGPLGTGFPIYNFDLFSHLITHNQILVTNYLQLDNFKQIFQIL

KYYLLDENGQIYNPYSCSNIILNPFHLNWYFLHYNYCEETSPIVSLGQFLCENVCIAKKG

PHLKSGQVLIVQVDSVVIRSAKPYLATPGATVHGHYGEILYEGDTLVTFIYEKSRSGDIT

QGLPKVEQVLEVRSIDSISMNLEKRIEGWNKSITRILGIPWAFLIGAELTIVQSRISLVN

KVQKVYRSQGVQIHNRHIEIIVRQITSKVLVSEDEMSNVFSPGELIGLLRAERMGRALEE

AICYQAVLLGITRASMNTQSFISEASFQETARVLAKAALLGRIDWLKGLKENVVLGGMIP

VGSGFKTPSSEPNNIPNNIAFELQKKNLLEGEMKDILFYHRKLFDSCLSNNFHDTQEQSF

F

>sp|Q56P12|RPOC1_LACSA DNA-directed RNA polymerase subunit beta' OS=Lactuca sativa OX=4236 GN=rpoC1 PE=3 SV=2

MIDRYTHQQLRIGLVSPQQISTWSKKILPNGEIVGEVTKPYTFHYKTNKPEKDGLFCERI

FGPIKSGICACGNYRVIGDEKEDPQFCEQCGVEFVDSRIRRYQMGYIKLAYPVMHVWYLK

RLPSYIVTLLDKPLNELEDLVYCNFYFARPIDKKPTFLRLRGLLEYEIQPWKYRIPIFFT

TRSFDTFRNREMSTGGGSIRQQLANLDLRIIIDYSLVEWKELEEEEPTGNEWEDRKVGRR

KDFLLRRMELAKHFIRTNIEPKWMVLRLLPVLPPELRPIYHIDEDKLVTSDINEIYRRII

YRNNTLTDLLTTSIATPEELIISQEKLLQEAVDALLDNGICGQPMRDDHNRVYKSLSDVI

EGKEGRVRETLLGKRVDYSGRSVIVVGPSLSLHRCGLPREIAIELFQAFVIRDLIRKHLA

SNIGVAKSQIRKKKPIVWEILQEILDDHPVLLNRAPTLHRLGIQAFLPVLVEGRAICLHP

LVCKGFNADFDGDQMAVHVPLSLEAQAEARLLMFSHMNLLSPTIGDPISAPTQDMLSGLY

VLTSGNRRGICVNRYNPCNRRNYQNEDNNYKYTKKKEPFFCNPYDAIGAYRQKRINLGSP

LWLRWRLDQRVIAAREVPIEIHYESVGTYYEIYGHYLIVRSIKKEILYIYIRTTLGHISL

YREIEEAIQGFWQGCCNSMLPTGIRVSPG

>sp|Q56P13|RPOB_LACSA DNA-directed RNA polymerase subunit beta OS=Lactuca sativa OX=4236 GN=rpoB PE=3 SV=2

MSTIPGFNQIQFEGFCRFIDQGLTEELSKFPKIEDTNQEIDFELFLERYQLVEPSIKERD

AVYESLTYSSELYVSARLIWKNDRRRYIQEQTILIGKIPLMTSLGAFIVNGIYRIVINQI

LQSPGIYYQSELNDNGISVYTGTIISDWGGRLELEIDRKTRIWVRVSRQQKLSILVLLSA

MGLNIREILENVCYPELFLSFLNDKKQIGSKENAILEFYQQFACVEGDPVFSESLSKDLQ

KKFFQQRCELGGIGRRNMNRRLNLDIPQNNTFLLPRDILAAADRLIRIKFGMGTLDDMNH

LQNKRIRSVADLLQEQFGLALVRLENMARGNIYAALKHNWTPTPQNLVNSTPLTDTYKVF

FRLHPLSQVLDRTNPLTQIVHGRKLSYLGPGGLTARTATFPIRDIHPSHYGRICPIDTSE

GINVGLIGSLAIHARIGRWGSLESPFYKISERSKGARMLYLSPGRDEYYMVAAGNSLALN

QGIQEEQVVPARYRQEFLTIAWEQVHLRSIFSFQYFSIGASLIPFIEHNDANRALMSSNM

QRQAVPLSQSEKCIVGTGLEGQAALDSGALAIAEHEGEIIYTDTDKILLSGNGDTLRIPL

VMYQRSNKNTCMHQKPQVQRGKCIKKGQILAYGAATVGGELALGKNVLVAYMPWEGYNFE

DAVLISERLVYEDIYTSFHIRKYEIQINQGSERVTNEIPHLEVHLLRNLDKNGIVMLGSW

VETGDILVGKLTPQMVKESSYAPEDRLLRTILGMRVYTSKETCLKLPIGGRGRVIDVRWV

QSSKTDETEKTESIRVYILQKREIKVGDKVAGRHGNKGIISKILPRQDMPYLQDGRPVDM

VFNPLGVPSRMNVGQIFESSLGLAGGLLDRHYRIAPFDERYEQEASRKLVFSELYEASKQ

TVNPWIFEPESPGKSRIFDGRTGDPFEQPVIIGKPYILKLIHQVDDKIHGRSSGRYSRLT

QQPLKGRAKKGGQRVGEMEVWALEGFGVAYILQEMLTYKSDHIRARQEVLGTIIFGGRIP

TPEDAPESFRLFVRELRSLALELNHFLVSEKTFQLNRKEA

>sp|Q9SEC2|MSRA_LACSA Peptide methionine sulfoxide reductase OS=Lactuca sativa OX=4236 PE=2 SV=1

MFLLRTTTATTTPASLPLPLLSISSHLSLSKPSSFPVTSTKPLFTLRHSSSTPKIMSWLG

RLGXGTRTPADASMDQSSIAQGPDDDIPAPGQQFAQFGAGCFWGVELAFQRVPGVSKTEV

GYTQGFLHNPTYNDICSGTTNHSEVVRVQYDPKACSFDSLLDCFWERHDPTTLNRQGNDV

GTQYRSGIYFYTPEQEKAAIEAKERHQKKLNRTVVTEILPAKKFYRAEEYHQQYLAKGGR

FGFRQSTEKGCNDPIRCYG

>sp|A0A2J6KL39|NLTP_LACSA Non-specific lipid-transfer protein Lac s 1 OS=Lactuca sativa OX=4236 GN=LSAT_1X82001 PE=1 SV=1

MARMAMMILCVVLTCMVVATPYTEAAISCGQVTANLAGCLNYLRNGGAVPPACCNGVRSL

NSAAKSTPDRKTACNCLKNASKSVSGIKAANAAGLPGKCGVNIPYQISPNTDCSKVQ

>sp|A0A3B7TLI7|BLR40_BRELC RxLR effector protein BLR40 OS=Bremia lactucae OX=4779 GN=BLR40 PE=2 SV=1

MLLSRAISVVALLACICCGVHTQDSKADLGTLRTTDSAIITSQRRLRTSVDLVDNEERFR

WPFQQFFKDRWHRKQIKTYFRDQKDNVSEGLVEQLIARHGLKNVEKVLSEVKFPLAVQIS

IRKILVNYKGKQAFTRPHLTPADTL

>sp|P00290|PLAS_LACSA Plastocyanin OS=Lactuca sativa OX=4236 GN=PETE PE=1 SV=1

AEVLLGSSDGGLVFEPSTFSVASGEKIVFKNNAGFPHNVVFDEDEIPAGVDASKISMSEE

DLLNAPGETYAVTLTEKGTYSFYCAPHQGAGMVGKVTVN

>sp|P0DI59|CASP1_LACSA Casparian strip membrane protein 1 OS=Lactuca sativa OX=4236 PE=2 SV=1

MKAGPLQLGVVPPANRAIAILDFFLRPIAIVGTLASAIAMATTNQTLPFFSQFIRFRAKF

NDLPSFTFFVVASSIVSAYLILSLGFSILHIAKSNLVNSRVLLLLLDTAAMGLLMAGSAA

ATAIVQLAHKGNNKVNWFAICQQYNSFCKRVSGSLIGSYAGVVVLILLILLSGVALSRR

>sp|Q1KXH4|TI214_LACSA Protein TIC 214 OS=Lactuca sativa OX=4236 GN=TIC214 PE=3 SV=1

MILKSFLLGNLVSLCMKIINSVVVVGLYYGFLTTFSIGPSYLFLLRAHIMEEGEEGTEKK

VSATTGFITGQLIMFISIYYAPLHLALGRPHTITVLALPYLLFHFFCNTHKHFFDYGSTN

RNSMRNLSIQCVFLNNLIFQLFNHFILPSSMLARLVNIFMFQCNNKIIFVTSSFVGWIIG

HIILMKSIGLLVVWIRKNRSIRKYIRSNKSFVSELANSMSIAGILNILLFVTCVYYLGRM

PLPIINKKLNNLEKMDQAKLKNNTPLSYIYKNQEDLYLEILGKKDKEKSFSLFEKPILTF

FFDYNRWNRPLRYIRKINKNLSVRKETSQYFFYTCQSDGKKRISFTYPPSLSTFGEMIAR

RISLSTLEKLSADALYTEWLYTNKEKNNNLNNEFINRIEALETVFLSINILDTKTRLCNV

ETEKKKNCLVNKKNYLVKMDDPFLTGMYRGRINKLFSSSIINQISIENYEKTYELNKIHY

NLLPYPNSREYEHKIDPNSPDYEQKIEKLEKIQAQIDLNNRFIFLWTTIIAKLKGQKNSS

RINEIGKKPPRWSYKLINELHQNYKKRRKEQGIIQGLRHQLRTRKYKHIYFLNRSTRTLE

TLKQSNLNNSNMDKKFNNKDLGFISYLEEPDFRRSLIKGSMRAQRRKLVIWGPYQGNPHS

PLFLEKKQDFPFPISDLIKLFLNIKDRLGKKSEFEILNKQSPPKRNNQEDVMEFWETIPH

GHKTRGILLLAQSTFRKYIKLPLLIIAKNIVRILLRQSPEWDEDFQDWNREIYLKCSSNG

LQFSKTKFPKNWLRGGFQIKILYPFHLKPWHRSKLRLYDSDRDLKQQEDFDSCFLTVLGM

ETEHPFGPPRKTPSFFEPIFKDIDYKVEIRKLNFRVRRIFKKIKKKEAKAFFFIKQKIKE

LLKGNKIPLFLPREIYESSETQTEKDSIISNQIIHESLSQIRSTGWTNYSQAEEEMKHRI

DRRKTIRNEIEIMKKNKINNAESSQKILKILKIKNIELLVKFFIEKIYIEIFLCIINMRR

IPLQLFIQSTKKIIDNDKYINNNETNQERINKTKQNKIDFILSMTIKRAFDNLRNSKRKS

DIFFDLSYLSQKYVFFKLSQTQIINFNKLRSILQYNGPSFCLKTEIKDFFGRQGIFHSEL

RHNKLPNYGMNPWKNWLRGHYQYDLSQITSSRLIPQKWRNRINQCQTSQNKDLNKWYSSE

KDQLLDSKKKQNSKVYLLPNKEDNLKKNYRYDLLSYKYIHSETKKNYYIYRSSLETNNNQ

ENSTRNKEKFFSILKNIPSKNYLGKSDIIYMEKNKDRKFLYKINKNIKVEPNKDQIKDKN

KDQKKDKIHNNGLFYLPIDSNLEINYKKVFFDWMGMNEKILILNCLISNPKVFFFPKFVR

LYHKYKEKPWFIPSKLLLFNLNITSNFSENQNINGKQEEYFLKQEEYFLKQEEDFLKQEE

DFLKQEEDFLKQEEDFLKQEEDFFKFRPSNSKQYFELNNQNNIEEYFLESTEKLKIFLKG

DFPLQLRWAGRVNQLNQKIMNNIQIYGLLLSLINVRKITISYIQRKEMDLGIMSRNLNLT

QLMKTGILILEPARLSLKNDGQFFMYQIISISLVHKSKYQSNQRYQKQENVAKNIDKKNS

DLLVPEKILSSRRRRELRILISFNLNLKNNTGVDRNTLVYNENKIKKMESIFG

>sp|Q1KXK8|PSBL_LACSA Photosystem II reaction center protein L OS=Lactuca sativa OX=4236 GN=psbL PE=3 SV=1

MTQSNPNEQNVELNRTSLYWGLLLIFVLAVLFSNYFFN

>sp|Q1KXM2|YCF3_LACSA Photosystem I assembly protein Ycf3 OS=Lactuca sativa OX=4236 GN=ycf3 PE=3 SV=1

MPRSRINGNFIDKTFSIVANILLRIIPTTSGEKEAFTYYRDGMSAQSEGNYAEALQNYYE

AMRLEIDPYDRSYILYNIGLIHTSNGEHTKALEYYFRALERNPFLPQAFNNMAVICHYRG

EQAIRQGDSEIAEAWFDQAAEYWKQAIALTPGNYIEAHNWLKITRRFE

>sp|Q332R5|RK2_LACSA 50S ribosomal protein L2, chloroplastic OS=Lactuca sativa OX=4236 GN=rpl2-A PE=3 SV=1

MAIHLYKTSTPSTRNGAVDSKVKSNPRNNLIYGQHHCGKGRNARGIITAGHRGGGHKRLY

RKIDFRRNEKDIYGRIVTIEYDPNRNAYICLIHYRDGEKRYILHPRGAIIGDTIVSGTEV

PIKMGNALPLTDMPLGTAIHNIEITLGKGGQLARAAGAVAKLIAKEGKSATLKLPSGEVR

LISKNCSATVGQVGNVGVNQKSLGRAGSKRWLGKRPVVRGVVMNPVDHPHGGGEGRAPIG

RKQPTTPWGYPALGKRSRKRNKYSDNLILRRRSK

>sp|Q332R6|RK23_LACSA 50S ribosomal protein L23, chloroplastic OS=Lactuca sativa OX=4236 GN=rpl23-A PE=3 SV=1

MDGIRYAVFTDKSIQLLGKNQYTSNVESGSTRTEIKHWVELFFGVKVIAMNSHRLRGKAR

RMGPIMGQTMHYRRMIITLQPGYSIPPLRKKRT

>sp|Q332R7|YCF2_LACSA Protein Ycf2 OS=Lactuca sativa OX=4236 GN=ycf2-A PE=3 SV=1

MTGHEFKSWILELREILREIKNSHYFLDSWTQFNSVGSFIHIFFHQERFIKLFDSRIWSI

LLSHNSQGSTSNRYFTIKGVILFGVAVLIYRINNRNMVERKNLYLIGLLPIPMNSIGPRN

DTLEESVGSSNINRLIVSLLYLPKGKKIYESSFLNPKESTWVLPITKKCSMPESNWGSRW

WRDWIGKKRDSSCKISNETVAGIEILFKEKDLKYLEFFFVYYRDDPIRKDHDWELFDRLS

LRKRQNRINLNSGPLFEILVKHWICYLMSAFREKIPIEVEGFFKQQGAGSTIKSNDIEHV

SHLFSRNKSAISLQNCAQFHMWQFRQDLFVSWGKNPPESDLLRNVSRENLIWLDNVWLVN

KDRFFRKVRNVSSNIQYDSTRSSFVQVRDSSQLKGSSDQSRDHFDSISNEDSEYHTLINQ

REIQQLKERSILWDPSFLQTEGTEIESNRFPKCLSGYSSMSRLFTEREKQMINHLLPEEI

EEFLGNPTRSVRSFFSDRWSEFHLGSNPTERSTRDQKLLKKQQDLSFLRRSENKEMVNLF

KIITYLQNTVSIHPISSDSGCDMVPKDEPDMDSSNKISFLNKNPFFDLFHLFHDRNRGGY

TLHHDFESEERFQELADLFTLSITEPDLVYHKRFAFSIDSYGLDPKQFLNGVFNSRYEWK

TTSLLVLLVLLPIFYEENESFYRRIRKKRVRISCGNDLEEPKPKIVVFASNNIMEAANQY

RLIRNLIQIQHSTHRYIRNVLNRFFLMNRSDRNFKYGIQRDQIGKDTLNHRTLMKYMINQ

HLSNLKKSQKRWFDPLIFFSRTKRSMNRDPDAYRYKWSTGSKNFQEHFVSEQKSRFQVVF

DRLRINQYSIDWSEVIDKKDLSKPLRFFLSKLLLFLSNSLPFLFVSFGNIPIHRSEIYIY

ELKGPNDPQFLESIGLQIVHLKKLKPFLLDDHETCQKSKFLINGGTISPFLFNKIPKWMI

DSFHTRNNRRKSFDNTDSYFSMIFHDQYNWLNPVKSFHRSSLRSSFYKANQLRFLNNPHH

FCFYCNKRFPFYVEKARINNYDFTYGQFLNILFIRNKIFSLCVGKKKHAFWGRDTISAIE

SQVSNIFIPKAFPQSGDETYNLYKSFHFPSRSNPFVRRAIYSIADISGTPLTEGQIVNFE

RTYCQPLSDMNLSDSEGKNLYQYLNFNSNMGLIHTPCYEKYLPSEKRKKRSLCLKKCVEK

GQMYRTFQRDSAYSTLSKWNLFQTYMPWFLTSTGYRYLKFLFLDTFSDLLPILSSSQKFV

SIFHDIMHGSNISWRILQKKFCLPQRNLISEISSKCLHNLLLSEEMIHRNNESPLISTHL

TNVREFLYAILFLLLVAAYLACTRLLFVFGASSELQTEFEKVKSLMIPSSMIELRKLLDR

YPTSEPNSFWFLKQLGDSLGGNMLLGGGPAYRVKSIRSKKKYLNINLIDIIDLISIIPNP

INRITFSRNTRHLSHTSKEIYSLIRKRKNVNGDWIDDKIESWVANSDSIDDEKREFLVQF

STLTTEKRIDQILLSLTHSDHFSKNDSGYQMIEQPGAIYLRYLVDIHKKYLMNYEFNTSS

LAERRIFLAHYQTITYSQTSCGANSLHFPSHGKPFSLRLALSLSRGTLVIGSIGTGRSYL

VKYLAKNSYLPFITVFLNKSLDNKSQGFDNIDVDASDDSDASDDIDASDDILDMELELLT

SMNALTMDMMPEDEDLLYITLQFELAKAMSPCIIWIPNIHDLDVNESNYLSPGLLVNLLS

RDYETRNILVIASTHIPQKVDPALIAPNKLNTCIKIRRLLIPQQRKHFFTLSYTRGFHLE

KKMFHTNGFGSITMGSNARDLVALTNEALSISITQNKSIIDTNTIRSALHRQIWDLRSQV

RSVQDHGILFYKIGRAVAQNVLLSNCPIDPISIYMKKKSCNEVDYYLYNWYFELGTSMKK

LTILLYLLSCSAGSVTQDLWSLPGPDEKNGITPYGLVENDSGLVRGLLEVEGALVGSSRT

CSQFDKDRVTLLLRPEPRNPLDMMQNGSCSILDQRFLYEKDESEFEEGDERQQIEEDLFN

HIVWAPRIWRPWGFLFDCIERPNELGFPYWSRSFRGKRIVYDEEDELQENDSEFLQNGTV

QYQTRDISSKEQGLFRISQFIWDPADPLFFLFKAQPFVSVFSHRELFADEEMSKGLLTPQ

KNRPTSLYKRWFIKKTQEKHFELLINRQRWLRTNRSLSNGSFRSNTLSESYQYLSNLFLS

NGTLLDQMTKALLRKRWLFPDEMQIGFMEQDKDFPFLSQKDMWP

>sp|Q332R9|RR7_LACSA 30S ribosomal protein S7, chloroplastic OS=Lactuca sativa OX=4236 GN=rps7-A PE=3 SV=1

MSRRGTAEEKTAKSDPIYRNRLVNMLVNRILKHGKKSLAYQIIYRAVKKIQQKTETNPLS

VLRQAIHGVTPGIAVKARRVGGSTQQVPIEIGSTQGKALAIRWLLAASRKRPGRNMAFKL

SSELVDAAKGSGDAIRKREETHKMAESNRAFAHFR

>sp|Q332S0|RR15_LACSA 30S ribosomal protein S15, chloroplastic OS=Lactuca sativa OX=4236 GN=rps15 PE=3 SV=1

MVKNSFISIIFQEQEENKENRGSVEFQVVSFTNKIRKLTSHLELHKKDYLSQRGLRKILG

KRQRLLAYLSKKNRARYKELIGQLNIRERKTR

>sp|Q332S8|CCSA_LACSA Cytochrome c biogenesis protein CcsA OS=Lactuca sativa OX=4236 GN=ccsA PE=3 SV=1

MIFSTLEHIFTHISFSIVSIVIIIHLITLLGNEIIKPYDSSEKGMIVTFLCLTGLLITRW

IYSGHFPLSDLYESLIFLSWSFSLIHIVPYFKIRKNYLTEITASSTIFTQGFATSGLLTE

IRKPTILVPALQSEWLIMHVSMMILSYAALLCGSLLSVALLVITFRKIFYSYKSNNFLKL

NESFSFGEIQYKNERNNILKKNYFLSAKNYYKAQLIQQLDYWSYRVISLGFIFLTIGILS

GAVWANEAWGSYWSWDPKETWAFITWIVFAIYLHIRTNKNFQGANSAIVATLGFLIIWIC

YFGVNLLGIGLHSYGSFTLTSS

>sp|Q332S9|RK32_LACSA 50S ribosomal protein L32, chloroplastic OS=Lactuca sativa OX=4236 GN=rpl32 PE=3 SV=1

MAVPKKRTSISKKRIRKNIWKRKGYWAALKALSLGKSLSTGNSKSFFVRQTNKS

>sp|Q332T6|RR19_LACSA 30S ribosomal protein S19, chloroplastic OS=Lactuca sativa OX=4236 GN=rps19 PE=3 SV=1

MTRSLKKNPFVANNLLKKINKLNTKAEKEIIITWSRASTIIPIMVGHTIAIHNGKEHLPI

YITDRMVGHKLGEFAPTLNFRGHAKSDNRSRR

>sp|Q332T7|RK22_LACSA 50S ribosomal protein L22, chloroplastic OS=Lactuca sativa OX=4236 GN=rpl22 PE=3 SV=1

MLNKRTTEVYALGQHISMSAHKARRVIDQIRGRSYEETLMILELMPYRACYPIFKLVYSA

AANASFNMGSNEVNLVISKAEVNEGTIVKRLKPRARGRSFAIQKPTCHITIVMKDISLDE

YIDTDSITWSQKPKSKKKHTTMSYYDMYSNGGTWDKK

>sp|Q332T8|RR3_LACSA 30S ribosomal protein S3, chloroplastic OS=Lactuca sativa OX=4236 GN=rps3 PE=3 SV=1

MGQKINPIGFRLGTTQGHHSLWFAQPKNYSEGLQEDKKIRTYIQNYVQKNMKTSSGVEGI

ARIEIQKRIDLIQIIIYMGFPKILIESRPRGIEELQMNLQKEFHSVNRKLNIAITRIEKP

YGNPNILAEFIAGQLKNRVSFRKAMKKAIELTEQADTKGIQVQIAGRIDGKEIARVEWIR

EGRVPLQTIRAKIDYCCYTVRTIYGVLGIKIWIFIDGE

>sp|Q332T9|RK16_LACSA 50S ribosomal protein L16, chloroplastic OS=Lactuca sativa OX=4236 GN=rpl16 PE=3 SV=1

MLSPKRTRFRKQHRGRMKGISYRGNAICFGKYALQALEPAWITSRQIEAGRRAMTRNARR

GGKIWVRIFPDKPVTVRPAETRMGSGKGSPEYWVAVVKPGRILYEMGGVTENIARRAISI

AASKMPIRAQFIISG

>sp|Q332U0|RK14_LACSA 50S ribosomal protein L14, chloroplastic OS=Lactuca sativa OX=4236 GN=rpl14 PE=3 SV=1

MIQPQTHLNVADNSGARELMCIRIIGASNRRYAHIGDVIVAVIKDAVPNMPLERSEVVRA

VIVRTCKELKRDNGMIIRYDDNAAVVIDQEGNPKGTRVFGAIARELRQFNFTKIVSLAPE

VL

>sp|Q332U1|RR8_LACSA 30S ribosomal protein S8, chloroplastic OS=Lactuca sativa OX=4236 GN=rps8 PE=3 SV=1

MGSDTIADIITSIRNADMYRKSVVRVASTNISQSIVKILLREGFIENVRKHRENNKSFLV

LTLRHRRNRKRTYRNLLNLKRISRPGLRIYSNYQRIPRILGGMGIVILSTSQGIMTDREA

RLERIGGEILCYIW

>sp|Q332U2|IF1C_LACSA Translation initiation factor IF-1, chloroplastic OS=Lactuca sativa OX=4236 GN=infA PE=3 SV=1

MKEQKWIHEGLITESLPNGMFRVRLDNEDMILGYVSGKIRRSFIRILPGDRVKIEVSRYD

STRGRIIYRLRNKDSKD

>sp|Q332U3|RK36_LACSA 50S ribosomal protein L36, chloroplastic OS=Lactuca sativa OX=4236 GN=rpl36 PE=3 SV=1

MKIRASVRKICEKCRLIRRRGRIRVICSNPRHKQRQG

>sp|Q332U4|RR11_LACSA 30S ribosomal protein S11, chloroplastic OS=Lactuca sativa OX=4236 GN=rps11 PE=3 SV=1

MAKAIPKKGSRGRIGSRKSTRKIPKGVIHIQASFNNTIVTVTDVRGRVVSWSSAGTSGFR

GTKRGTPFAAQTAAGHAIRAVVDQGMQRAEVMIKGPGLGRDAALRAIRRSGILLTFVRDV

TPMPHNGCRPPKKRRV

>sp|Q332U7|CYB6_LACSA Cytochrome b6 OS=Lactuca sativa OX=4236 GN=petB PE=3 SV=1

MSKVYDWFEERLEIQAIADDITSKYVPPHVNIFYCLGGITLTCFLVQVATGFAMTFYYRP

TVTDAFASVQYIMTEANFGWLIRSVHRWSASMMVLMMILHVFRVYLTGGFKKPRELTWVT

GVVLGVLTASFGVTGYSLPRDQIGYWAVKIVTGVPEAIPVIGSPLVELLRGSASVGQSTL

TRFYSLHTFVLPLLTAVFMLMHFPMIRKQGISGPL

>sp|Q332U9|PSBN_LACSA Protein PsbN OS=Lactuca sativa OX=4236 GN=psbN PE=3 SV=1

METATLVAIFISGLLVSFTGYALYTAFGQPSQQLRDPFEEHGD

>sp|Q332V0|PSBT_LACSA Photosystem II reaction center protein T OS=Lactuca sativa OX=4236 GN=psbT PE=3 SV=1

MEALVYTFLLVSTLGIIFFAIFFREPPKVPTKK

>sp|Q332V1|PSBB_LACSA Photosystem II CP47 reaction center protein OS=Lactuca sativa OX=4236 GN=psbB PE=3 SV=1

MGLPWYRVHTVVLNDPGRLLSVHIMHTALVAGWAGSMALYELAVFDPSDPVLDPMWRQGM

FVIPFMTRLGITNSWGGWSITGGTITNPGIWSYEGVAGAHIVFSGLCFLAAIWHWVYWDL

EIFSDERTGKPSLDLPKIFGIHLFLAGVACFGFGAFHVTGLYGPGIWVSDPYGLTGKVQA

VNPSWGVEGFDPFVPGGIASHHIAAGTLGILAGLFHLSVRPPQRLYKGLRMGNIETVLSS

SIAAVFFAAFVVAGTMWYGSATTPIELFGPTRYQWDQGYFQQEIYRRVSAGLAENQSLSE

AWSKIPEKLAFYDYIGNNPAKGGLFRAGSMDNGDGIAVGWLGHPIFRDKEGRELFVRRMP

TFFETFPVVLVDGDGIVRADVPFRRAESKYSVEQVGVTVEFYGGELNGVSYSDPVTVKKY

ARRAQLGEIFELDRATLKSDGVFRSSPRGWFTFGHASFALLFFFGHIWHGARTLFRDVFA

GIDPDLDAQVEFGAFQKLGDPTTRRQIG

>sp|Q332V3|RR12_LACSA 30S ribosomal protein S12, chloroplastic OS=Lactuca sativa OX=4236 GN=rps12-A PE=3 SV=1

MPTIKQLIRNTRQPIRNVTKSPALRGCPQRRGTCTRVYTITPKKPNSALRKVARVRLTSG

FEITAYIPGIGHNSQEHSVVLVRGGRVKDLPGVRYHIVRGTLDAVGVKDRQQGRSSAL

>sp|Q332V5|RK20_LACSA 50S ribosomal protein L20, chloroplastic OS=Lactuca sativa OX=4236 GN=rpl20 PE=3 SV=1

MTRIRRGYIARRRRTKIRLFASSFRGAHSRLTRTITQQKIRALVSAHRDRDKQKINFRRL

WITRINAAIRERGVCYSYSRLINGLYKRQLLLNRKILAQIAISNRNCLYMISNEIIKEVG

WKESTG

>sp|Q332V6|RR18_LACSA 30S ribosomal protein S18, chloroplastic OS=Lactuca sativa OX=4236 GN=rps18 PE=3 SV=1

MDKSKRTFLKSKRSFRRRLPPIQSGDRIDYKNMSLISRFISEQGKILSRRVNRLTLKQQR

LITIAIKQARILSLLPFLNNEKQFERTESTTRTPSLRARKR

>sp|Q332V7|RK33_LACSA 50S ribosomal protein L33, chloroplastic OS=Lactuca sativa OX=4236 GN=rpl33 PE=3 SV=1

MAKGKDVRIPVLLECTACVRNGVNVNKASTGISRYITQKNRHNTPNRLELRKFCPYCYKH

TIHGEVKK

>sp|Q332V8|PSAJ_LACSA Photosystem I reaction center subunit IX OS=Lactuca sativa OX=4236 GN=psaJ PE=3 SV=1

MRDLKTYLSVAPVLSTLWFGSLAGLLIEINRFFPDALTFPFFSF

>sp|Q332V9|PETG_LACSA Cytochrome b6-f complex subunit 5 OS=Lactuca sativa OX=4236 GN=petG PE=3 SV=1

MIEVFLFGIVLGLIPITLAGLFVTAYLQYRRGDQLDL

>sp|Q332W0|PETL_LACSA Cytochrome b6-f complex subunit 6 OS=Lactuca sativa OX=4236 GN=petL PE=3 SV=1

MLTITSYFGFLLTALTITSALFIGLSKIRLI

>sp|Q332W4|PSBJ_LACSA Photosystem II reaction center protein J OS=Lactuca sativa OX=4236 GN=psbJ PE=3 SV=1

MADTTGRIPLWIIGTVAGILVIGLVGVFFYGSYSGLGSSL

>sp|Q332W5|CYF_LACSA Cytochrome f OS=Lactuca sativa OX=4236 GN=petA PE=3 SV=1

MQTRNNFSWIKEQITRSISVSLMIYIITRASISNAYPIFAQKGYENPREATGRIVCANCH

LANKPVDIEVPQTVLPDTVFEAVVRIPYDMQLKQVLANGKKGALNVGAVLILPEGFELAP

PDRISPEIKEKMGNLSFQSYRPNQKNILVIGPVPGQKYSEITFPILSPDPATKKDIHFLK

YPIYVGGNRGRGQIYPDGSKSNNTVYNATASGIVSKILRKEKGGYEITIADASDGRQVVD

IIPPGPELLVSEGESIKFEQPLTSNPNVGGFGQGDAEIVLQDPLRVQGLLFFLASVILAQ

IFLVLKKKQFEKVQLSEMNF

>sp|Q332W6|CEMA_LACSA Chloroplast envelope membrane protein OS=Lactuca sativa OX=4236 GN=cemA PE=3 SV=1

MEKKKAFTPLLYLASIIFLPWWISLSFQKSMESWVTNWWNTRQSEPFLNDIEEKSILEKF

IELEELLFLEEMIKEYSETHLQNLRIGIHKETIQLIKIHNEGRIHTILHFSTNIICFIIL

SGYSLLGNKELVILNSWVQEFLYNLSDTIKAFSLLLLTDLCIGFHSPHGWELMIGFVYKD

FGFVHNEQIISGLVSTFPVILDTIFKYWIFRYLNRVSPSLVVIYHSMND

>sp|Q332W7|YCF4_LACSA Photosystem I assembly protein Ycf4 OS=Lactuca sativa OX=4236 GN=ycf4 PE=3 SV=1

MSCRSEHIWIEPITGARKTSNFCWAVILFLGSLGFLLVGTSSYLGRNLISLFPSQEIVFF

PQGIVMSFYGIAGLFISSYLWCTISWNVGSGYDRFDRKDGIVCIFRWGFPGKNRRVFLQF

LIKDIQSVRIEVKEGIYARRVLYMDIRGQGAIPLTRTDENFTPREMEQKAAELAYFLRVP

IEVF

>sp|Q332W8|PSAI_LACSA Photosystem I reaction center subunit VIII OS=Lactuca sativa OX=4236 GN=psaI PE=3 SV=1

MTTLNFPSVLVPLVGLVFPAIAMASLFLHVQKNKIV

>sp|Q332X2|ATPE_LACSA ATP synthase epsilon chain, chloroplastic OS=Lactuca sativa OX=4236 GN=atpE PE=3 SV=1

MTLNLCVLTPNRIVWDSEVKEIILSTNSGQIGVLPNHAPIATSVDIGILRIRLNDQWLTM

ALMGGFARIGNNEITVLVNDAEKSGDIDPQEAQQTLEIAEAALRKAEGKRQTIEANLALR

RARTRVEAINAIS

>sp|Q332Y0|RR14_LACSA 30S ribosomal protein S14, chloroplastic OS=Lactuca sativa OX=4236 GN=rps14 PE=3 SV=1

MAKKSLIQREKKRQKLEQKYHLIRRSSKKEISKVRSLSDKWEIYGKLQSPPRNSAPTRLH

RRCFSTGRPRANYRDFGLSGHILREMVHACLLPGATRSSW

>sp|Q332Y1|PSBZ_LACSA Photosystem II reaction center protein Z OS=Lactuca sativa OX=4236 GN=psbZ PE=3 SV=1

MTLAFQLAVFALIATSSILLISVPVVFASPDGWSSNKNVVFSGTSLWIGLVFLVGILNSL

IS

>sp|Q332Z6|RR16_LACSA 30S ribosomal protein S16, chloroplastic OS=Lactuca sativa OX=4236 GN=rps16 PE=3 SV=1

MVKLRLKRCGRKQRAVYRIVAIDVRSRREGRDLRKVGFYDPIKNQTYLNVPAILYFLEKG

AQPTGTVQDILKKAEVFKELCPNQTKFN

>sp|Q332Z7|MATK_LACSA Maturase K OS=Lactuca sativa OX=4236 GN=matK PE=3 SV=1

MEKFQSYLGLDRSQQHHFLYPLIFQEYIYVLAHDHGLTRSILLENAGYDNKSSLLIVKRL

INRMYQQNHLILSVNNSKQTPFLGHNKNFYSQVMSEVSSIIMEIPLSLRLISYLERKGVV

KSDNLRSIHSIFSFLEDNFSHLNYVLDILIPYPAHLEILVQALRYWIKDASSLHLLRFFL

HECHNWDSLITSNSKKASSSFSKRNHRLFFFLYTSHLCEYESGFLFLRNQSSHLRSTSSG

ALIERIYFYGKIDHLAEVFARAFQANLWLFKDPFMHYVRYQGKSILGSKGTFLLMNKWKY

YFVNFWKSYFYLWSQPGRIYINQLSNHSLDFLGYRSSVRLKPSMVRSQMLENAFIIENAI

KKFETIVPIMPLIGSLAKSKFCNALGHPIGKAIWADFSDSDIIDRFGRIYRNLSHYHSGS

SKKKSLYRVKYILRLSCARTLARKHKSTVRAFLKRFGSELLEEFFTEEEQVFSLTFPRVS

SISRRLSRRRIWYLDIICINDLANHE

>sp|Q40250|RBS_LACSA Ribulose bisphosphate carboxylase small subunit, chloroplastic OS=Lactuca sativa OX=4236 GN=RBCS PE=2 SV=1

MASISSSAIATVNRTTSTQASLAAPFTGLKSNVAFPVTKKANNDFSSLPSNGGRVQCMKV

WPPIGLKKYETLSYLPPLSDEALSKEIDYLIRNKWIPCLEFELEHGFVYREHHHSPGYYD

GRYWTMWKLPMFGCTDSAQVMKEVGECKKEYPNAFIRVIGFDNIRQVQCISFIVAKPPGV

L

>sp|Q56P07|ATPF_LACSA ATP synthase subunit b, chloroplastic OS=Lactuca sativa OX=4236 GN=atpF PE=3 SV=1

MKNVTDSFVSLGHWPSAGSFGFNTDILATNLINLSVVLGVLIFFGKGVLSDLLDNRKQRI

LNTIRNSEELREGAIEQLEKARARLRKVEIEADQFRVNGYSEIEREKLNLIDSTYKTLEQ

LENYKNETINFEQQKASNQVRQRVFQQALQGALGTLNSCLNSELHLRTISANIGILGAMK

EITD

>sp|Q56P09|ATPI_LACSA ATP synthase subunit a, chloroplastic OS=Lactuca sativa OX=4236 GN=atpI PE=3 SV=1

MNVLSCSINTLNGLYDISGVEVGQHFYWKIGGFQVHGQVLITSWVVIAILLASATLAVRN

PQTIPTSGQNFFEYVLEFIRDVSKTQIGEEYGPWVPFIGTMFLFIFVSNWSGALLPWKII

QLPHGELAAPTNDINTTVALALLTSVAYFYAGLSKKGLGYFGKYIQPTPILLPINILEDF

TKPLSLSFRLFGNILADELVVVVLVSLVPSVVPIPVMFLGLFTSGIQALIFATLAAAYIG

ESMEGHH

>sp|Q56P10|RR2_LACSA 30S ribosomal protein S2, chloroplastic OS=Lactuca sativa OX=4236 GN=rps2 PE=3 SV=1

MTRRYWNINLEEMMEAGVHFGHGTRKWNPKMAPYISAKRKGIHITNLTRTARFLSEACDL

VFDAASRGKQFLIVGTKNKEADSVAWAAIRARCHYVNKKWLGGMLTNWSTTETRLHKFRD

LRTEQKTGGLDRLPKRDAAMLKRQLSHLQTYLGGIKYMTGLPDIVIIVDQHEEYTALQEC

ITLGIPTICLIDTNCDPDLADISIPANDDAISSIRLILNKLVFAICEGRSGYIRNP

>sp|Q56P14|PSBM_LACSA Photosystem II reaction center protein M OS=Lactuca sativa OX=4236 GN=psbM PE=3 SV=1

MEVNILAFIATALFILVPTAFLLIIYVKTVSQNN

>sp|Q56P15|PETN_LACSA Cytochrome b6-f complex subunit 8 OS=Lactuca sativa OX=4236 GN=petN PE=3 SV=1

MDIVSLAWAALMVVFTFSLSLVVWGRSGL

>sp|Q56P16|PSBI_LACSA Photosystem II reaction center protein I OS=Lactuca sativa OX=4236 GN=psbI PE=3 SV=1

MLTLKLFVYTVVIFFVSLFIFGFLSNDPGRNPGREE

>sp|Q56P17|PSBK_LACSA Photosystem II reaction center protein K OS=Lactuca sativa OX=4236 GN=psbK PE=3 SV=1

MLNIFSLICLNSALYPSSLFFAKLPEAYAFLNPIVDVMPVIPLFFFLLAFVWQAAVSFR
