## Supplementary material for "Sustained non-photochemical quenching and regulation of PSII repair cycle during combined low temperature and high light stress in lettuce": Supplemental file 4_Sequences removed from combined database.docx

>tr|A0A2J6LSJ6|A0A2J6LSJ6_LACSA ATP synthase epsilon chain, chloroplastic OS=Lactuca sativa OX=4236 GN=LSAT_7X68801 PE=3 SV=1

MTLNLCVLTPNRIVWDSEVKEIILSTNSGQIGVLPNHAPIATSVDIGILRIRLNDQWLTM

ALMGGFARIGNNEITVLVNDAEKSGDIDPQEAQQTLEIAEAALRKAEGKRQTIEANLALR

RARTRVEAINAIS

>tr|A0A2J6MCD9|A0A2J6MCD9_LACSA ATP synthase subunit alpha OS=Lactuca sativa OX=4236 GN=LSAT_0X28780 PE=3 SV=1

MVTIQADEISNIIRERIEQYNREVKIVNTGTVLQVGDGIARIHGLDEVMAGELVEFEEGT

IGIALNLESTNVGVVLMGDGLLIQEGSSVKATGRIAQIPVSEAYLGRVINALAKPIDGRG

EISSSEYRLIESPAPGIISRRSVYEPLQTGLIAIDSMIPIGRGQRELIIGDRQTGKTAVA

TDTILNQQGKNVICVYVAIGQKASSVAQVVTNFQERGAMEYTIVVAETADSPATLQYLAP

YTGAALAEYFMYRERHTSIIYDDPSKQAQAYRQMSLLLRRPPGREAYPGDVFYLHSRLLE

RAAKLSSLLGEGSMTALPIVETQSGDVSAYIPTNVISITDGQIFLSADLFNAGIRPAINV

GISVSRVGSAAQIKAMKQVAGKLKLELAQFAELEAFAQFASDLDKATQNQLARGQRLREL

LKQSQSAPLGVEEQVLTIYTGTNGYLDSLEIGQVRKFLVELRTYLKTNKPQFQEIISSTK

TFTEEAEAILKEAIKEQRERFILQEQAA

>tr|A0A2J6MCG3|A0A2J6MCG3_LACSA Cytochrome f OS=Lactuca sativa OX=4236 GN=LSAT_1X38720 PE=3 SV=1

MIYIITRASISNAYPIFAQKGYENPREATGRIVCANCHLANKPVDIEVPQTVLPDTVFEA

VVRIPYDMQLKQVLANGKKGALNVGAVLILPEGFELAPPDRISPEIKEKMGNLSFQSYRP

NQKNILVIGPVPGQKYSEITFPILSPDPATKKDIHFLKYPIYVGGNRGRGQIYPDGSKSN

NTVYNATASGIVSKILRKEKGGYEITIADASDGRQVVDIIPPGPELLVSEGESIKFEQPL

TSNPNVGGFGQGDAEIVLQDPLRVQGLLFFLASVILAQIFLVLKKKQFEKVQLSEMNF

>tr|A0A2J6KXP3|A0A2J6KXP3_LACSA DNA-directed RNA polymerase subunit beta OS=Lactuca sativa OX=4236 GN=LSAT_1X118620 PE=3 SV=1

MSTIPGFNQIQFEGFCRFIDQGLTEELSKFPKIEDTNQEIDFELFLERYQLVEPSIKERD

AVYESLTYSSELYVSARLIWKNDRRRYIQEQTILIGKIPLMTSLGAFIVNGIYRIVINQI

LQSPGIYYQSELNDNGISVYTGTIISDWGGRLELEIDRKTRIWVRVSRQQKLSILVLLSA

MGLNIREILENVCYPELFLSFLNDKKQIGSKENAILEFYQQFACVEGDPVFSESLSKDLQ

KKFFQQRCELGGIGRRNMNRRLNLDIPQNNTFLLPRDILAAADRLIRIKFGMGTLDDMNH

LQNKRIRSVADLLQEQFGLALVRLENMARGNIYAALKHNWTPTPQNLVNSTPLTDTYKVF

FRLHPLSQVLDRTNPLTQIVHGRKLSYLGPGGLTARTATFPIRDIHPSHYGRICPIDTSE

GINVGLIGSLAIHARIGRWGSLESPFYKISERSKGARMLYLSPGRDEYYMVAAGNSLALN

QGIQEEQVVPARYRQEFLTIAWEQVHLRSIFSFQYFSIGASLIPFIEHNDANRALMSSNM

QRQAVPLSQSEKCIVGTGLEGQAALDSGALAIAEHEGEIIYTDTDKILLSGNGDTLRIPL

VMYQRSNKNTCMHQKPQVQRGKCIKKGQILAYGAATVGGELALGKNVLVAYMPWEGYNFE

DAVLISERLVYEDIYTSFHIRKYEIQINQGSERVTNEIPHLEVHLLRNLDKNGIVMLGSW

VETGDILVGKLTPQMVKESSYAPEDRLLRTILGMRVYTSKETCLKLPIGGRGRVIDVRWV

QSSKTDETEKTESIRVYILQKREIKVGDKVAGRHGNKGIISKILPRQDMPYLQDGRPVDM

VFNPLGVPSRMNVGQIFESSLGLAGGLLDRHYRIAPFDERYEQEASRKLVFSELYEASKQ

TVNPWIFEPESPGKSRIFDGRTGDPFEQPVIIGKPYILKLIHQVDDKIHGRSSGRYSRLT

QQPLKGRAKKGGQRVGEMEVWALEGFGVAYILQEMLTYKSDHIRARQEVLGTIIFGGRIP

TPEDAPESFRLFVRELRSLALELNHFLVSEKTFQLNRKEA

>tr|A0A2J6JMI9|A0A2J6JMI9_LACSA DNA-directed RNA polymerase subunit OS=Lactuca sativa OX=4236 GN=LSAT_1X96161 PE=3 SV=1

MIDRYTHQQLRIGLVSPQQISTWSKKILPNGEIVGEVTKPYTFHYKTNKPEKDGLFCERI

FGPIKSGICACGNYRVIGDEKEDPQFCEQCGVEFVDSRIRRYQMGYIKLAYPVMHVWYLK

RLPSYIVTLLDKPLNELEDLVYCGIRNEKLSSHLIQSGCPETFTSDFEGGRSYPNFYFAR

PIDKKPTFLRLRGLLEYEIQPWKYRIPIFFTTRSFDTFRNREMSTGGGSIRQQLANLDLR

IIIDYSLVEWKELEEEEPTGNEWEDRKVGRRKDFLLRRMELAKHFIRTNIEPKWMVLRLL

PVLPPELRPIYHIDEDKLVTSDINEIYRRIIYRNNTLTDLLTTSIATPEELIISQEKLLQ

EAVDALLDNGICGQPMRDDHNRVYKSLSDVIEGKEGRVRETLLGKRVDYSGRSVIVVGPS

LSLHRCGLPREIAIELFQAFVIRDLIRKHLASNIGVAKSQIRKKKPIVWEILQEILDDHP

VLLNRAPTLHRLGIQAFLPVLVEGRAICLHPLVCKGFNADFDGDQMAVHVPLSLEAQAEA

RLLMFSHMNLLSPTIGDPISAPTQDMLSGLYVLTSGNRRGICVNRYNPCNRRNYQNEDNN

YKYTKKKEPFFCNPYDAIGAYRQKRINLGSPLWLRWRLDQRVIAAREVPIEIHYESVGTY

YEIYGQS

>tr|A0A2J6KMV4|A0A2J6KMV4_LACSA Maturase K OS=Lactuca sativa OX=4236 GN=matK PE=3 SV=1

MYQQNHLILSVNNSKQTPFLGHNKNFYSQVMSEVSSIIMEIPLSLRLISYLERKGVVKSD

NLRSIHSIFSFLEDNFSHLNYVLDILIPYPAHLEILVQALRYWIKDASSLHLLRFFLHEC

HNWDSLITSNSKKASSSFSKRNHRLFFFLYTSHLCEYESGFLFLRNQSSHLRSTSSGALI

ERIYFYGKIDHLAEVFARAFQANLWLFKDPFMHYVRYQGKSILGSKGTFLLMNKWKYYFV

NFWKSYFYLWSQPGRIYINQLSNHSLDFLGYRSSVRLKPSMVRSQMLENAFIIENAIKKF

ETIVPIMPLIGSLAKSKFCNALGHPIGKAIWADFSDSDIIDRFGRIYRNLSHYHSGSSKK

KSLYRVKYILRLSCARTLARKHKSTVRAFLKRFGSELLEEFFTEEEQVFSLTFPRVSSIS

RRLSRRRIWYLDIICINDLANHE

>tr|A0A2J6KR40|A0A2J6KR40_LACSA Maturase K OS=Lactuca sativa OX=4236 GN=LSAT_2X89740 PE=3 SV=1

MEIPLSLRLISYLERKGVVKSDNLRSIHSIFSFLEDNFSHLNYVLDILIPYPAHLEILVQ

ALRYWIKDASSLHLLRFFLHECHNWDSLITSNSKKASSSFSKRNHRLFFFLYTSHLCEYE

SGFLFLHHLAEVFARAFQANLWLFKDPFIYVRYQGKSIIGSKGTSLLMNKWKYYFKSYFY

LWSQPRRIYINQLSNHSLDFLGYRSSVRLKPSMVRSQMLENAFIIENAIKKFETIVPIMP

LIGSLANSKFCNALGHPIGETIWANLLDPDIIDRFGRI

>tr|A0A2J6MGL0|A0A2J6MGL0_LACSA mRNA cap-binding protein OS=Lactuca sativa OX=4236 GN=LSAT_6X68620 PE=3 SV=1

MEVFKPSKLPANADFHLFKAGVQPKWEDPECASGGKWTVTSSRKPTLETMWLETLMALIG

EQFDEADEICGVVASVRQRQDKLSLWTKNAANEAAQMSIGRKWKEIIDVTDKIIYNFHDD

SKTRTSKGRYNV

>tr|A0A2J6LHG4|A0A2J6LHG4_LACSA Photosystem I P700 chlorophyll a apoprotein A1 OS=Lactuca sativa OX=4236 GN=LSAT_1X88121 PE=3 SV=1

MIIRSPEPEVKILVDRDHIKTSFEEWARPGHFSRTIAKGPETTTWIWNLHADAHDFDSHT

SDLEEISRKVFSAHFGQLSIIFLWLSGMYFHGARFSNYEAWLSDPTHIRPSAQVVWPIVG

QEILNGDVGGGFRGIQITSGFFQIWRASGITSELQLYCTAIGGLVFAALMLFAGWFHYHK

AAPKLAWFQDVESMLNHHLAGLLGLGSLSWAGHQVHVSLPINQFLNAGVDPKEIPLPHEF

ILNRDLLAQLYPSFAEGATPFFTLNWSKYADFLTFRGGLDPVTGGLWLTDTAHHHLAIAI

LFLIAGHMYRTNWGIGHGLKDILEAHKGPFTGQGHKGLYEILTTSWHAQLSLNLAMLGSL

TIVVAHHMYAMPPYPYLATDYGTQLSLFTHHMWIGGFLIVGAAAHAAIFMVRDYDPTTRY

NDLLDRVLRHRDAIISHLNWACIFLGFHSFGLYIHNDTMSALGRPQDMFSDTAIQLQPVF

AQWIQNTHALAPGATAPGATASTSLTWGGGDLVAVGGKVALLPIPLGTADFLVHHIHAFT

IHVTVLILLKGVLFARSSRLIPDKANLGFRFPCDGPGRGGTCQVSAWDHVFLGLFWMYNS

ISVVIFHFSWKMQSDVWGSISDQGVVTHITGGNFAQSSITINGWLRDFLWAQASQVIQSY

GSSLSAYGLFFLGAHFVWAFSLMFLFSGRGYWQELIESIVWAHNKLKVAPATQPRALSIV

QGRAVGVTHYLLGGIATTWAFFLARIIAVG

>tr|A0A2J6LR30|A0A2J6LR30_LACSA photosystem I OS=Lactuca sativa OX=4236 GN=LSAT_1X5820 PE=3 SV=1

MALRFPRFSQGLAQDPTTRRIWFGIATAHDFESHDDITEERLYQNIFASHFGQLAIIFLW

TSGNLFHVAWQGNFESWVQDPLHVRPIAHAIWDPHFGQPAVEAFTRGGALGPVNIAYSGV

YQWWYTIGLRTNEDLYTGALFLLLISAISLIAGWLHLQPKWKPSVSWFKNAESRLNHHLS

GLFGVSSLAWTGHLVHVAIPASRGEYVRWNNFLDVLPHPQGLGPLFTGQWNLYAQNPDSG

SHLFGTSQGAGTAILTLLGGFHPQTQSLWLTDMAHHHLAIAFLFLIAGHMYRTNFGIGHS

MKDLLDAHIPPGGRLGRGHKGLYDTINNSLHFQLGLALASLGVITSLVAQHMYSLPAYAF

IAQDFTTQAALYTHHQYIAGFIMTGAFAHGAIFFIRDYNPEQNEDNVLARMLEHKEAIIS

HLSWASLFLGFHTLGLYVHNDVMLAFGTPEKQILIEPIFAQWIQSAHGKTSYGFDILLSS

TNGPAFNAGRSIWLPGWLNAINENSNSLFLTIGPGDFLVHHAIALGLHTTTLILVKGALD

ARGSKLMPDKKDFGYSFPCDGPGRGGTCDISAWDAFYLAVFWMLNTIGWVTFYWHWKHIT

LWQGNVSQFNESSTYLMGWLRDYLWLNSSQLINGYNPFGMNSLSVWAWMFLFGHLVWATG

FMFLISWRGYWQELIETLAWAHERTPLANLIRWRDKPVALSIVQARLVGLAHFSVGYIFT

YAAFLIASTSGKFG

>tr|A0A2J6M888|A0A2J6M888_LACSA Photosystem II D2 protein OS=Lactuca sativa OX=4236 GN=LSAT_1X88080 PE=3 SV=1

MTIALGKVTKDENDLFDIMDDWLRRDRFVFVGWSGLLLFPCAYFAVGGWFTGTTFVTSWY

THGLASSYLEGCNFLTAAVSTPANSLAHSLLLLWGPEAQGDFTRWCQLGGLWTFVALHGA

FGLIGFMLRQFELARSVQLRPYNAIAFSGPIAVFVSVFLIYPLGQSGWFFAPSFGVAAIF

RFILFFQGFHNWTLNPFHMMGVAGVLGAALLCAIHGATVENTLFEDGDGANTFRAFNPTQ

AEETYSMVTANRFWSQIFGVAFSNKRWLHFFMLFVPVTGLWMSALGVVGLALNLRAYDFV

SQEIRAAEDPEFETFYTKNILLNEGIRAWMAAQDQPHENLIFPEEVLPRGNAL

>tr|A0A2J6MEE0|A0A2J6MEE0_LACSA Ribosomal protein S7 OS=Lactuca sativa OX=4236 GN=LSAT_3X14241 PE=3 SV=1

MSRRGTAEEKTAKSDPIYRNRLVNMLVNRILKHGKKSLAYQIIYRAVKKIQQKTETNPLS

VLRQAIHGVTPGIAVKARRVGGSTQQVPIEIGSTQGKALAIRWLLAASRKRPGRNMAFKL

SSELVDAAKGSGDAIRKREETHKMAESNRAFAHFR

>tr|A0A2J6LMH1|A0A2J6LMH1_LACSA Ribulose bisphosphate carboxylase small subunit, chloroplastic OS=Lactuca sativa OX=4236 GN=RBCS PE=3 SV=1

MASISSSAIATVNRTTSTQASLAAPFTGLKSNVAFPVTKKANNDFSSLPSNGGRVQCMKV

WPPIGLKKYETLSYLPPLSDEALSKEIDYLIRNKWIPCLEFELEHGFVYREHHHSPGYYD

GRYWTMWKLPMFGCTDSAQVMKEVGECKKEYPNAFIRVIGFDNIRQVQCISFIVAKPPGV

L

>tr|A0A2J6JZ26|A0A2J6JZ26_LACSA VDE domain-containing protein OS=Lactuca sativa OX=4236 GN=LSAT_7X62281 PE=4 SV=1

MALSLHTVFLCKEEALNLYARSPCNERFHRSGQPPTNIIMMKIRSNNGYFNSFRLFTSYK

TSSFSDSSHCKDKSQICSIDTSFEEIQRFDLKRGMTLILEKQWRQFIQLAIVLVCTFVIV

PRVDAVDALKTCACLLKECRIELAKCIANPSCAANVACLQTCNNRPDETECQIKCGDLFE

NSVVDQFNECAVSRKKCVPRKSDVGEFPVPDRNAVVQNFNMKDFSGKWYITSGLNPTFDA

FDCQLHEFHMENDKLVGNLTWRIKTLDGGFFTRSAVQTFVQDPDLPGALYNHDNEFLHYQ

DDWYILSSQIENKPDDYIFVYYRGRNDAWDGYGGSVIYTRSPTLPESIIPNLQKAAKSVG

RDFNNFITTDNSCGPEPPLVERLEKTAEEGEKLLIKEAVEIEEEVEKEVEKVRDTEMTLF

QRLLEGFKELQQDEENFVRELSKEEKEILNELQMEATEVEKLFGRALPIRKLR

>sp|A0A2J6L8Y7|IF4E1_LACSA Eukaryotic translation initiation factor 4E-1 OS=Lactuca sativa OX=4236 GN=eIF4E PE=1 SV=1

MVEEIMKSEEQKLIDVNKHRGVRSDGEEDEQLEEGEIVGGDADTLSSSSSSRPGTAIAQH

PLEHSWTFWFDTPSAKSKQVAWGSSMRPIYTFSSVEEFWSLYNNIHRPSKLAQGADFYCF

KNKIEPKWEDPVCANGGKWTMTFTKAKSDTCWLYTLLAMIGEQFDHGDDICGAVVNVRAR

QEKIALWTKNAANESAQLSIGKQWKEFIDYNDTIGFIFHEDAKTLDRSAKNKYTV

>tr|A0A2J6KE35|A0A2J6KE35_LACSA Germacrene A oxidase OS=Lactuca sativa OX=4236 GN=LSAT_8X115261 PE=3 SV=1

MELSITTSIALATIVFFLYKLATRPKSTKKQLPEASRLPIIGHMHHLIGTMPHRGVMDLA

RKHGSLMHLQLGEVSTIVVSSPKWAKEILTTYDITFANRPETLTGEIIAYHNTDIVLAPY

GEYWRQLRKLCTLELLSVKKVKSFQSIREEECWNLVKEVKESGSGKPINLSESIFKMIAT

ILSRAAFGKGIKDQREFTEIVKEILRQTGGFDVADIFPSKKFLHHLSGKRARLTSIHKKL

DNLINNIVAEHHVSTSSKANETLLDVLLRLKDSAEFPLTADNVKAIILDMFGAGTDTSSA

TVEWAISELIRCPRAMEKVQAELRQALNGKEKIQEEDIQDLAYLNLVIRETLRLHPPLPL

VMPRECREPVNLAGYEIANKTKLIVNVFAINRDPEYWKDAEAFIPERFENNPNNIMGADY

EYLPFGAGRRMCPGAALGLANVQLPLANILYHFNWKLPNGASHDQLDMTESFGATVQRKT

ELLLVPSF

>tr|A0A2J6KR38|A0A2J6KR38_LACSA Ribulose bisphosphate carboxylase large chain OS=Lactuca sativa OX=4236 GN=LSAT_2X89801 PE=3 SV=1

MSCREGFMSPQTETKASVGFKAGVKDYKLTYYTPEYETKDTDILAAFRVTPQPGVPPEEA

GAAVAAESSTGTWTTVWTDGLTSLDRYKGRCYGIEPVPGEENQYIAYVAYPLDLFEEGSV

TNMFTSIVGNVFGFKALRALRLEDLRIPTAYVKTFQGPPHGIQVERDKLNKYGRPLLGCT

IKPKLGLSAKNYGRAVYECLRGGLDFTKDDENVNSQPFMRWRDRFLFCAEAIFKSQAETG

EIKGHYLNATAGTCEEMMKRAIFARELGVPIVMHDYLTGGFTANTSLAHYCRDNGLLLHI

HRAMHAVIDRQKNHGIHFRVLAKALRMSGGDHIHSGTVVGKLEGEREITLGFVDLLRDDF

IEKDRSRGIYFTQDWVSLPGVLPVASGGIHVWHMPALTEIFGDDSVLQFGGGTLGHPWGN

APGAVANRVALEACVQARNEGRDLATEGNEIIREATKWSPELAAACEVWKEIKFEFQAMD

TLDQ

>tr|A0A2J6LH98|A0A2J6LH98_LACSA Photosystem II protein D1 OS=Lactuca sativa OX=4236 GN=LSAT_4X5440 PE=3 SV=1

MIAILERCENKILWGRFCNSITNTENRLYLGWFGVLMIPTLLTTNYVFIIPFIAAPLVDI

DGIRELVSGSLLYRNNIISCAIIPTSAASLHFYLIWEATYVDEWLYNGGPYELVVLSFLL

GVACYMGREWELSFCLGMQPWIVVAYSTPVAAKHNILMHPFHMLGVAGVFDGSLFSVMHG

SLVTSSLIRESTKNESANEGYKFSQEEETYNIVAAHGYFGQLIFQYASFNSSHSLHLFLA

AWPVVGIWVTALVINTMAFNLNGFDFNQSVVDSQGRVIHTWADIIKRANLGMKFMHERNA

HNFPLDLAAIEALSTNG

>tr|A0A2J6LQX2|A0A2J6LQX2_LACSA Photosystem II protein D1 OS=Lactuca sativa OX=4236 GN=LSAT_8X133980 PE=3 SV=1

MGREWELSFRLGMRPWIAVAYSAPVAAATAVFLIYPIGQGSFSDGMPLGISGTFNFMIVF

QAEHNILMHPFHMLGVAGVFGGSLFSAMHGSLVTSSLIRETTENESANEGYRFGQEEETY

NIVAAHGYFGRLIFQYASFNNSRSLHFFLAAWPVVGIWFTALGISTMAFNLNGFNFNQSV

VDSQGRVINTWADIINRANLGMEVMHERNAHNFPLDLAAIEAPSTNG

>tr|A0A2J6JPG4|A0A2J6JPG4_LACSA Photosystem II protein D1 OS=Lactuca sativa OX=4236 GN=LSAT_7X82620 PE=3 SV=1

MIVFQDEHNILMHPFHMLCVTIVFDGSLFSVMHGSLVTSSLIRETTENKSANEGYIFSQE

EETYNIVAAHGYFGQLIFQFASFNNSRSLHFFLTDCPVVGIWFTALGISTMAFNLNGFNF

KQSVVDCQGRVIKTWVDIINRANLSMEVMHERNAYNFPLDLAAIEAP

>tr|A0A2J6LMD3|A0A2J6LMD3_LACSA Photosystem II protein D1 OS=Lactuca sativa OX=4236 GN=LSAT_3X104200 PE=3 SV=1

MIPTLLTATSVFIIAFIAAPPVDIDGIREPVSGSLLYGNNIISGAIIPTSAAIGLHFYPI

WEAASVDEWLYNGGPYELIVLHFLLGVACYMGREWDF

>tr|A0A2J6JM92|A0A2J6JM92_LACSA Photosystem II protein D1 OS=Lactuca sativa OX=4236 GN=LSAT_4X51841 PE=3 SV=1

MNSRNRKECSEIQLKFKFVIRVFDYHKSLSYPQLDILRDPNLIRSFAGKRIRIGKWGEPN

LLRLCNSLEAPKTKRRHVQARYAATSVFIITFIVAPPLDIVGIREPVFGSLLYGNNIISD

AIIPTSAAIGLHFYPIWEAASVDEWLYNGGPYELIVLHFLLGVACYMGREWELSFRLGM

>tr|A0A2J6KRM9|A0A2J6KRM9_LACSA NAD(P)H-quinone oxidoreductase subunit 2, chloroplastic OS=Lactuca sativa OX=4236 GN=LSAT_9X18660 PE=4 SV=1

MAIIEFLLFVLTATIGGILCSYLLSGYTKKDVRSNEATMKYLLMGGASSFILVHGFSWPY

GSSRGEIELQEIVNGLINTQMYNSPGISIVLIFIIVGIRFKLSPAPFHQWTPNVYEGVRF

VKQVRNDESLYDKQIRFASSLLRVVPTKYQTNDMLHSWFSSFRDYECNRSIRRQKDHPKM

IISWLLRTNQIRWFYFLTCSYGTKIEKIEKISHSQPLMKDSSKKVRNPLFDSNSPTPVVA

FLSVTSKVAASASATRIFDIPFYFSSNEWHLLLEILAILSMILGNLIAITQTSMKRMLAY

SSIGQIGYVIIGIIVGDSNDGYASMITYMLFYISMNLGTFACIVLFGLRTGTENIRDYAG

LYTKDPFLALSLALCLLSLGGLPPLAGFFGKLYLFWCGWQAGLYFLVLIGLLTSVVSIYY

YLKIIKLLMTGRNQEITPHVRNYRRSPLRSNNSIELSMIVCVIASTIPGISMNPIIAIAQ

DTLF

>tr|A0A2J6KNT3|A0A2J6KNT3_LACSA NAD(P)H-quinone oxidoreductase subunit 2, chloroplastic OS=Lactuca sativa OX=4236 GN=LSAT_3X92301 PE=4 SV=1

MAQCPICSRKAAQLNVSEDFLTEFLLFVLTATIGGMFLCGANGLITIFVAPECFSLCSYL

LSGYTKKDVRSNEATIKYLLMGGASSSILVHGFSWLYGSSGGEIELQEIVNGLINTQMYN

SPGISIALIFITVGIGFKLSPAPSHQWTPDVYEGVRFVKQVRNDESLYDKQIRFASSLLR

VVPTKYQTNDMLHSWFSSFRDYECNRSIRRQKDHPKMIISWLLRTNQIRWFYFLTCSYGT

KIEKIEKISHSQPLMKDSSKKVRNPLFDSNSPTPVVAFLSVTSKVAASASATRIFDIPFY

FSSNEWHLLLEILAILSMILGNLIAITQTSMKRMLAYSSIGQIGYVIIGIIVGDSNDGYA

SMITYMLFYISMNLGTFACIVLFGLRTGTENIRDYAGLYTKDPFLALSLALCLLSLGGLP

PLAGFFGKLYLFWCGWQAGLYFLVLIGLLTSVVSIYYYLKIIKLLMTGRNQEITPHVRNY

RRSPLRSNNSIELSMIVCVIASTIPGISMNPIIAIAQDTLF

>tr|A0A2J6L0I4|A0A2J6L0I4_LACSA Proton_antipo_M domain-containing protein OS=Lactuca sativa OX=4236 GN=LSAT_3X112740 PE=4 SV=1

MYNSPGISIALIFITVGIGFKLSPAPSHQWTPDVYEGVRFVKQVRNDESLYDKQIRFASS

LLRVVPTKYQTNDMLHSWFSSFRDYECNRSIRRQKDHPKMIISWLLRTNQIRWFYFLTCS

YGTKIEKIEKISHSQPLMKDSSKKVRNPLFDSNSPTPVVAFLSVTSKVAASASATRIFDI

PFYFSSNEWHLLLEILAILSMILGNLIAITQTSMKRMLAYSSIGQIGYVIIGIIVGDSND

GYASMITYMLFYISMNLGTFACIVLFGLRTGTENIRDYAGLYTKDPFLALSLALCLLSLG

GLPPLAGRPIFLGFNRTPYKRCFYLLLSKNNQI

>tr|A0A2J6LTM8|A0A2J6LTM8_LACSA Proton_antipo_M domain-containing protein OS=Lactuca sativa OX=4236 GN=LSAT_6X37880 PE=4 SV=1

MLHSWFSSFREYECNRSIRRQKDHPKMIISWLLRTNQIRWFYFLTCSYGTKIEKIEKISH

SQPLMKDSLKSYGLKEVRNLLFDSNSPTPVVAFLSVTSKVAASASATRIFDIPFYFSSNE

WHLLLEILAILSMILGNLIAITQTSMKRMLAYSSIGQIGYVIIGIIVGDSNDGYASMITY

TLFYISTNLGTFACIVLFGLHTETQNIRDYAGLYTKDPSQSFVIVSLLSRA

>tr|A0A2J6KBS0|A0A2J6KBS0_LACSA Glutamine synthetase OS=Lactuca sativa OX=4236 GN=LSAT_2X68621 PE=3 SV=1

MALLSDLVNLDLSSITDKIIAEYIWIGGSGMDLRSKARTLSGPVSDPSELPKWNYDGSST

GQAPGEDSEVIIYPQAIFKDPFRRGNHILVMCDAYTPAGEPIPTNKRAAAAKIFSNPEVE

KEVTWYGIEQEYTLLQKDTNWPLGWPLGGFPGPQGPYYCGIGADKAFGRDIVDAHYKACL

YAGVNISGINGEVMPGQWEFQVGPSVGIAAADQIWVARYILERITEIYGVVVSFDPKPIP

GDWNGAGAHTNYSTKTMREEGGYEVIKKAIEKLGLRHKEHIAAYGEGNERRLTGRHETAD

INTFLWGVANRGASIRVGRDTEKEGKGYFEDRRPASNMDPYVVTSMIAETTILWDNKS

>tr|A0A2J6L084|A0A2J6L084_LACSA Glutamine synthetase OS=Lactuca sativa OX=4236 GN=LSAT_3X63880 PE=3 SV=1

MALLNDLINLNLTESTTKIIAEYIWIGGSGMDLRSKARTLPEPVTDPKKLPKWNYDGSST

GQAPGEDSEVIVWPQAIFKDPFRGGNNILVICDAYTPAGEPIPTNKRHAAAKIFSDPKVE

KEIPWYGIEQEYTLLQKDINWPLGWPQGGFPGPQGPYYCGIGADKAFGRDIVDAHYKACL

YAGINISGINGEVMPGQWEFQVGPSVGISAGDEIWAARYILERITEIAGVVVSFDPKPIK

GDWNGAGAHTNYSTKSMREEGGYEVIKKAIEKLGLRHKEHIAAYGEGNERRLTGRHETAD

INTFKWGVANRGASIRVGRDTEKEGKGYFEDRRPASNMDPYVVTAMIAETTLLL

>tr|A0A2J6KBK9|A0A2J6KBK9_LACSA glutamine synthetase OS=Lactuca sativa OX=4236 GN=LSAT_2X68520 PE=3 SV=1

MSRVTDFINLNLNPSTIIAEYIWIGGSGLDLRSKARTLSTPIDDPQKLPKWNYDGSSTGQ

AHGNDTEVILHPQAVFKDPFRRGNNILVLCDAYNPLGDPIYTNKRFDAAKIFGHPDVVAE

APWFGLEQEYTLLQKDTKWPLGWPIGGFPRPQGPYYCGVGAEKAFGRDIVDAHYKACLYA

GITIGGVNAEVMPGQWEFQVGPSAGITAADELWVARYILERVAEIAGVIVSFNPKPVPGD

WNGAGAHTNYSTKSMRKEGGYEAIQKAIEKLGLRHEEHIASYGEGNEHRLTGLHETADIN

TFSWGVAKRGVSIRVGRETEKEGKGYFEDRRPGSNMDPYVVTSMIAETTILWKP

>tr|A0A2J6MJ80|A0A2J6MJ80_LACSA Glutamine synthetase OS=Lactuca sativa OX=4236 GN=LSAT_5X77601 PE=3 SV=1

MAQCLAPSVQWQTRLTKNAMETSSMTSKMWNSVSFKQSKKGAFKSSTKFRICASSSGTIN

RVEDLLNLDVTPYTDKIIAEYIWIGGSGTDVRSKSRTLSKPVEHPSELPKWNYDGSSTGQ

APGEDSEVILYPQAIFKDPFRGGNNILVICDTYTPQGVPIPTNKRAKAAEIFSDPKVVAQ

VPWFGIEQEYTLLQQDVKWPLGWPVGGYPGPQGPYYCGAGADKSFGRDISDAHYKACLYA

GINISGTNGEVMPGQWEFQVGPSVGIEAGDHIWCARYLLERITEQAGVVLTLDPKPIEGD

WNGAGCHTNYSTLSMREEGGFEVIKKAILNLSLRHADHISAYGEGNERRLTGKHETASIN

TFSWGVANRGCSIRVGRDTEKAGKGYLEDRRPASNMDPYTVTGLLAETTLLWEPTLEAEA

LAAQKLALNV

>tr|A0A2J6LYN5|A0A2J6LYN5_LACSA Glutamine synthetase OS=Lactuca sativa OX=4236 GN=LSAT_0X10461 PE=3 SV=1

MASSLGLTHLNGLTSSRVALPSSRRRVSGRARVGGLKIVASEMKQQEPDLSVNVNGLHMP

NPFVIGSGPPGTNYKVMKKAFDEGWGAVIAKTVSLDAAKVINVTPRYARLRAGVNGSSKG

QVIGWQNIELISDRPLETMLKEFKQLKEEYPDRILIASIMEEYNKAAWEELVDRVEQTGI

DAIEVNFSCPHGMPERKMGAAVGQDCDLLEEVCGWINAKATVPVWAKMTPNITDITQPAR

VALKSGCEGIAAINTIMSVMGINLKTLHPEPCVEGYSTPGGYSSKAVHPIALAKVMSIAQ

MMKSEFQDGDYSISGIGGVESGGDAAEFILLGANTVQVCTGVMMYGYDMVSKLCEELKDF

MRAHNFATIEEFRGASLQYFTTHTELVRIQQEAIKERRAIKKGLASDKDWTGDGFVKESE

SMTLPGPVSDPSELPKWNYDGSSTKQARGEDSEIIIHPQAIYKDPFRRGNNILVMCDAYN

PQGDPILTNKRHNAAKIFSHSDVIAEEPWFGIEQEYTLLQKDVKWPIGWPIGGYPGPQGP

YYCGTGADKAFGRDIVDAHYKACLYAGINISGTNGEVMPGQWEFQVGPSVGISAGDQVWV

ARYILERITEIAGVVLSFDPKPVHGDWNGAGAHTNYSTKSMRSDGGFDVIRKAIGKLEKR

HKEHMAAYGEGNERRLTGKHETADINNFNWGVADRGASVRVGRETEKANKGYFEDRRPSS

NMDPYVVTSMIAETTIIWQP

>tr|A0A2J6JYU9|A0A2J6JYU9_LACSA Triosephosphate isomerase, cytosolic OS=Lactuca sativa OX=4236 GN=LSAT_2X5900 PE=3 SV=1

MGRKFFVGGNWKCNGTAEDVKKIVATLNAGQIPSTDVVEVVVSPPFVFLTSVKSELRPEI

QVAAQNCWVKKGGAFTGEVSAEMLANLGVPWVILGHSERRALLNETNEFGGDKVAYALSQ

GLKVIACVGETLEQREAGTTMEVVAAQTKAIADKISSWDNVVLAYEPVWAIGTGKVASPA

QAQEVHAGLRKWFCDNVSAEVSASTRIIYGGSVSGSNCKELGGQTDVDGFLVGGASLKPE

FIDIIKAAEVKKSA

>tr|A0A2J6L4E3|A0A2J6L4E3_LACSA Proton_antipo_M domain-containing protein OS=Lactuca sativa OX=4236 GN=LSAT_1X14200 PE=4 SV=1

MGFILIGIASITDTGLNGAILQIISHGFIGAALFFLAGTSYDRIRLVYLDQMGGVAIPMP

KIFTMFSSFSMASLALPGMSGFVAEFS

>tr|A0A2J6KQY3|A0A2J6KQY3_LACSA Complex1_49kDa domain-containing protein OS=Lactuca sativa OX=4236 GN=LSAT_4X33860 PE=3 SV=1

MTGPATRKDLMIVNMGPHHPSMHGVLRLIVTLDGEDVIDCEPILGYLHRGMEKIAENRTI

IQYLPYVTRWDYLATMFTEAITVNAPEQLGNIQVPKRASYIRVIMLELSRIASHLLWLGP

FMADIGAQTPFFYIFRERELIYDLFEAATGMRMMHNFFRIGGIAADLPHGWIDKCLDFCD

YFLTGIAEYQKLITRNPIFLERVEGVGIIGGEEAINWGLSGPMLRASGIQWDLRKVDHYE

CYDEFDWEVQWQNEGDSLARYLVRISEMTESIKIIQQALEGIPGGPYENLEIRRFDRVKD

TVWNEFDYRFISKKPSLTFELSKQELYARFEAPKGELRIVLIRDKGVFPWRYKIRPPGFI

NLQILPQLVKRMKLADIMTILGSIDIIMGEVDR

>tr|A0A2J6M709|A0A2J6M709_LACSA Complex1_49kDa domain-containing protein OS=Lactuca sativa OX=4236 GN=LSAT_9X115440 PE=3 SV=1

MTGPATRKDLMIVNMGPHHPSMHGVLRLIVTLDGEDVIDCEPILGYLHRGMEKIAENRTI

IQYLPYVTRWDYLATMFTEAITVNAPEQLGNIQVPKRASYIRVIMLELSRIASHLLWLGP

FMADIGAQTPFFYIFRERELIYDLFEAATGMRMMHNFFRIGGIAADLPHGWIDKCLDFCD

YFLTGIAEYQKLITRNPIFLERVEGVGIIGGEEAINWGLSGPMLRASGIQWDLRKVDHYE

CYDEFDWEVQWQNEGDSLARYLVRISEMTESIKIIQQALEGIPGGPYENLEIRRFDRVKD

TIKDSDSDFTMISVMNSVNTSINEVGKCLDSQLWHACAGGMVQLPPLNSKVFYLPQGHAE

HAASGNVNFGDFSPIPPYILCQVSKVTFMADPDTDEVYAKIGLLPLLKHKY

>tr|A0A2J6KWB1|A0A2J6KWB1_LACSA Complex1_49kDa domain-containing protein OS=Lactuca sativa OX=4236 GN=LSAT_9X49240 PE=3 SV=1

MADIGAQTPFFYIFRERELIYDLFEAATGMRMMHNFFRIGGIAADLPHGWIDKCLDFCDY

FLTGIAEYQKLITRNPIFLERVEGVGIIGGEEAINWGLSGPMLRASGIQWDLRKVDHYEC

YDEFDWEVQWQNEGDSLARYLVRISEMTESIKIIQQALEGIPGGPYENLEIRRFDRVKDT

VWNEFDYRFISKKPSPTFELSKQELYARVEAPKGELGIFLIGDKGVFPWRYKIRPPGFIN

LQILPQLVKRMKLADIMTILGITIGVLVIVWLEREISAGIQQRIGPEYAGPLGILQALAD

GTKLLFKENLLPSRGDTRLFSIGPSIAVISILLSYLVIPFSYHLVLADLSIGVFLWIAIS

SIAPVGLLMSGYGSNNKYSFLGGLRAAAQSISYEIPLTLCVLSISLRVIR

>tr|A0A2J6LKC9|A0A2J6LKC9_LACSA Complex1_49kDa domain-containing protein OS=Lactuca sativa OX=4236 GN=LSAT_8X137301 PE=3 SV=1

MTGPATRKDLMIVNMGPHHPSMHGVLRLIVTLDGEDVIDCEPILGYLHRGMEKIAENRTI

IQYLPYVTRWDYLATMFTEAITVNAPEQLGNIQVPKRASYIRVIMLELSRIASHLLWLGP

FMADIVAQTPFFYIFRERELIYDLFEAATGMRMMHNFFRIGGIAADLPHGWIDKCLDFCD

YFLTGIAEYQKLITRNPIFLERVEGVGIIGGEEAIMSPLFGRTESLLEKCL

>tr|A0A2J6LE62|A0A2J6LE62_LACSA Complex1_49kDa domain-containing protein OS=Lactuca sativa OX=4236 GN=LSAT_5X821 PE=3 SV=1

MTGPATRKDLMIVNMGPHHPSMHGVLRLIVTLDGEDVIDCEPILGYLHRGMEKIAENRTI

IQYLPYVTRWDYLATMFTEAITVNAPEQLGNIQVPKRASYIRVIMLELSRIASHLLWLGP

FMADIGAQTPFFYIFRERELIYDLFEVVTEAYYTESHFLERVEGVGIIGGEEAINWGLSG

PMLRASGIQRDLRKVDHYECYDKFDWXKSNGKTKGIH

>tr|A0A2J6KYK0|A0A2J6KYK0_LACSA Complex1_49kDa domain-containing protein OS=Lactuca sativa OX=4236 GN=LSAT_5X66001 PE=3 SV=1

MNQEKPMTGPATRKDLMIVNMGPHHPSMHSVLRLIVTLDGEDVISYEPILGYIYRENRTI

IQYLPYVTRWDYLATMFTEAITVNAPEQLGNIQVPKRASYIRVIMLELSRIAAHLLWLGL

FMTDIGAQTPFFYIFRERELIYDLFEAATGIAEYQKLITRNPIFLERVEGVGIIGGEEAL

NWGLSGPMLQASGIQ

>tr|A0A2J6JX31|A0A2J6JX31_LACSA Complex1_49kDa domain-containing protein OS=Lactuca sativa OX=4236 GN=LSAT_4X118061 PE=3 SV=1

MHGVLRLIVTLDGEDVIDCEPILGYLHRGMEKIAENRTIIQYLPYVTRWDYLATMFTEAI

TVNAPEQLGNIQVPKRASYIRVIMLELSRIASHLLWLGPFMADIGAQTPFFYIFRERELI

YDLFEAATSMRMMHIFFRIGGIAADLPHDHLDNISTS

>tr|A0A2J6KLJ7|A0A2J6KLJ7_LACSA NAD(P)H dehydrogenase subunit H OS=Lactuca sativa OX=4236 GN=LSAT_5X124461 PE=3 SV=1

MTGPATRKDLMIVNMGPHHPSMHGVLRLIVTLDGEDVIDCEPILGYLHRGMEKIAENRTI

IQYLPYVTRWDYLATMFTEAITVNAPEQLGNIQVPKRASYIRVIMLELPCSLEXNHSLEA

HPLHHKRSACPEMTWTNREIPIHWAFRYNQIESPKGAIF

>tr|A0A2J6M2I5|A0A2J6M2I5_LACSA NAD(P)H dehydrogenase subunit H OS=Lactuca sativa OX=4236 GN=LSAT_6X87021 PE=3 SV=1

MIGPATRKDLMIVNMGPHHPSMHGVLRLIVTLDGEDVIECEPILGYLHRGMEKIAENRTI

IQYLPYVTRWDYLATMFIEAITINAPEQLGNIQVPKRANYIRVIMLELSRIASLQETIST

SATCILSIVIVEQKMEVVRLKEEMSRDLVLSRVESLALHRCIEHTEKKLTSLTLLVIGLV

IVSFCFMVEKVLHIVNK

>tr|A0A2J6KYJ4|A0A2J6KYJ4_LACSA Complex1_49kDa domain-containing protein OS=Lactuca sativa OX=4236 GN=LSAT_5X65981 PE=3 SV=1

MTESIKIIQQALEGILGGPYENLEICRFDRVKDIFNYQFISKKPSATFELSKQELYARVE

APKGELGIFLIGDKGVFPWRYKIHPPGFINLQILPKLVKIMKFTDIMTILGSIDIIMGEV

DLQAINSFSILETLKEVYGIIWMLIPIFTLLLGITIGVLVIVWLEREISAGIQQRIGPEY

ARPLGTLQALADGTKLLFKENLLPSRGDTRLFSNGPSIAVISILLSYLLSPCTSRSQYRI

APVGLLMSGYGSNNKYSFLGGLRVAAQSISYEIPLTLCVLSISLRVIC

>tr|A0A2J6JLZ5|A0A2J6JLZ5_LACSA Complex1_49kDa domain-containing protein OS=Lactuca sativa OX=4236 GN=LSAT_7X67821 PE=3 SV=1

MAQEHAHSSSVERLLNCEVPLQAQYIRVLFREITRILNHSLALTTHAMDVGASTPFLWAF

EEREKLLEFYERVSGARMHASFIQPGGVAQDLPLGLCRDIDSSTQQFASRIDELEEMSTG

NRIWKQRLVDIGTVTAQQAKDWGFSGVMLRGRAT

>tr|A0A2J6L4P7|A0A2J6L4P7_LACSA NADH-plastoquinone oxidoreductase subunit 1 OS=Lactuca sativa OX=4236 GN=LSAT_1X58461 PE=3 SV=1

MIIDTTEVQAINSFSILESLKEVYGIIWMLIPIFTLVLGITIGVLVIVWLEREISAGIQQ

RIGPEYAGPLGILQALADGTKLLFKENLLPSRGDTRLFSIGPSIAVISILLSYLVIPFSY

HLVLADLSIGVFLWIAISSIAPVGLLMSGYGSNNKYSFLGGLRAAAQSISYEIPLTLCVL

SISLRVIR

>tr|A0A2J6JX53|A0A2J6JX53_LACSA MFS domain-containing protein OS=Lactuca sativa OX=4236 GN=LSAT_4X118121 PE=3 SV=1

MKISCIVLEYGWERSSTVDIVEAQSKYGFWGWNLWRQPIGFLVFLISSLAECERLPFDLP

EAEEELVVGYQTKYSVLYFGGWNLSIPYIFVPEVFEITKRGRVFGTIIGIFITLAKTYLF

LFIPIATRWTLPRLRMDQLLNLGWKFLLPISLDIHDMFPMVTEFMNYGQQIVRVARYIG

>tr|A0A2J6LM95|A0A2J6LM95_LACSA NADH-quinone oxidoreductase subunit H OS=Lactuca sativa OX=4236 GN=LSAT_8X62921 PE=3 SV=1

MLWNYGKLNVILQIYSYSILKKVLNRIRDNEHCYMQCILMYGSGIGMGIVMLSSTILSNN

SSTVDIVEAQSKYGFRGWNLWRQPIGFLVFLISSLAECERLPFDLPEAEEELVAGYQTEY

SDINFGLFYVDSFLNLLVSSLFVTVLYLGGWNRKQGYDIEFG

>tr|A0A2J6LKZ1|A0A2J6LKZ1_LACSA NADH-ubiquinone oxidoreductase chain 1 OS=Lactuca sativa OX=4236 GN=LSAT_8X103040 PE=3 SV=1

MLTTILTLVLAIKIGALVIIWLEREIFAGIQQCIGPEYAGPLGILQALADGTKLLFKENL

LQSRGDTHLFSIGPSITFICILLSYHLVLANLSSGLLRQMVNPEGPLVIATNMENNIVTV

SNF

>tr|A0A2J6M8M6|A0A2J6M8M6_LACSA NADH-ubiquinone oxidoreductase chain 1 OS=Lactuca sativa OX=4236 GN=LSAT_5X174020 PE=3 SV=1

MDGSLDLFVTFEEPYEMKISFKIWFLGMKFVASTYRLSSYQTEYSGIKFGLFFVVSYLNL

LVSSLFITVLYLGGWNLFIPYIFVLEVFEITKRGRVFGTIIWYLYYIS

>tr|A0A2J6JX55|A0A2J6JX55_LACSA 4Fe-4S ferredoxin-type domain-containing protein OS=Lactuca sativa OX=4236 GN=LSAT_4X118101 PE=4 SV=1

MITLSHANRLPVTIQYPYEKLITSELCVRVCPIDLPVVDWKLETDIRKKRLLNYSIDFGI

CIFCGNCVEYCPTNCLSMTEEYELSTYDRHELNYNQIALGRLPMSVIDDYTIRTIFNLPE

IKT

>tr|A0A2J6LY16|A0A2J6LY16_LACSA Transmembrane protein OS=Lactuca sativa OX=4236 GN=LSAT_6X54121 PE=4 SV=1

MDLPGLIHDFLLVFLGLGLILGGLGVVLLPNPIYSAFSLGLVFVCISLSNIYQTPICSSC

ATNYSHRIHKCFNNFCCDVHKWFRIFKRFPSLDRWRWNNFGGMYKSVCFTNYYYSRYVVV

QNNLDSKSKPKYRTRFDNWLMMLEHVLVLSANLFSICLYGLIAS

>tr|A0A2J6LXX3|A0A2J6LXX3_LACSA NADH-plastoquinone oxidoreductase subunit 4L OS=Lactuca sativa OX=4236 GN=LSAT_6X54101 PE=4 SV=1

MVRALICLELILNAVNLNFVTFFCQLKGAIFSVFVIAITTAEVASGLAFVSSIYRNRKST

LINQLNLLNKLKTSHVQKWIDPMSHSIKIYDTCIGCTLMCPSPSHGCIRNDTLGRM

>tr|A0A2J6L457|A0A2J6L457_LACSA Photosystem I iron-sulfur center OS=Lactuca sativa OX=4236 GN=LSAT_1X14180 PE=3 SV=1

MKEGCMSLFKDPEPQDDKESENKYEKEPMLSRQLKGAIFSIFVIAIAAAEAAIGLAIVSS

IYRNRKSTRINQSNLLNKLKPSHVQKWIDPMSHSVKIYDTCIGCTQCVRACPTDVLEMIP

WDGCKAKQIASAPRTEDCVGCKRCESACPTDFLSVRVYLWHETTRSMGIAY

>tr|A0A2J6JK70|A0A2J6JK70_LACSA DNA-directed RNA polymerase OS=Lactuca sativa OX=4236 GN=LSAT_0X41600 PE=3 SV=1

MKEQKWIHEGLITESLPNGMFRVRLDNEDMILGYVSGKIRRSFIRILPGDRVKIERSYNL

STPKQRFKRLGVGNMKRRASVRKICEKCRLICPIPKKGSRGHIGSRKSTRKIPKGVIHIQ

ASFNNTIVTVTNVRGRVVSWSSPGTSGFRGTKRGTPFAAQTAAGHAIRAVVDQGMQRAEV

MIKGPGLGRDAALRAIRRSGILLTFVRDVTPMPHNGCRPPKKRLSTRTLQWKCVESAADS

KRLLYGRFILSPLMKGQADTIGIAMRRALLGEIEGTCITRAKSEKISHEYSTIMGIQESV

HEILMNLKEIVLRSNLYGTCEASICVRGPGYVTAQDIILPPYVEILDNTQHIASLTEPIE

LVIGLQIEKNRGYLIKAPNTFQDGSYPIDPVFMPVRNANHSIHSYENGNKEILFLEIWTN

GSLTPKEALYEASRNLIDLLIPFLHTKEENLNLEGNQHMVPLPPFTFYDKLAKLTKNKKK

MALKSIFIDQSELPPRIYNCLKRSNIYTLLDLLNNSQEDXDEEKREFLVQFSTLTTEKRI

DQILLSLTHSDHFSKNDSSYQMIEQPGAIYLRYLVDIHKPLLKYE

>tr|A0A2J6LPV9|A0A2J6LPV9_LACSA DNA-directed RNA polymerase OS=Lactuca sativa OX=4236 GN=LSAT_5X50401 PE=3 SV=1

MRRALLGEIEGTSITRAKSEKISHEYSTIMDIQESVHEILMNLKEIVLRSNLYGTCEASI

CVRGPGYVTAQDIILPPYVEILDNTQHIASLTEPIELVIGLQIEKNRRYLIKAPNTFQDG

SYPINPVFMPIRNANHSIHFYENGNKEILFLEIWKNGSFTPKEALYEASRNLIDLLIPFL

HTKEENLNLEGNQHMVPLPPFTFYDKLAKLTKNKKKMALKSIFIDQSELPPRIYNCLKRS

NIYTLLDLLNNSQEDLMKMEHFRIEDVKQILGILEKNFVIDLPKNNPKIGFESLGQFGIG

IDS

>tr|A0A2J6KRU4|A0A2J6KRU4_LACSA DNA-directed RNA polymerase OS=Lactuca sativa OX=4236 GN=LSAT_2X92080 PE=3 SV=1

MAKAIPKKGSRGRIGSRKSTRKIPKGVIHIQASFNNTIVTVTDVRGRVVSWSSAGTSGFR

GTKRGTPFAAQTAAGHAIRAVVDQGMQRAEVMIKGPGLGRDAALRAIRRSVSTRTLQWKC

VESAADSKRLLYGRFILSPLMKGQADTIGIAMRRALLGEIEGTCITRAKSEKISHEYSTI

MGIQESVHEILMNLKEIVLRSNLYGTCEASICVRGPGYVTAQDIILPPYVEILDNTQHIA

SLTEPIELVIGLQIEKNCGYLIKASNNFQDGSYPIDPVFMLVQNANHSIHSYENGNKEIL

FLEIWTNRSLTPKEALYET

>tr|A0A2J6MEG1|A0A2J6MEG1_LACSA CYTOSOL_AP domain-containing protein OS=Lactuca sativa OX=4236 GN=LSAT_8X161300 PE=4 SV=1

MSGSSGGWIYKNSPIPITKKPDLNDPVLRAKLAKGTIACNVGLAVLEPSMIGELVDPFAT

PLEILPEWYFFPRCKWNGIEKTVLET

>tr|A0A2J6K7A7|A0A2J6K7A7_LACSA PSII-H OS=Lactuca sativa OX=4236 GN=LSAT_5X67060 PE=3 SV=1

MATQIVENGAISGPRRTTVGDLLKPLNSEYGKVAPGWGTTPLMGVAMALFAVFLSIILEI

YNSSVLLDGISMN

>tr|A0A2J6LPU9|A0A2J6LPU9_LACSA Photosystem II subunit H OS=Lactuca sativa OX=4236 GN=LSAT_5X50340 PE=4 SV=1

MNTIRFMATQTVENGARSGPRRTTVGDLLKPLNSENGKVVHGWGTTPLMGVAMALFARMG

LSS

>tr|A0A2J6L6B7|A0A2J6L6B7_LACSA ATP-dependent Clp protease proteolytic subunit OS=Lactuca sativa OX=4236 GN=LSAT_4X11460 PE=3 SV=1

MHQKTPVRFRRDLTLINRLYRERLLFLGQEVDSEISNQLIGLMIYLSIEDDTKDLYLFIN

SPGGWVIPGVALYDTMQFVQPDFLTLINRLYREILLFLGQEVDSEISNQLIGLMIYLSIE

DDTKDLYLFINSPGGWVIPGVALYDTMQFVQPDVHTICMGSAASMGSFILVGGEITKLAT

GEFILEVGELLKLRETLTRVYVQRTGKPLWVVSEDMERDVFMSATEAQAYGIVDLVAVE

>tr|A0A2J6LPT8|A0A2J6LPT8_LACSA ATP-dependent Clp protease proteolytic subunit OS=Lactuca sativa OX=4236 GN=LSAT_5X50301 PE=3 SV=1

MPVRFRRGQEVDSEISNQRIGLMIYLSIEDDTKDLYLFINSPGGWVIPGVALYDTMQFVQ

PDVHTICMGSAASMGSFILARVNFIVEPVMIHQPAGSFSEVATGEFIMEVGKLLKLHESL

TRVYVQRTGKTLWVVSEDMEIDVFMSAT

>tr|A0A2J6KPB2|A0A2J6KPB2_LACSA ATP-dependent Clp protease proteolytic subunit OS=Lactuca sativa OX=4236 GN=LSAT_3X3401 PE=3 SV=1

MAVSLNSITCSSYLHPPQPSLSFGDKVFVGLRLQSPNSYGISRPNLSAELHKKIHKSIEP

RIGIKPARGRITMMPIGTPRVPYRVPGEGTWQWVDLWNALYRERVIFIGQNIDEEFSNQI

LATMLYLDSVDNSKRMYMYINGPGGDLTPSMAIYDTMQSLQSPIGTHCVGYAYNLAGFLL

AAGEKGQRFAMPLSRIALESPAGAARGQADDIQNEANELLRIRDYLFKELAQKTGQPVEQ

VHKDLSRIKRFNAQEALDYGLIDRIVRPPRIKADAPRQEAGTGLGVDIKQTDCFIHLRIC

CDYT

>tr|A0A2J6M1D3|A0A2J6M1D3_LACSA ATP-dependent Clp protease proteolytic subunit OS=Lactuca sativa OX=4236 GN=LSAT_9X50981 PE=3 SV=1

MCPYLFIIKGHYSVMIHQPAGSFSEVATGEFILEVGELLKLRETLTRVYVQRTGKPLWVV

SEDMERDVFMSATEAQAYGIVDLVAVE

>tr|A0A2J6JUF6|A0A2J6JUF6_LACSA Cytochrome b559 subunit alpha OS=Lactuca sativa OX=4236 GN=LSAT_1X27981 PE=3 SV=1

MSGSTGERSFADIITSIRYWVIHSITIPSLFIAGWLFVSTGLAYDVFGSPRPNEYFTENR

QGIPLITGRFDSLEQLDEFSRSF

>tr|A0A2J6JPQ0|A0A2J6JPQ0_LACSA Cytochrome b559 subunit alpha OS=Lactuca sativa OX=4236 GN=LSAT_2X43901 PE=3 SV=1

MSGSTGERSFADIITSIRYWVIHSITIPSLFIAGWLFVSTGLAYDVFGSPRPNEYFTENR

QGIPLITGRFDSLEQLDEFSRSFLEASMTIDRTYPIFTVRWLTVSCTYXLLILLPVFATG

SFIALLYLPYSLRVGYSSALV
