## Supplementary material for "Sustained non-photochemical quenching and regulation of PSII repair cycle during combined low temperature and high light stress in lettuce": Lettuca and Arabidopsis LHC and LIL aligments.pptx

### Slide 1
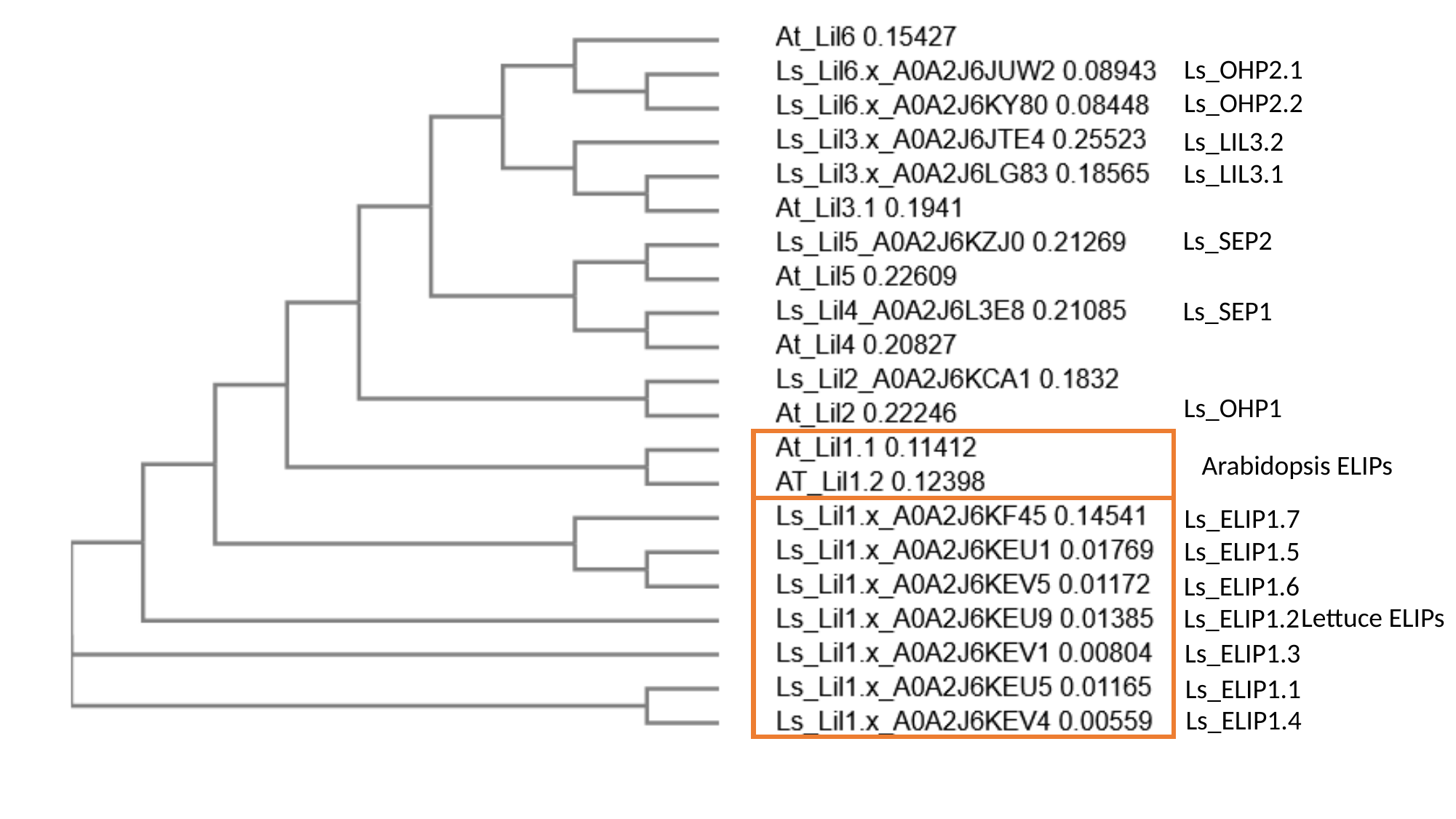

Ls_OHP2.1
Ls_OHP2.2
Ls_LIL3.2
Ls_LIL3.1
Ls_SEP2
Ls_SEP1
Ls_OHP1
Arabidopsis ELIPs
Ls_ELIP1.7
Ls_ELIP1.5
Ls_ELIP1.6
Lettuce ELIPs
Ls_ELIP1.2
Ls_ELIP1.3
Ls_ELIP1.1
Ls_ELIP1.4

### Slide 2
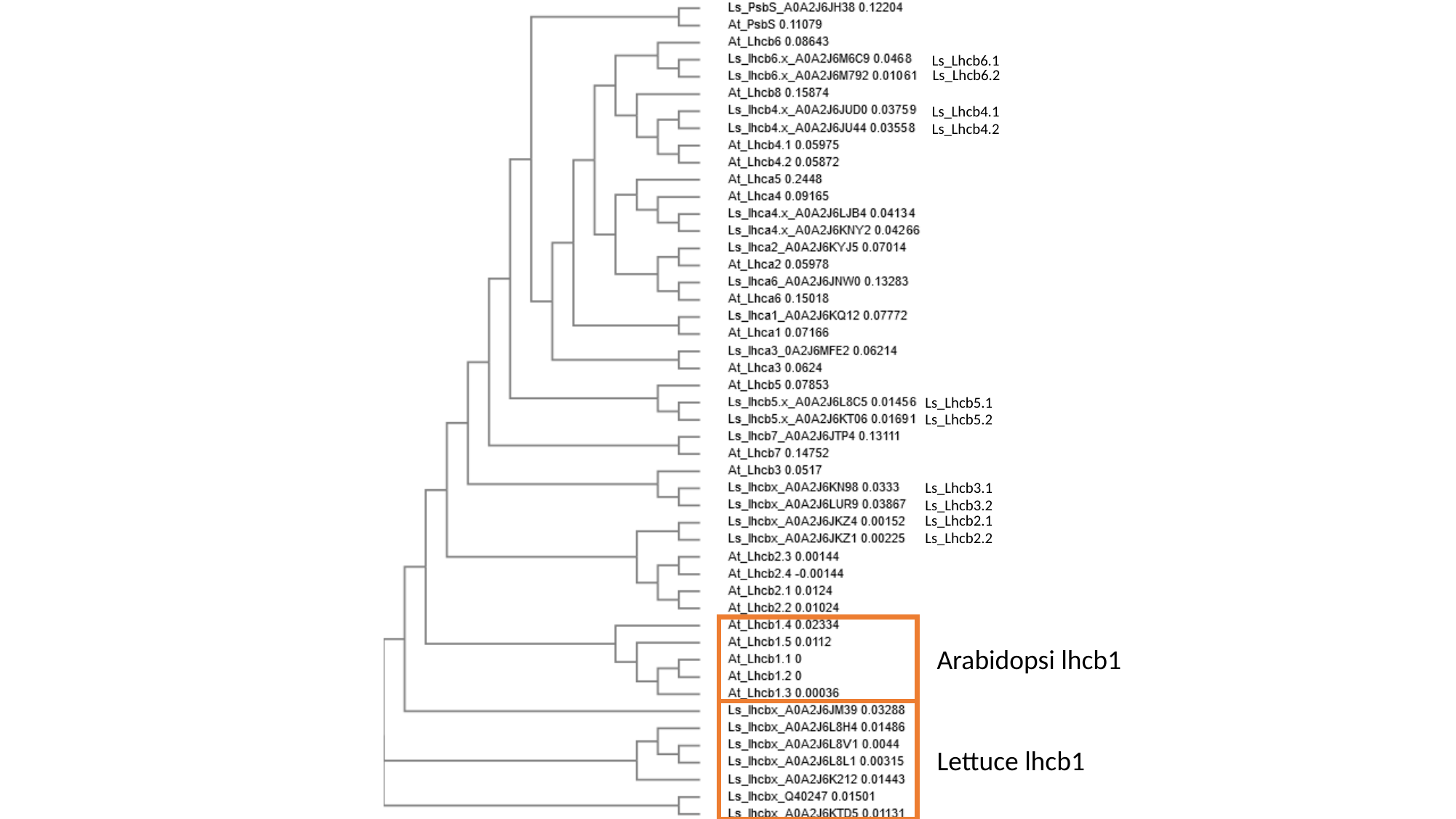

Ls_Lhcb6.1
Ls_Lhcb6.2
Ls_Lhcb4.1
Ls_Lhcb4.2
Ls_Lhcb5.1
Ls_Lhcb5.2
Ls_Lhcb3.1
Ls_Lhcb3.2
Ls_Lhcb2.1
Ls_Lhcb2.2
Arabidopsi lhcb1
Lettuce lhcb1
