## Supplementary figures and images for "Sustained non-photochemical quenching and regulation of PSII repair cycle during combined low temperature and high light stress in lettuce"

### Lettuca and Arabidopsis LHC and LIL aligments.pdf

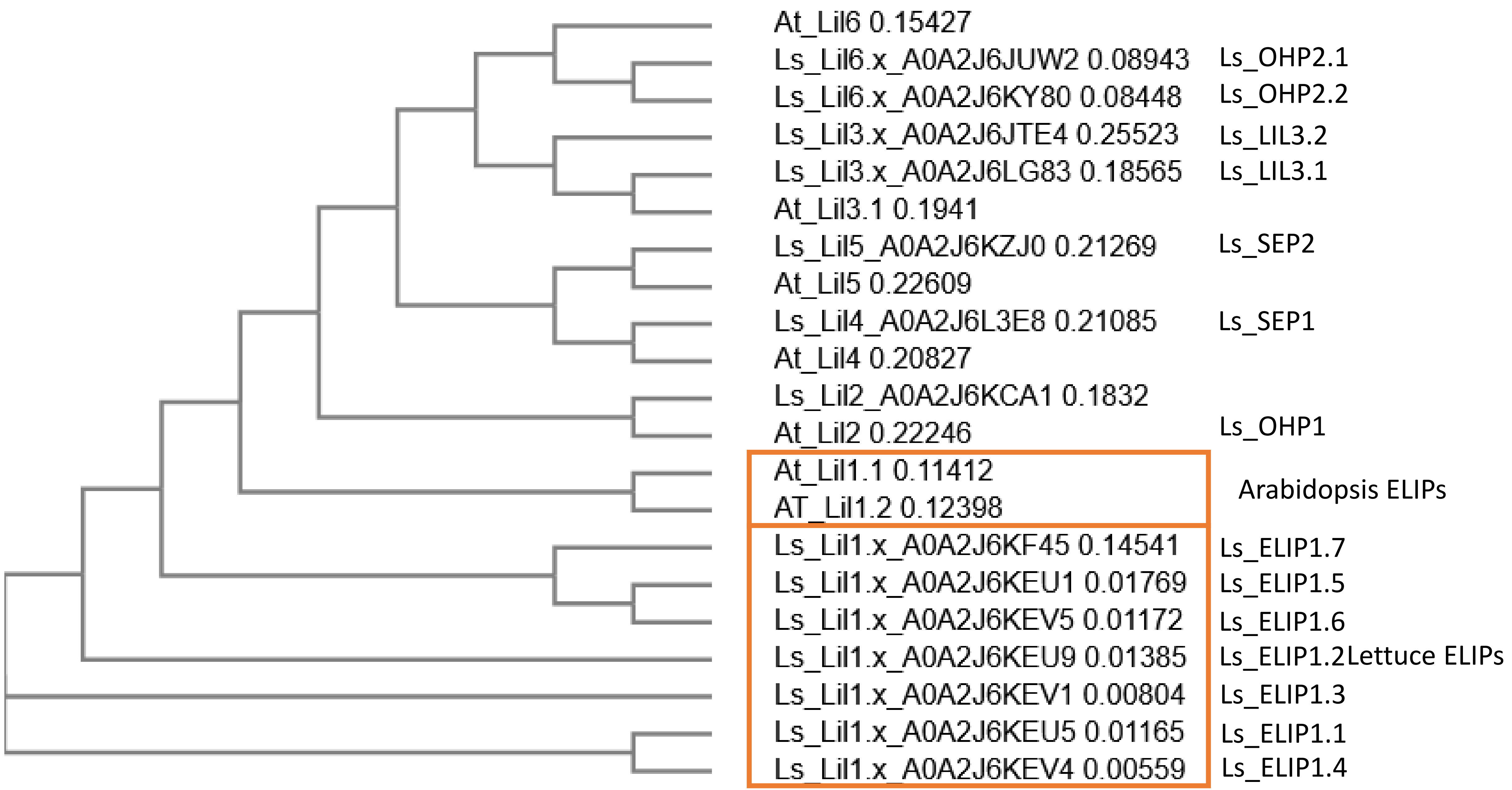

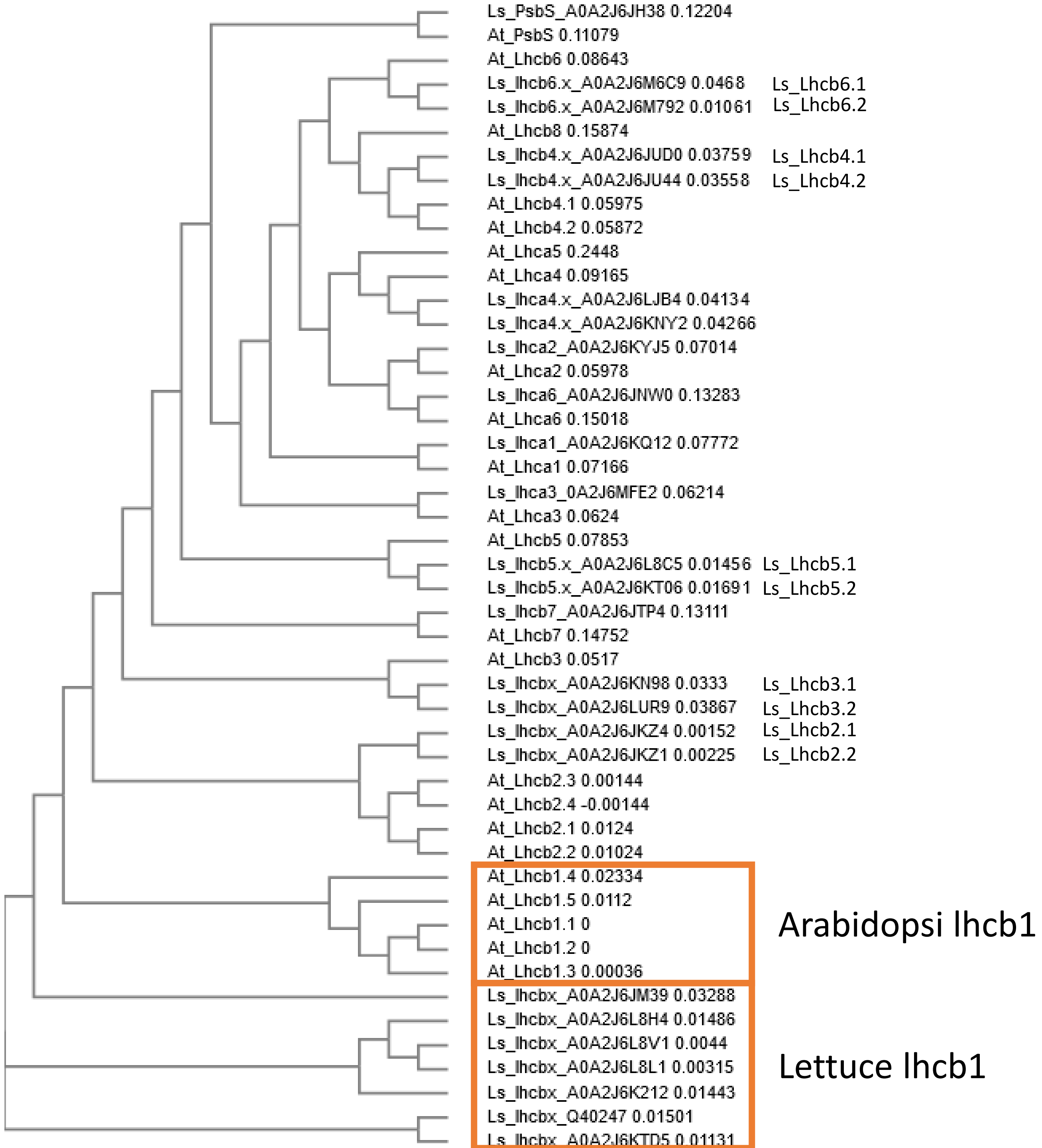
